# Post-Transcriptional Modular Synthetic Receptors

**DOI:** 10.1101/2024.05.03.592453

**Authors:** Xiaowei Zhang, Luis S. Mille-Fragoso, K. Eerik Kaseniit, Arden P. Lee, Meng Zhang, Connor C. Call, Yixin Hu, Yunxin Xie, Xiaojing J. Gao

**Affiliations:** Department of Bioengineering, Stanford University; Stanford, 94305, USA; Department of Chemical Engineering, Stanford University; Stanford, 94305, USA; Department of Biology, Stanford University; Stanford, 94305, USA; Sarafan ChEM-H, Stanford University; Stanford, 94305, USA; Stanford Bio-X, Stanford University; Stanford, 94305, USA; The Chinese Undergraduate Visiting Research (UGVR) Program; Stanford, 94305, USA

## Abstract

Inspired by the power of transcriptional synthetic receptors and hoping to complement them to expand the toolbox for cell engineering, we establish LIDAR (Ligand-Induced Dimerization Activating RNA editing), a modular post-transcriptional synthetic receptor platform that harnesses RNA editing by ADAR. LIDAR is compatible with various receptor architectures in different cellular contexts, and enables the sensing of diverse ligands and the production of functional outputs. Furthermore, LIDAR can sense orthogonal signals in the same cell and produce synthetic spatial patterns, potentially enabling the programming of complex multicellular behaviors. Finally, LIDAR is compatible with compact encoding and can be delivered as synthetic mRNA. Thus, LIDAR expands the family of synthetic receptors, holding the promise to empower basic research and therapeutic applications.

## Introduction

Cells in multicellular organisms modulate their behaviors in response to cues present in their environment or sent by other cells using endogenous receptors. Therefore, to study and engineer mammalian cells, it is critical to possess the ability to understand and further manipulate such interactions. This capability is endowed by the development of synthetic receptors, engineered membrane proteins that can be programmed to sense specific inputs and produce customized outputs ^1^. Remarkably, synthetic receptors have empowered basic research such as investigating the spatiotemporal dynamics of signaling events ^2,3^ and neuronal connections ^4,5^, as well as biomedical applications including but not limited to screening small molecule for drug development ^6,7^, directing therapeutic cells to ablate anomalous cells ^8,9^ and building complex multicellular structures for potential tissue engineering ^10,11^.

Despite the advances of synthetic receptors, all modular ones to date rely on transcriptional actuation. Here we postulate that to expand the applications of synthetic receptors, these could be complemented by using post-transcriptional activation mechanisms, producing synthetic receptors that can be delivered using either DNA or, uniquely, RNA vectors. For the former, post-transcriptional receptors would synergize with DNA-level controls to improve specificity (e.g., producing a payload based on a tissue-specific enhancer AND the response to a ligand). For the latter, taking advantage of the developments spurred by mRNA vaccines, such receptors could avoid the risk of insertional mutagenesis ^12^ and enable dynamic dosing. In contrast to DNA vectors that require nuclear entry, the cytosolic translation of mRNA vectors might also improve the speed, efficiency, and consistency of receptor expression – relevant for cases of transient delivery ^13,14^. In summary, there remains an opportunity for developing modular post-transcriptional synthetic receptors that can expand and complement applications of synthetic receptors.

To design post-transcriptionally actuated receptors, we surveyed different mechanisms. Many inspiring examples of post-transcriptional regulators emerged in the past decade ^15–18^, yet they rely on endogenous biomolecular complexes, such as the translation initiation complex ^15,17^ or the exon junction complex that controls mRNA stability ^18^. We postulated that the engagement with these complexes subjects the actuation mechanisms to intracellular regulations and that their performance might be sensitive to cellular context ^19,20^, making them difficult to engineer or optimize. Alternatively, we identified RNA editing as a potentially robust post-transcriptional actuation mechanism, as it only requires standalone enzymatic reactions, and small-molecule control of such editing has been previously demonstrated ^21,22^. We decided to harness adenosine deaminases acting on RNA (ADAR), an enzyme that edits adenosine (A) to inosine (I), treated as guanosine (G) in translation. Each ADAR consists of a catalytic deaminase domain (e.g., ADAR2dd) and double-strand RNA binding domains (dsRBDs) that recruit it to editing sites ^23^. We propose the design of LIDAR (<u>L</u>igand-Induced <u>D</u>imerization <u>A</u>ctivating <u>R</u>NA editing), by replacing the dsRBDs on ADAR2 with domains that recruit ADAR2dd to a synthetic RNA substrate in the presence of a specific ligand (**Fig. 1A**). More specifically, LIDAR consists of three components: two customizable dimerization domains – one fused to an ADAR2dd and the other to a specific RNA binding protein (RBP) – and a reporter RNA molecule harboring multiple RBP recruiting sequences and an in-frame stop codon (UAG) within a double-stranded region as an ADAR substrate. When the reporter RNA is brought into the proximity of ADAR2dd, the stop codon gets edited into a permissive tryptophan codon (UGG), resulting in the translation of the downstream payload.

**Fig. 1.**
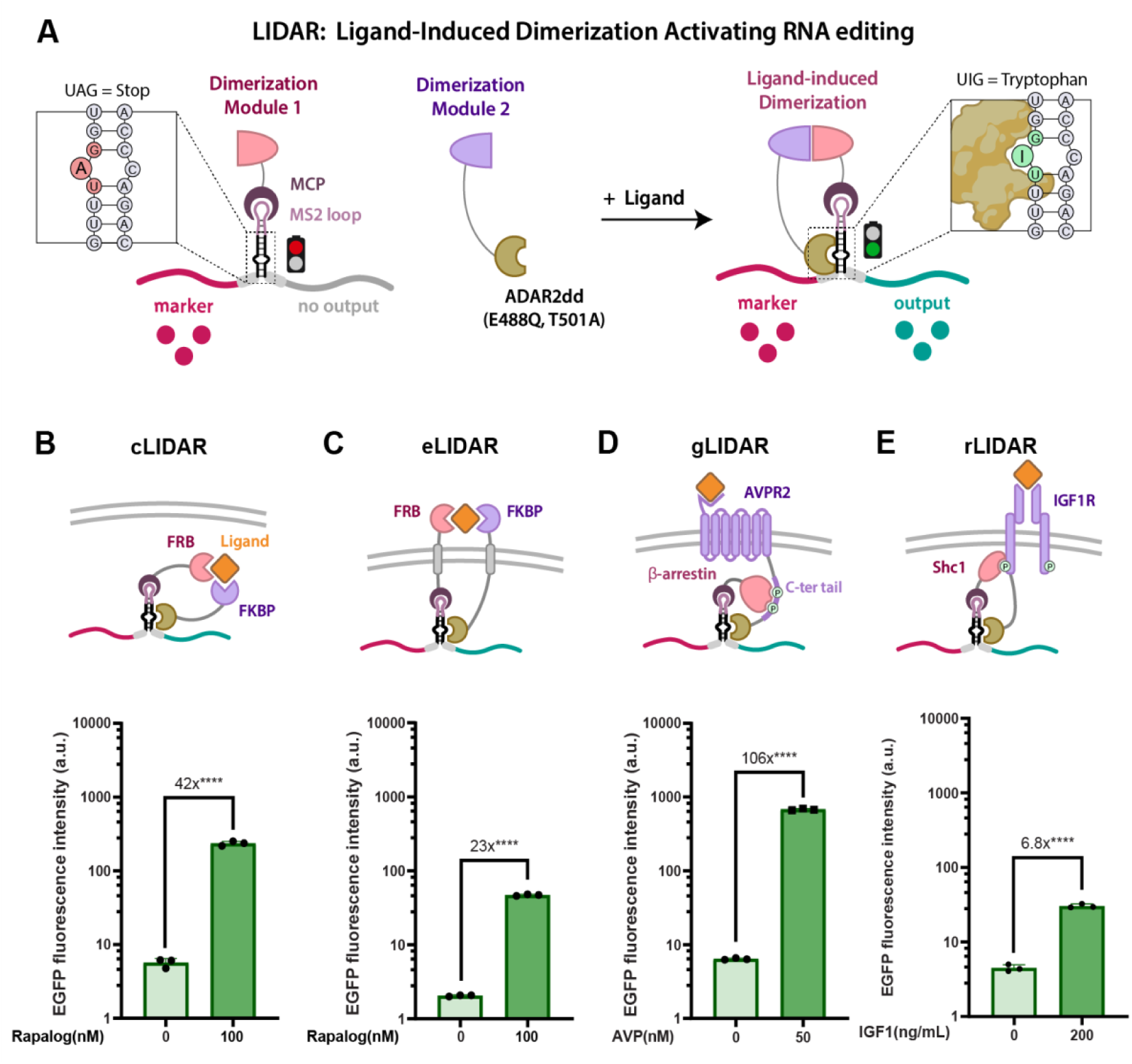
Design of a modular post-transcriptional synthetic receptor architecture. (**A**) The general LIDAR (<u>L</u>igand-Induced <u>D</u>imerization <u>A</u>ctivating <u>R</u>NA editing) architecture consists of three main components. The first component incorporates a dimerization module fused to an RNA Binding Protein (RBP, i.e., MCP). The second component contains the other dimerization module fused to an ADAR2dd which can edit dsRNAs when in close proximity. The third component is a reporter RNA harboring an in-frame stop codon within a double-stranded stem loop that functions as an ADAR substrate, and multiple RNA stem loops that recruit specific RBPs (e.g., MS2 that recruits MCP). The stem-loop is placed in between the marker and the output, flanked by F2A self-cleavage peptide, preventing translation of the output. The presence of ligand brings the two dimerization modules together such that ADAR2dd comes in proximity to the reporter RNA, triggering editing of the UAG stop codon into a UIG tryptophan codon, leading to payload translation. (**B**) The LIDAR architecture is used to implement a cytosolic binder-based receptor (cLIDAR), (**C**) an extracellular binder-based receptor (eLIDAR), (**D**) a GPCR-based receptor (gLIDAR), and (**E**) an RTK-based receptor (rLIDAR). The data for each of the implementations represents fluorescent intensities and activation fold-change for induced vs. uninduced conditions. Refer to **Methods** section for statistical analysis.

Here we demonstrate that LIDAR is compatible with a variety of receptor architectures. We show that it is capable of sensing diverse ligands and producing different functional outputs, and is compatible with various cellular contexts. Furthermore, we show that it has the potential to enable the engineering of complex cellular behaviors, such as processing orthogonal inputs in the same cell and spatial patterning based on intercellular communications. Finally, we show that LIDAR is compatible with compact, single-transcript encoding and, critically, it is indeed compatible with RNA delivery. We expect to further develop this versatile LIDAR platform for post-transcriptional sense-and-response capabilities, expanding the family of synthetic receptors for basic research and therapeutic applications.

### Design and optimization of LIDAR

To test the feasibility of LIDAR, we started with a system containing FRB and FKBP, two domains that dimerize in response to rapalog (AP21967) ^24^, to create a cytosolic binder-based LIDAR (cLIDAR). (**Fig. 1B**). To facilitate quantification and iterative engineering, we used a reporter RNA encoding mCherry upstream of the stop codon as the marker and GFP downstream of the stop codon as the output. The stop codon is located in a short dsRNA sequence modified from the endogenous GluR-B R/G site, a canonical ADAR2 substrate ^25^ (**Supplementary Fig. 1A**), and the reporter RNA includes several MS2 stem-loops to recruit the corresponding phage-derived RBP, MS2 Coat Protein (MCP) ^26^. We then performed optimizations on this initial design. We added the self-cleaving F2A peptides ^27^ to insulate the marker and the output proteins from the non-functional peptide encoded by the ADAR substrate region. We tested different numbers of tandem MS2 stem-loops (**Supplementary Fig. 1B-C**) ^28^ and concluded that the output benefits from an increase in MS2 stem-loop repeats. As for ADAR2dd, we hypothesized that the optimal design would possess high catalytic activity, while maintaining low substrate binding affinity to minimize ligand-independent recruitment of the substrate. This motivated us to test two ADAR2dd point mutations: the “hyperactive” mutation E488Q with enhanced catalytic activity, and the “de-dimerization” mutation T501A which might reduce ligand-independent baseline (because dimerized ADAR increase substrate affinity) ^29–31^, and validated their benefits (**Supplementary Fig. 1D-E**). As a result, our optimal design contains 3 tandem MS2 stem-loops connected to the GluR-B-derived ADAR substrate and adopts both E488Q and T501A point mutations in ADAR2dd, exhibiting minimal leaky baseline and substantial responsiveness to rapalog (**Fig. 1B**).

We then progressed to sense extracellular ligands, while examining the compatibility of LIDAR with a variety of receptor architectures. First, ligand-induced dimerization could be mediated by directly binding to the same ligand in cLIDAR but on the extracellular side. Inspired by MESA ^32^, we added a CD28 transmembrane domain between each of the dimerization and actuation modules of cLIDAR (**Fig. 1C**). We validated the response of such an extracellular binder-based LIDAR (eLIDAR) to rapalog (**Fig. 1C**). Next, ligand-induced dimerization could also be facilitated indirectly by downstream protein recruitment upon natural receptor activation. Inspired by TANGO ^33^, we harnessed the recruitment of β-arrestin2 (ARRB2) to the phosphorylated C-terminal tail of a G protein-coupled receptor (GPCR) to create a GPCR-based LIDAR (gLIDAR). Using AVPR2, a GPCR for vasopressin (AVP) previously used in TANGO, we validated an AVP-responsive LIDAR by fusing ADAR2dd to the C-terminal tail of AVPR and MCP to the C-terminus of ARRB2 (**Fig. 1D; Supplementary Fig. 1G**). After linker optimization, we determined that short 4 amino acid linkers between LIDAR binding domains and actuation domains results in a low baseline (**Supplementary Fig. 1F**). We quantified the temporal dynamics of gLIDAR with a PiggyBac stably integrated cell line (**Supplementary Fig. 1H-I**) and transient transfection (**Supplementary Fig. 1K**), where the activation dynamics between transient transfection and stable integration are comparable. Through transient transfection, we also demonstrated that gLIDAR can reach activation levels comparable to those of Transcription Factor (TF)-based systems while maintaining a lower baseline (**Supplementary Fig. 1J**), and that these two systems exhibit comparable temporal responses to the ligand (**Supplementary Fig. 1K-L**). Furthermore, to generalize LIDAR to other signaling-induced dimerization events, we demonstrated that LIDAR works with IGF1R, a receptor tyrosine kinase (RTK), and its activation-dependent binding partner Shc1 (**Fig. 1E**) ^33^. Therefore, LIDAR can be implemented using a variety of architectures and can induce robust outputs upon ligand activation.

To maintain a fast design-build-test cycle – unless specified – we performed all our prototyping experiments by transiently co-transfecting receptor-encoding DNA plasmids into human embryonic kidney (HEK) 293 cells. Because LIDAR operates at the post-transcriptional level, we expect the results to reflect key features of LIDAR’s performance in other delivery scenarios such as stable integration or via mRNA.

### Compatibility with diverse inputs and cellular contexts

As a demonstration of modularity, we show that LIDAR can sense diverse inputs by employing different receptor architectures. Inspired by recent progress in NatE MESA that enables recombinant ligand sensing ^34^, we first sought to build a modular eLIDAR by leveraging natural receptor ectodomains (**Fig. 2A**) and prototyped a VEGF-sensing eLIDAR using VEGFR1 and VEGFR2 ecto-and transmembrane domains (**Fig. 2B, Supplementary Fig. 2A**). Signal activation by recombinant VEGF-165 is improved by attaching Golgi traffic sequence and ER export sequence to the receptor intracellular domains ^35^ (**Supplementary Fig. 2B**), yet likely through a mechanism distinct from improving overall cell surface expression (**Supplementary Fig. 2C-D**). Co-transfection of an ER-retained catalytic inactive ADAR2dd(E396A) further improves signal activation (**Supplementary Fig. 2E**), potentially through inhibition of ligand-independent activation due to spatial proximity in the secretory process. These optimizations resulted in an eLIDAR that, when transiently transfect, responds to both autocrine and recombinant VEGF-165 (**Fig. 2C**). When its components are stably integrated into the genome (cell line XZCP14), eLIDAR responds even more potently to recombinant VEGF-165 at a physiologically relevant level ^36^ (**Fig. 2D** and **Supplementary Fig. 2F-G**). Using similar design principles, we also built IL-2 sensing eLIDAR that responds to both autocrine and recombinant IL-2 (**Fig. 2E-G** and **Supplementary Fig. 2H-I**).

**Fig. 2.**
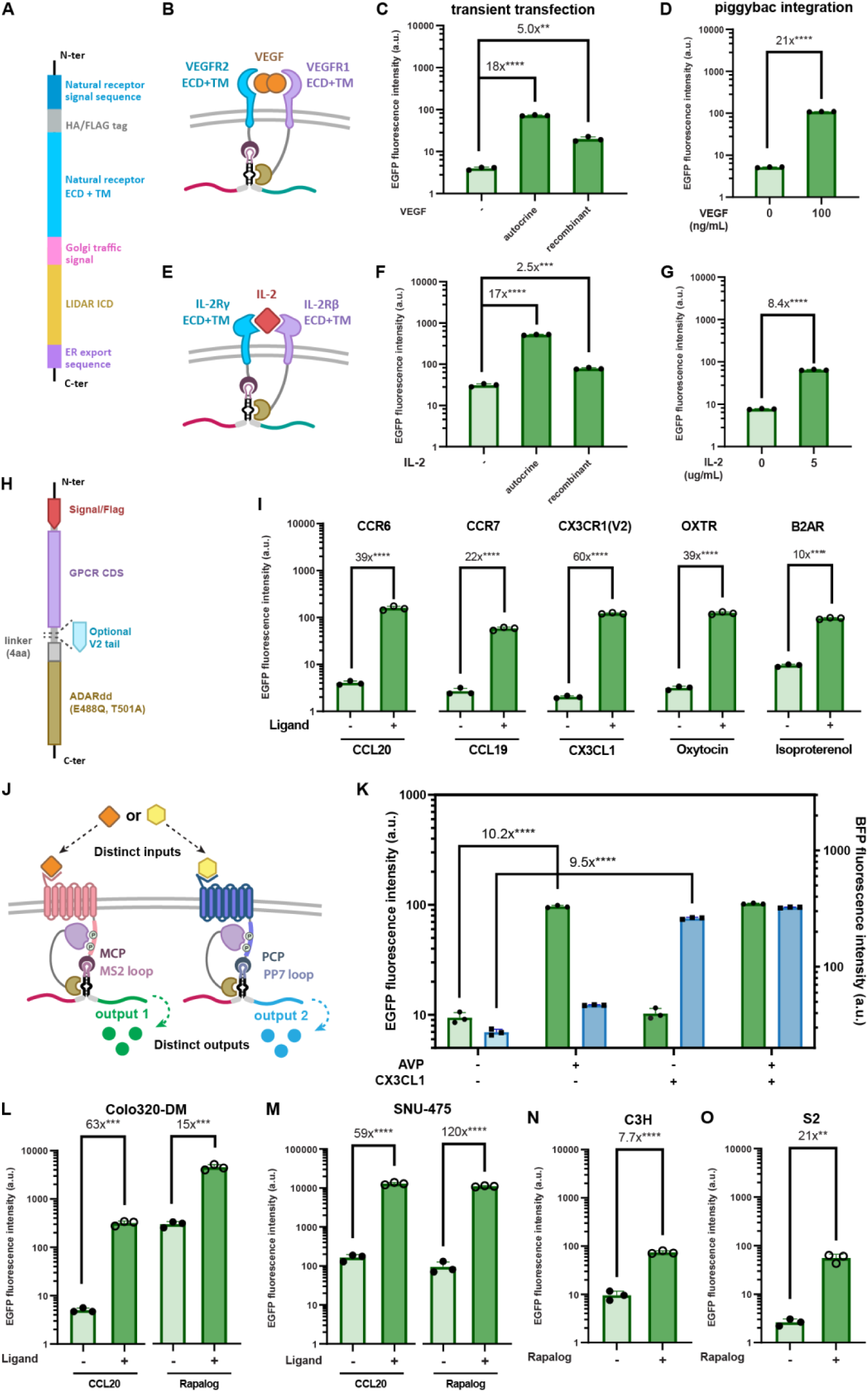
Molecular and cellular versatility of LIDAR. (**A**) Modular design principles of eLIDAR leveraging natural receptor ectodomains. The Golgi traffic signal (KSRITSEGEYIPLDQIDINV) is inserted between the natural receptor transmembrane domain and the LIDAR intracellular domains without additional GS linkers. The ER export sequence (FCYENEV) is appended to the C-terminus of the LIDAR intracellular domains. (**B**) Schematics of VEGF-sensing eLIDAR leveraging VEGFR1 and VEGFR2 ectodomains. (**C**) VEGF-sensing eLIDAR in response to autocrine and recombinant VEGF by transient transfection, co-transfected with ER-retained catalytically inactive ADARdd(E396A) to reduce baseline editing. Autocrine: co-transfected CMV-VEGF-165; Recombinant: 200 ng/mL recombinant VEGF-165. (**D**) VEGF-sensing eLIDAR in response to 100 ng/mL recombinant VEGF-165 in a cell line with all components stably integrated (XZCP14), using optimal gating determined by **Supplementary Fig. 2F**. (**E**) Schematics of IL-2-sensing eLIDAR leveraging the ectodomains of IL-2R β and γ chains. (**F**) IL-2 sensing eLIDAR in response to autocrine and recombinant IL-2 by transient transfection, co-transfected with ER-retained catalytically inactive ADARdd(E396A) to reduce baseline editing. Autocrine: co-transfected CMV-IL-2; Recombinant: 1 μg/mL recombinant IL-2. (**G**) IL-2-sensing eLIDAR in response to 1 μg/mL recombinant IL-2 in a cell line with all components stably integrated (XZCP17), using optimal gating determined by **Supplementary Fig. 2H**. (**H**) Modular design principles of gLIDAR. (**I**) Different GPCRs with LIDAR exhibit robust signal activation. Refer to **Supplementary Table 3** for ligand catalog and concentrations used. Unless specified, all ligands are applied at the same concentration as listed in Supplementary Table 3. (**J**) Orthogonal GPCR-based LIDARs (oLIDARs) can be implemented in a single cell to sense different inputs and produce separate outputs. **(K)** Both AVPR-based and CX3CR1-based oLIDARs are transfected in the same cell. The y-axes for BFP and GFP are in disparate range to portrait both baselines at similar apparent levels while still showing the outputs using their measured values and units, to be consistent with other plots in the manuscript. (**L-O**) LIDAR is robust in distinct cellular contexts. LIDAR generates strong activation signal comparable with benchmarking conditions in (**L**) Colon cancer (Colo 320DM) and (**M**) Liver cancer (SNU-475) cell lines with FKBP-FRB eLIDAR and CCR6-based gLIDAR, as well as eLIDAR in **(N)** C3H mouse embryonic fibroblast cell line and (**O**) Schneider 2 (S2) drosophila cells.

In a similar fashion, we demonstrated the modularity of gLIDAR by replacing the GPCR portion with several other GPCRs, including chemokine receptors CCR6, CCR7, CX3CR1, as well as the oxytocin receptor (OXTR) and β-2 adrenergic receptor (B2AR) (**Fig. 2H-I**). It’s worth noting that in PRESTO-TANGO ^6^, an additional V2 tail was included to enhance the signal. For gLIDAR, the additional V2 tail increases both the signal and the baseline, yet its effect on fold activation is not always beneficial (**Supplementary Fig. 3A**). For GPCRs like AVPR2 and CX3CR1 that produce a low baseline, the V2 tail improves the fold activation by increasing the activation signal more than the baseline. However, for GPCRs like CCR6 and B2AR that produce higher baselines, the V2 tail drastically reduces fold activation. Finally, similar to eLIDAR, a stably integrated CCR6 gLIDAR cell line (XZCP08) was generated and its dose responsive curve was compared to that of a transiently transfected CCR6 gLIDAR (**Supplementary Fig. 3B-C**). Both configurations share a similar dose-response curve, which gave us confidence that we could continue testing designs by transient transfection.

To ultimately program more complex multi-cellular behaviors such as cellular homeostasis ^37^ or directed differentiation ^11^, as well as to integrate multiple inputs for targeted cell therapies ^38^, it is often required to have receptors that process orthogonal signals in the same cell. As demonstrated above, gLIDAR is robust and produces strong outputs, making it an ideal candidate for biomedical applications. However, in the current configuration, the ARRB2-RBP fusion protein is recruited to all GPCRs without discrimination. With such design, if we have two gLIDAR receptors in a single cell, the system can only produce one output when triggered by either ligand. We reasoned that, analogous to TANGO vs. ChaCha receptors ^39^, an inverted version of gLIDAR (orthogonal GPCR-based LIDAR; oLIDAR) can accommodate two or more receptors in a single cell, each producing its corresponding output when activated by the respective ligand. We demonstrated that by fusing ARRB2 to ADAR2dd (ADAR2-ARRB2) and GPCR to an RBP (GPCR-RBP), we can create functional LIDAR receptors (**Supplementary Fig. 3D-E**). Furthermore, using an AVPR2-MCP oLIDAR with a reporter RNA containing MS2 stem-loops (GFP output) and a CX3CR1-PCP (PP7 bacteriophage coat protein) oLIDAR with a reporter RNA containing PP7 stem-loops (TagBFP output) ^40^, we demonstrate the orthogonal response of each oLIDAR to its respective ligand (**Fig. 2J-K**).

Next, we examined LIDAR’s performance in different cellular contexts. To explore the feasibility of cell/gene therapy applications, such as the autonomous ablation of diseased cells based on extracellular cues, and given that that CCL20 is elevated in some tumor microenvironments ^41^, we validated rapalog-and CCL20-responsive LIDAR in human colon cancer (Colo 320DM) and liver cancer (SNU-475) cell lines (**Fig. 2L-M**). As for its potential use in a variety of organisms, we validated LIDAR in mouse embryonic fibroblast (C3H) cells (**Fig. 2N**), as well as drosophila Schneider 2 (S2) cells (**Fig. 2O**).

### Further characterization of LIDAR

To better understand the factors that affect LIDAR performance, we decided to investigate the effects of component stoichiometry, as well as the design of the reporter RNA, on the output levels. For gLIDAR, the output and baseline levels are modulated by the relative amount of GPCR-ADAR2dd and ARRB2-MCP to the reporter RNA _(_**Supplementary Fig. 3F-G**). Furthermore, given the high baselines observed in some of the LIDAR experiments (e.g. B2AR gLIDAR; **Fig. 2**), we found that it’s possible to reduce baseline by including additional stop codons on the reporter RNA, though at the cost of reducing activation levels (**Supplementary Fig. 3H-I**).

We then explored the source of high baseline. Using experiments where at least one component of LIDAR was left out, we show that higher baselines are mainly caused by the presence of the component containing the ADAR2dd (**Supplementary Fig. 4A-E**). Similarly, using HEK293 cells stably integrated with full-length ADAR2, we verified that overexpression of ADAR2 substantially increases the baseline (**Supplementary Fig. 4F**). This effect, however, is not observed when transfected via mRNA (**Supplementary Fig. 10E**), as we hypothesize that the RNA reporter transcribed from plasmids are primarily edited in the nucleus by ADAR2 before traveling to the cytosol. Finally, a similar effect could potentially account for the elevated baseline in SNU-475 cells, which have been reported to have ∼3x higher expression of ADAR1 ^42^.

Moreover, we directly quantified the editing of the LIDAR reporter via amplicon NGS (see **Methods**). Using transient transfection, we saw a significant increase in editing rates with ligand treatment for three gLIDARs of different efficiencies (**Supplementary Fig. 5A**). Significant editing was also observed for cell lines that stably express VEGF-sensing eLIDAR or CCL20-sensing gLIDAR (**Supplementary Figs. 5B**). We note that, although we observed some consistency between the fluorescent outputs and editing rates, we do not expect strict correlation between them, as the former reflects the product of total reporter mRNA counts and editing rates. That could also partially account for the apparent difference in editing rates between transient and stable conditions, despite their comparable fluorescent outputs.

Next, we asked whether LIDAR produces substantial off-target editing in endogenous transcripts. By bulk transcriptomic sequencing, a stably integrated AVP-sensing gLIDAR (cLSM02) shows elevated editing at few off-target sites when compared to WT HEK293 (Nsig < 300 out of 1,868,346 sites tested; **Supplementary Fig. 6A**). Similarly, when AVP is added to this cell line, the number of significantly edited sites increased from the no-ligand condition (see **Methods**), yet the editing frequencies at most off-target sites are still below the on-target site (green arrow). To further understand the effect of overexpressing ADAR2dd in a LIDAR setting, we generated 4 stable cell lines integrated only with the ADAR2dd-containing component of LIDAR, or full-length ADAR2 (**Supplementary Fig. 6B**). When compared with WT HEK293 cells, we found that cLIDAR, eLIDAR and gLIDAR cause very few significant off-target editing events (Nsig < 22 of 3,297,939 sites tested in all three conditions), in contrast to the large number with full-length ADAR2 (Nsig=4640 of 3,297,939 sites tested; **Supplementary Fig. 6B**). These suggest that the modest off-target effects of LIDAR are of no major concern.

### Functional outputs

In addition to fluorescent signals, we validated that the output levels of LIDAR are sufficient to produce functional outcomes. We used gLIDAR to produce an iCASP9-based ligand-dependent cell death controller ^43^ (**Fig. 3A**). We chose iCASP9 as it can efficiently induce apoptosis in response to the dimerizing molecule AP1903 ^43^. Utilizing a monoclonal HEK cell line of a CCL20-sensing gLIDAR with iCASP9-EGFP as the output (cLSM01), and by counting the number of remaining live cells in a well, we observed substantial cell death only when both the ligand (CCL20) and AP1903 are present (**Fig. 3B**). Such quantification based on flow cytometry is consistent with our microscopy-based visual assessments (**Supplementary Fig. 7A**) as well as with Annexin-V staining (**Supplementary Fig. 7B**). Notably, we observed no significant killing when AP1903, but not CCL20, is present, suggesting that the low baseline of LIDAR might be adequate for the autonomous ablation paradigm. Similarly, we also did not see any significant death when iCASP9 (co-expressed with EGFP) was highly expressed but in the absence of AP1903 (**Supplementary Fig. 7C**). Additionally, given the broad use of the tobacco etch virus protease (TEVP) as a critical intermediate in protease-based circuitry^44^, we validated that LIDAR-produced TEVP can rescue a fluorescent protein linked via a TEVP cleavage site to a dihydrofolate reductase (DHFR) degron^45^ (**Fig. 3C**), using both cLIDAR and gLIDAR (**Fig. 3D-E**). The dose-response curve resembles that of the corresponding LIDAR directly outputting EGFP (**Supplementary Fig. 8A**), which indicates that the terminal output of the system is not dampened by the use of the protease. Such results depicts the potential for LIDAR to be implemented in more complex circuits ^46^.

**Fig. 3.**
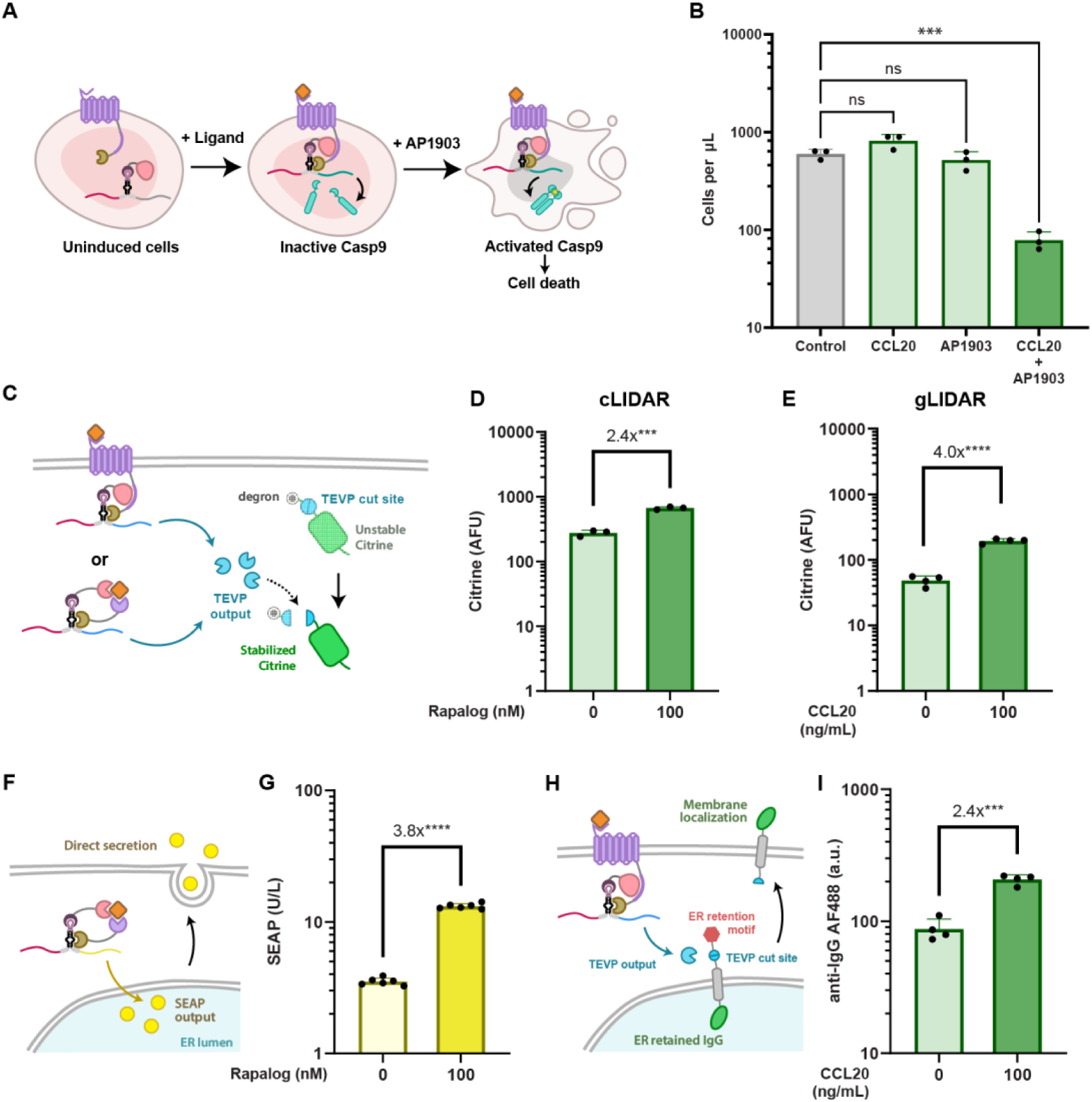
Expanding LIDAR’s output diversity. (**A**) Schematics of the capacity of LIDAR to drive cell apoptosis by producing a functional iCasp9. (**B**) Quantification of the remaining live cells after LIDAR-induced cell apoptosis. Additional supporting data and information on this assay can be found in **Supplementary Fig. 7** and the **Methods** section. (**C)** LIDAR is capable of interfacing with protease-based circuitry ^49^ through outputting TEV protease. The TEV protease output further controls the stability of a degron-tagged fluorescent protein Citrine with a flanking TEVP cut site. (**D**-**E**) Citrine AFU of (**D**) Rapalog-sensing cLIDAR or (**E**) CCL20-sensing gLIDAR with TEV output to control Citrine stability. Cells were gated on positive transfection marker population (mCherry+). (**F**) cLIDAR is able to directly output a secreted protein. (**G**) Direct secretion of secreted embryonic alkaline phosphatase (SEAP) via a rapalog-sensing cLIDAR. (**H**) gLIDAR is capable of interfacing RELEASE ^48^ to control protein membrane display by outputting TEV. ER-retained IgG: a membrane-bound human IgG1-Fc retained in the ER by a C-terminal -RXR motif with a flanking TEVP cut site in the middle. (**I**) IgG membrane display controlled by a CCL20-sensing gLIDAR. IgG1 surface expression levels are measured by staining with a AF488-conjugated anti-human IgG antibody. Cells are gated on positive transfection marker population (mCherry+). Refer to the **Methods** section for a detailed protocol on surface staining.

To further enable the manipulation of intercellular interactions, we were also interested in whether LIDAR could produce secretory outputs. We first demonstrated that cLIDAR is capable of outputting secreted embryonic alkaline phosphatase (SEAP) directly (**Fig. 3F-G**). Regarding gLIDAR, we started with testing the direct secretory output of a membrane-bound human IgG1 protein ^47^ (**Supplementary Fig. 8D-E**). Despite a significant increase in IgG membrane expression, the absolute levels are too low to be biologically relevant. We hypothesize that the low levels are due to the membrane tethering of the reporter RNA which reduced the translation capacity of secretory proteins that requires ER localization, so we decided to complement gLIDAR with our RELEASE (Retained Endoplasmic Cleavable Secretion) platform ^48^. We used TEVP output to control the protein secretion through the removal of endoplasmic reticulum-retention signals (**Fig. 3H**). Using this strategy, we observed a significant increase in membrane IgG display induced by the ligand, and the absolute membrane expression levels is much higher than that of direct membrane IgG output, though with a noticeable high baseline **(Fig. 3I and Supplementary Fig. 8 B-C**). This is likely due to the leaky expression of TEVP on the reporter RNA since the RELEASE platform is very sensitive to TEVP levels ^48^, which could be tackled by introducing more stop codons (**Supplementary Fig.3 H-I**). Taken together, LIDAR’s capability of outputting secreted proteins adds to its composability into complex circuits, and will enable intercellular communication essential for key applications we envision.

### Synthetic spatial patterning

Inspired by examples from synNotch ^10,11,50^ and given the advantage that LIDAR could sense both membrane-bound and soluble ligands, we next asked whether we could use LIDAR for synthetic spatial patterning, as a prelude to programming complex multicellular behaviors ^10,11,51^. We first validated that CCR6-based LIDAR indeed senses its ligand, CCL20, when the receiver cells are either exposed to conditioned media from CCL20-secreting cells (**Fig. 4A**) or co-cultured with cells displaying CCL20 on their surface (**Fig. 4B**) ^52^. We tested whether such contact-dependent and soluble ligand sensing features of LIDAR could be used to program distinct spatial patterns as a preliminary demonstration of spatial programming ^50^. When we plated receiver cells (monoclonal cLSM01) surrounding sender cells that secrete CCL20 (monoclonal cLSM03; **Supplementary Fig. 9A**), we observed a response gradient going from the sender cells outwards beyond the interface between sender (blue) and receiver cells (red) **(Fig. 4C-D; Supplementary Fig. 9B**). In contrast, when we tested receiver cells with sender cells producing membrane-bound CCL20 (monoclonal XZCP07), the response pattern is limited to the sender-receiver interface (**Fig. 4E-F; Supplementary Fig. 9B**). In this case, cell migration causes an overlap between the two populations, producing a band of activated cells that nevertheless remain within the interface of the two populations (**Fig. 4F**). Furthermore, we noted different output intensities produced by the membrane-bound and secreted ligand conditions (**Supplementary Fig. 9C**), presumably due to the different amounts of available ligand per receiver cell given the spatial differences of direct cell-cell contact vs. soluble ligand. In sum, these results demonstrate the feasibility of using LIDAR for juxtracrine and paracrine spatial patterning.

**Fig. 4.**
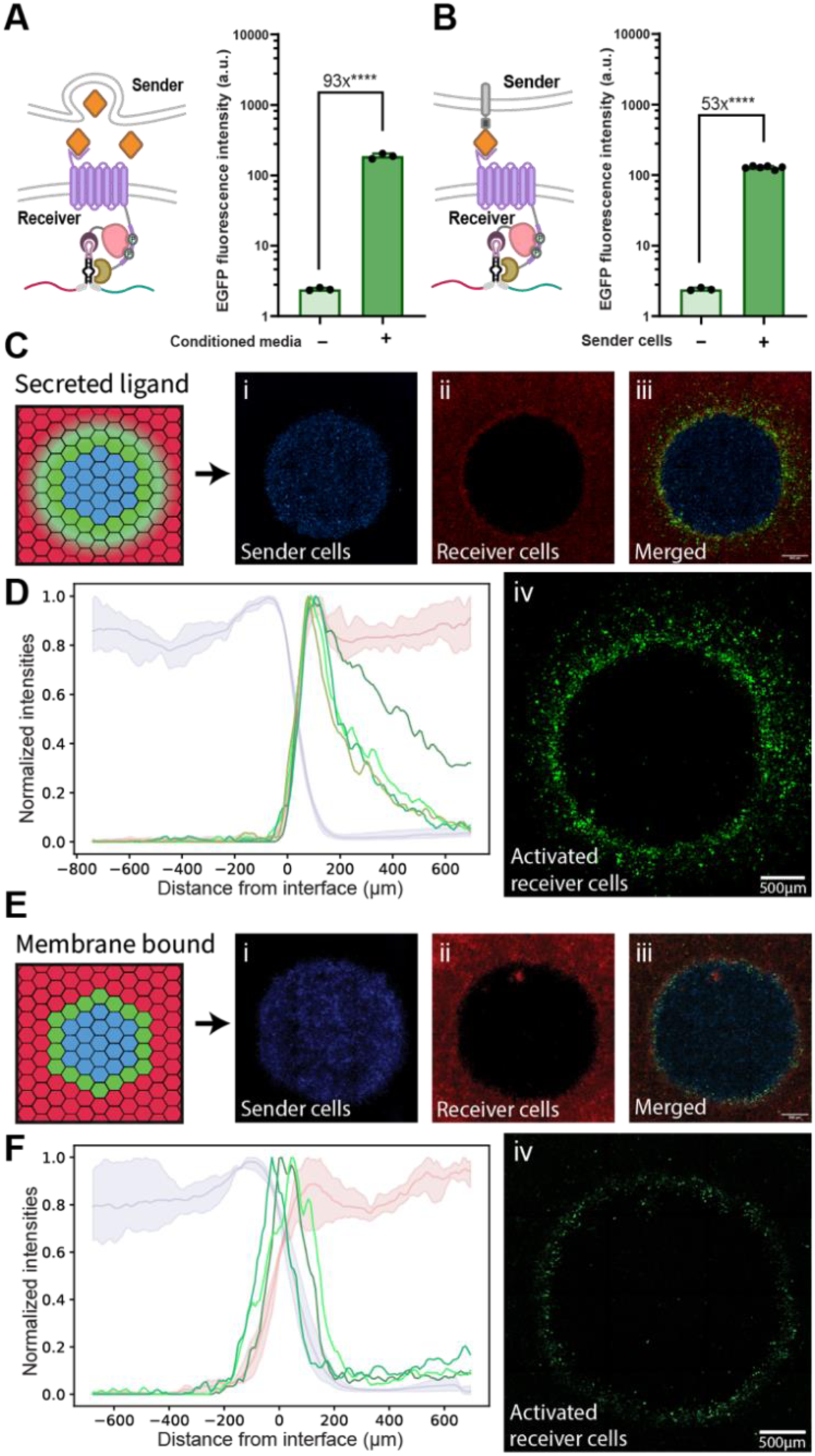
LIDAR as a tool to drive cell-cell interactions and synthetic patterning. **(A)** Left: LIDAR can detect soluble ligands secreted by other cells. Right: Data showing the response of transiently transfected receiver cells (with CCR6-based gLIDAR) treated with the supernatant from sender cells transiently transfected with soluble CCL20 or WT HEK cells. **(B)** Left: LIDAR is also capable of detecting ligands bound to the membrane of neighboring cells. Right: Data showing the response of transiently transfected receiver cells (CCR6-based gLIDAR) co-cultured with receiver cells transfected with membrane-bound CCL20 or WT HEK cells. See **Methods** section for more details and **Supplementary Fig. 9C** for data in stable cell lines. (**C-F**) LIDAR can be used to generate spatial patterns. **(C)** Sender cells producing secreted CCL20 (monoclonal cLSM03; TagBFP) were co-cultured with receiver cells (monoclonal cLSM01; mCherry) in a circular pattern to demonstrate LIDAR’s sensing capabilities and sensitivity. This setup produces a gradient of CCL20 from the center of the circular pattern outwards. The diagram to the left shows the experimental setup. Representative images of seeded sender cells (Ci), receiver cells (Cii), all the channels merged (Ciii), and the activated (EGFP) receiver cells (Civ) are shown on the right. See **Supplementary Fig. 9B. (D)** Normalized concentric intensity profiles of all channels for all replicates (n=4). **(E)** Sender cells producing membrane-bound CCL20 (XZCP07) were co-cultured with receiver cells (cLSM01) in a similar fashion as in (C). Individual panels are the same as for the secreted ligand condition (Ei-iv). For the membrane-bound CCL20, we see a thinner circular pattern, constrained to the sender-receiver interface. The overall GFP intensity is lower compared to the soluble ligand condition due to weaker activation (**Supplementary Fig. 9C**). **(F)** Concentric intensity profiles for the membrane-bound conditions (n=3). The plot in (F) shows that the peak mean GFP intensities stay within the interface between TagBFP and mCherry, supporting the activation of cells only when sender and receiver cells are in direct contact. In contrast, panel (D) shows mean GFP intensities extending beyond the interface and into the receiver cells (mCherry). For (D) and (F), an independent line is plotted for activated cells (green; EGFP), while sender (TagBFP) and receiver (mCherry) cells are plotted as the mean of all replicates with shades depicting maximums and minimums. A graphical demonstration of how the raw intensity values are measured can be found in **Supplementary Fig. 9D**. Refer to the **Methods** section for more information on the seeding and co-culture process, image acquisition, and processing and quantification of the intensity profiles. A linear enhancement of the brightness and contrast to aid with visualization were performed on all images as detailed in the **Methods** section.

### Unique delivery features

Lastly, we aimed to demonstrate the unique delivery features enabled by LIDAR’s post-transcriptional nature. Firstly, as many applications are limited by a delivery bottleneck, a post-transcriptional design could enable the compact encoding of all components. We encoded all LIDAR components on a single transcript by separating the components with encephalomyocarditis virus (EMCV) internal ribosome entry sites (IRES) (**Fig. 5A**), and we validated that the single-transcript performance is comparable to the corresponding multi-transcript version in terms of EGFP production (**Fig. 5B**, compared to **Fig. 1D**) as well as editing rated revealed by amplicon sequencing (**Fig. 5C**, compared to **Supplementary Fig. 5A**).

**Fig. 5.**
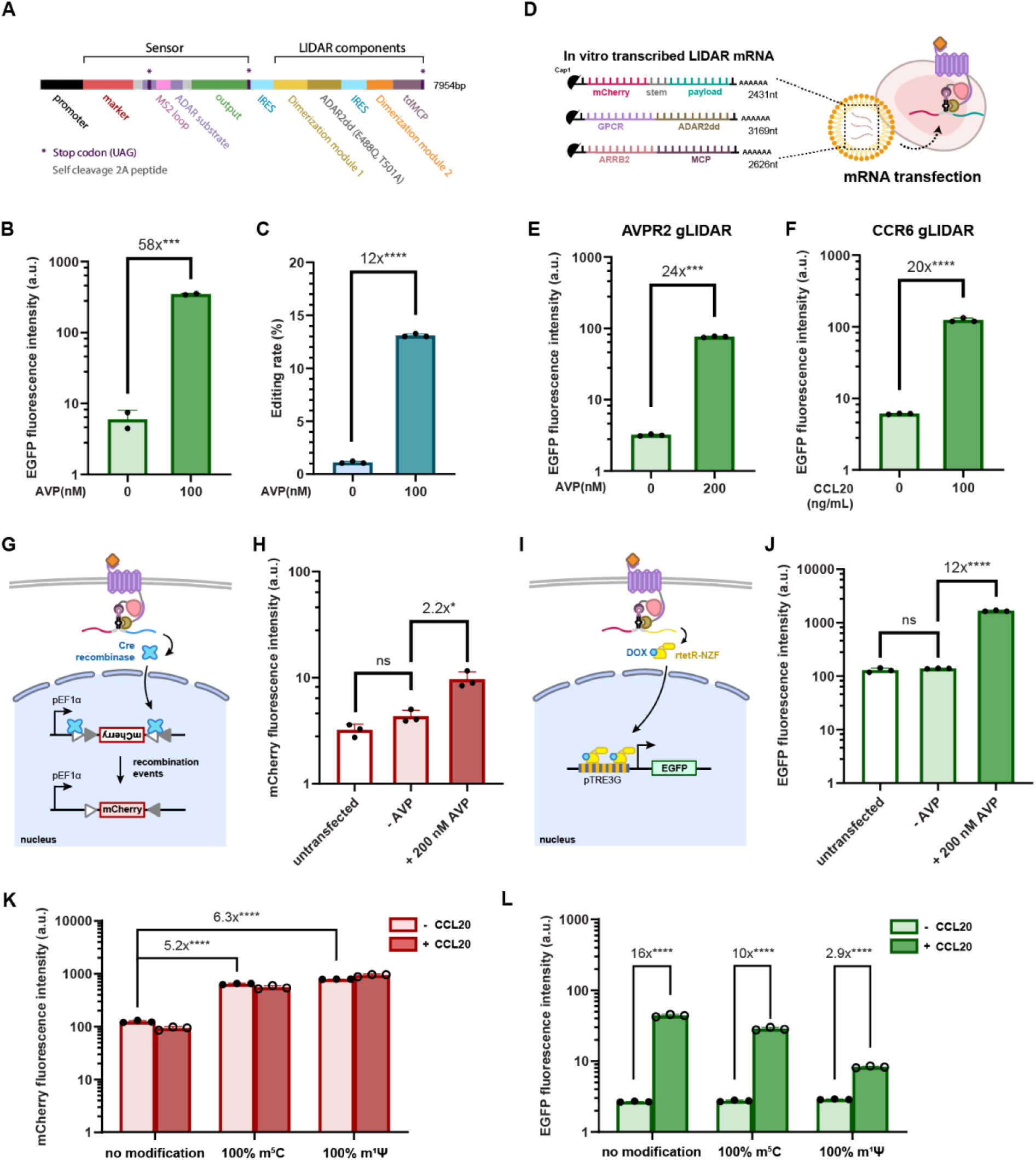
LIDAR can be encoded as a single transcript and is RNA-compatible. (**A**) Single-transcript encoding schematics of LIDAR. (**B**) EGFP activation by transient transfection of a single transcript-encoded AVP sensing gLIDAR. (**C**) Cognate target adenosine editing rate of (B) revealed by amplicon sequencing. (**D**) Schematics of LIDAR mRNA delivery setup. Each LIDAR component is encoded as an individual *in vitro* transcribed (IVT) mRNA strand and are delivered together via mRNA transfection. See **Methods** section for details on mRNA delivery. (**E-F**) EGFP activation by mRNA-delivered (**E**) AVP sensing gLIDAR and (**F**) CCL20 sensing gLIDAR using Thermo Messengermax transfection reagent. If not specified, all mRNA transfection data are gated on all transfection marker positive cells. (**G**) Schematics of mCherry production mediated by Cre recombinase output through mRNA-delivered LIDAR activation. The experiment is done in a cell line with genome integration of DIO-mCherry cassette, where the presence of Cre induces recombination that flips an inverted mCherry coding sequence, enabling the production of mCherry. (**H**) mCherry activation via AVP-sensing gLIDAR with Cre output assayed by flow 24 hours post mRNA transfection. (**I**) Schematics of EGFP production mediated by a transcription factor (rtetR-NZF) output through mRNA-delivered LIDAR activation. The experiment is done in a cell line with genome integration of TRE3G-GFP cassette. The rtetR-NZF output from LIDAR activates transcription of EGFP in the presence of Doxycycline (DOX). (**J**) EGFP activation via AVP-sensing gLIDAR with rtetR-NZF output assayed by flow cytometry 24 hours post mRNA transfection. (**K**) The effect of base modifications on IVT sensor mRNA in terms of marker expression. (**L**) The effect of base modifications on IVT sensor mRNA in terms of activation through CCL20 sensing gLIDAR.

Secondly, given our initial motivation for compatibility with RNA delivery, we tested whether LIDAR is still functional when encoded and delivered by mRNAs transcribed *in vitro*. We *in vitro* transcribed (IVT) all three LIDAR components into individual mRNAs (**Fig. 5D**). The mRNAs encoding the two protein components (CCR6-ADARdd and ARRB2-MCP) were synthesized with 100% of uridine substituted by N^1^-methylpseudouridine (m^1^Ψ) to enhance translational capacity, biological stability, and reduce immunogenicity ^53,54^. Since base modifications are known to affect stop codon recognition ^55^ and might potentially affect ADAR editing, we initially synthesized the LIDAR reporter mRNA without base modifications. We then showed that mRNA-delivered LIDAR produces strong EGFP output upon ligand induction using AVP-and CCL20-sensing gLIDAR (**Fig. 5E-F**), and demonstrate that the fold activation is comparable to that of DNA when using an optimized mRNA transfection strategy (**Supplementary Fig. 10A-C**). We further validated that the mRNA-delivered LIDAR system is capable of producing functional outputs. We demonstrated that mRNA-delivered gLIDAR is capable of outputting a Cre recombinase to induce mCherry expression in a DIO-mCherry cell line (cJKCC121; **Fig. 5G-H and Supplementary Fig. 10D**), and outputting a rtetR-NZF transcription factor to control genomic expression of EGFP in a TRE3G-EGFP cell line (cNK90; **Fig. 5I-J**).

Lastly, given that the output from unmodified synthetic LIDAR reporter mRNA might get hampered by its instability, we sought to assay if the LIDAR reporter mRNA is compatible with two types of base modifications commonly used in synthetic mRNA, m^1^Ψ and 5-methylcytosine (m^5^C) ^56,57^. LIDAR reporter synthesized with 100% m^5^C or 100% m^1^Ψ substitutions showed evident increase in overall marker expression, which indicates improved expression with base modifications (**Fig. 5K**); yet the EGFP output in response to ligand is dampened slightly with m^5^C and reduced heavily with m^1^Ψ (**Fig. 5L**). To assess the upper-bound performance of the reporter delivered as mRNA, we created cell lines that stably expresses ADAR2dd-MCP (cLSM13), ADAR2dd-PCP (cLSM14, which represents the background signal caused by ADAR2dd overexpression), and full-length ADAR2. The results are consistent with the conclusion that m^5^C is a more favorable modification than m^1^Ψ for LIDAR (**Supplementary Fig. S10E)**, and suggests that further optimization of the all-mRNA regime might bring the performance of modified mRNA to a level comparable to or surpassing unmodified mRNA. Taken together, our data shows that LIDAR can indeed be delivered by synthetic mRNA to produce functional outputs, and can accommodate modified bases.

## Discussion

In this work, we present LIDAR, a novel platform for building post-transcriptional synthetic receptors. The use of standalone ADAR editing as an actuation mechanism enabled hypothesis-driven and iterative engineering, resulting in a modular receptor platform compatible with various receptor architectures (**Fig. 1, 2**), output types (**Fig. 3**), complexities (**Fig. 4**) and cellular contexts (**Fig. 2L-O**). These characteristics, combined with compact encoding and compatibility with mRNA delivery (**Fig. 5**), could eventually enable broad applications in basic research and biomedicine.

Building on the foundation established by previous modular synthetic receptors, LIDAR both inherits some of their features and offers unique benefits. The modular nature of LIDAR allows us to leverage prior advancements in receptor engineering to develop RNA-compatible versions of MESA ^32^, Cha-Cha ^39^, and TANGO ^33^. Inspired by recent advances in MESA ^34,58^, we built eLIDAR, receptors capable of sensing recombinant ligands with relatively straightforward designs. Similarly, as demonstrated by our best-performing LIDAR architecture, gLIDAR can achieve low baselines (**Supplementary Fig. 3A**) while still maintaining ligand-induced signals comparable to those of a transcription-based synthetic receptor (**Supplementary Fig. 1J)**. Given the (often non-linear) two-step amplification in transcriptional synthetic receptors, and through our personal experiences and communications, careful dosage or engineering is needed to mitigate runaway baselines ^59,60^. We postulate that LIDAR might have minimized the baseline by employing a more limited degree of amplification, yet further comparisons are needed to confirm this across contexts. Further, although **Fig. 3** demonstrates that LIDAR can output sufficient levels of proteins to alter cellular phenotypes, in scenarios requiring stronger outputs, transcriptional receptors may still be more suitable. Therefore, albeit RNA compatibility is the most distinctive feature of LIDAR, such nuanced comparisons are crucial when selecting the appropriate receptor for a specific application. Additionally, we have identified that LIDAR is sensitive to high levels of ADAR1/2 expression (**Fig 2L-M, Supplementary Fig. 4F**) but presents no other apparent source of leakiness. To tackle the baseline issue in those specific scenarios, we have designed a more stringent reporter RNA by including tandem stop codons (**Supplementary Fig. 3I)** and can further adjust the stoichiometry of components to modulate the signal (**Supplementary Fig. 4)**. It’s worth noting that due to our gating process, a very high transfection efficiency can lead to disproportionately high signal intensities and baselines, compared to less efficiently transfected experiments. Therefore, comparisons across panels and cell lines should be made with caution. For instance, the rapalog-sensing eLIDAR in **Fig. 2L** was repeated in **Supplementary Fig. 4C**, but the latter exhibits a much lower baseline.

Further optimizations of LIDAR are required to prepare it for the envisioned applications. On the functionality front, due to the bypass of transcription, LIDAR could potentially exhibit a faster response than transcription-based synthetic receptors, yet the current signal upon ligand activation still takes over a day to saturate (**Supplementary Fig. 1H-I**). We will systematically identify the rate-limiting steps in LIDAR operation to optimize its kinetics. Additionally, while RELEASE enables LIDAR to indirectly secrete proteins at biologically relevant levels, we are investigating the challenges associated with direct secretion of output proteins and exploring alternative strategies that could streamline the system. We also acknowledge that for LIDAR receptors that leverage endogenous compartments (i.e. gLIDAR), activation of native signaling pathways alongside with LIDAR activation could be a limiting factor for certain applications. On the delivery front, firstly, although compact encoding by DNA is achieved, compact encoding on IVT mRNA has remained challenging, potentially due to reduced efficiency in mRNA synthesis and decreased stability with longer transcript ^61–63^, while such single-RNA encoding will help ensure the optimal stoichiometry of LIDAR components. We envision that this could be addressed through advances in mRNA synthesis protocols^61^, incorporation of stabilizing elements on mRNA^63^, and optimizing IRES to enhance translational efficiency ^64^. Secondly, careful considerations and more comprehensive assays should further be taken when incorporating base modifications on reporter mRNA since we have seen a tradeoff between overall protein production and ADAR editing efficiency regarding incorporation of m^5^C and m^1^Ψ. Other stabilizing and de-immunizing base modifications could potentially be considered, and it’s also worth testing an optimal ratio between combinations of different base modifications. We will also explore alternative RNA synthesis protocols to synthesize a base modification-free stem-loop region, then using splint ligation to connect it to the rest of the reporter RNA harboring most stabilizing base modifications ^65^. Lastly, we are aware that our demonstrations on functional output of LIDAR mRNA delivery still rely on transcriptional activation that involves synthetic compartments. With optimizations on mRNA delivery, we are seeking functional outputs that are compatible with a more “endogenous” cellular context for practical applications, including outputting a transcription factor that tunes endogenous gene expression or an effector protein that directly alters cellular physiological states.

The current iteration of LIDAR already offers useful tools, and, pending future improvements, it has the potential to empower even more applications in both basic research and biomedicine. Firstly, there are exciting opportunities to generalize LIDAR towards user-defined inputs beyond leveraging existing receptor scaffolds, particularly given recent advancements in Programmable Antigen-gated G protein-coupled Engineered Receptors (PAGERs), with which LIDAR is mechanistically compatible ^66^. In terms of applications, a rapalog-responsive LIDAR can be utilized directly as a post-transcriptional inducible system, either on its own for scenarios where transcriptional control is unfeasible (e.g., delivery using an RNA viral vector), or in synergy with transcriptional control to improve specificity or signal-to-noise ratio. For basic research, LIDAR could be used to visualize or manipulate specific neurons based on their exposure to endogenous neurotransmitters or connection with other neurons. By reporting activities integrated over time, LIDAR could complement those sensors that have high temporal resolutions ^67,68^ when the change of activities occur over longer time periods or invasive measurements are less feasible. More importantly, with the emergence of technical advances in synthetic mRNA production and delivery, LIDAR has the potential to offer unique capabilities to next-generation therapeutics. First, similar to the use of SynNotch to modulate the efficiency and specificity of *ex vivo* engineered CAR T cells ^69–71^, LIDAR would be able to regulate *in vivo* CAR T cells generated by mRNA delivery. Second, we will more thoroughly and rigorously test LIDAR in cancers, so that it could be applied to precisely deliver cargos into cancer cells to ablate the tumor and engage the immune system. Beyond removing anomalous cells, LIDAR could also help create beneficial cell types or tissues by processing and guiding spatial cues, again using mRNA to transiently modify endogenous cells without the need for cell extraction and transplantation. Taken together, we are hopeful that future iterations of LIDAR will inhabit an opportune position at the junction between RNA medicine and synthetic biology.

## Acknowledgments

This research was supported by NIH (R00EB027723, R21EB033858, DP2OD034951; X.J.G), Longevity Impetus Grants (X.J.G.), Stanford Bio-X Interdisciplinary Initiatives Seed Grant Program (IIP) [R11-6] (X.J.G.), Stanford Cancer Institute Innovative Award (X.J.G), Stanford Interdisciplinary Graduate Fellowship (X.Z., L.S.M., K.E.K.), the Stanford Graduate Fellowship (L.S.M., J.K., Y.H.), the Sarafan ChEM-H CBI training program (L.S.M.), the Sarafan ChEM-H/IMA Postbaccalaureate Program in Target Discovery (A.P.L.), the CMB training grant NIH (T32 GM007276; C.C.C.), and the National Science Foundation Graduate Research Fellowship (DGE-2146755; C.C.C.).

We thank Prof. Lacramioara Bintu for gift of the HEK293 cell line, Prof. Howard Chang for gift of the Colo 320DM cell line and Prof. Liqun Luo for the gift of Drosophila S2 cells. We thank Prof. Alice Ting for membrane-bound chemokine and their receptor plasmids. We thank Andrew Lu and Prof. Michael Elowitz for the IVT plasmid backbone. We thank Prof. Qin Li for guidance on analyzing sequencing data. We thank Siranush Babakhanova for testing early versions of LIDAR. We thank the Gao lab members for their feedback. We thank Noa Katz for tTA reporter plasmid and cell line, Jenny Kang for DIO-mCherry cell line, and Alex E. Vlahos for plasmid encoding IL-2 and RELEASE.

## Author Contributions

XZ, LSM, KEK and XJG designed the study. XZ, LSM, KEK, CCC, MZ, YH and XJG developed the methodologies. XZ, LSM, KEK, APL, CCC, MZ and YH performed the experiments. XZ, LSM, APL, CCC, MZ and YH analyzed and processed the data. XZ, LSM and XJG wrote the manuscript with input from all authors.

## Materials and Methods

### Plasmid generation

Plasmids were generated using standard molecular cloning practices, including InFusion of linearized plasmids and PCR fragments, restriction–ligation of linearized fragments and annealed phosphorylated oligonucleotides. A complete list of plasmids and associated maps can be found in Supplementary Table 1. Plasmids are available upon request from the corresponding author and are deposited to Addgene. Plasmid sources for VEGFR1, VEGFR2, AVPR2, β-arrestin2 (ARRB2), OXTR, B2AR were acquired from Addgene. Plasmid sources for CCR6, CCR7 as well as their cognate ligands were gifts from Prof. Alice Ting. Plasmid sources for other components are synthesized through Twist Bioscience.

### Tissue culture

All cell lines were cultured in a humidity-controlled incubator under standard culture conditions (37 °C with 5% CO2). HEK293 cells (catalog no. CRL-1573) and SNU-475 cells (catalog no. CRL-2236), purchased from ATCC, were cultured using Dulbecco’s modified Eagle’s medium, supplemented with 10% fetal bovine serum (FBS) (Fisher Scientific catalog no. FB12999102), 1 mM sodium pyruvate (EMD Millipore catalog no. TMS-005-C), 1× penicillin–streptomycin (Genesee catalog no. 25-512), 2 mM L-glutamine (Genesee catalog no. 25-509) and 1× MEM non-essential amino acids (Genesee catalog no. 25-536). C3H mouse fibroblast cells (ATCC catalog no. CCL-226) were cultured according to ATCC’s suggested protocol using Eagle’s Basal medium (Thermofisher catalog no. 21010-046) supplemented with heat-inactivated FBS to a final concentration of 10%, 1% Penicillin/Streptomycin and 2mM L-glutamine (ATCC 30-2214). Similarly, Colo 320DM cells (gift from Prof. Howard Chang) were cultured according to ATCC’s suggested protocol. RPMI medium supplement with 10% FBS and 1% Penicillin/Streptomycin was used to culture these cells. Finally, Drosophila Schneider 2 (S2) cells (gift from Prof. Liqun Luo) were cultured in tissue culture-treated 10 cm plates at 20°C–22°C, outside an incubator and without antibiotics and in Schneider’s Drosophila Medium (ThermoFisher Scientific catalog no. 21720024) with 10% heat-inactivated FBS.

### DNA Transient Transfection

All cells were cultured in 24-well or 96-well tissue culture-treated plates under standard culture conditions. When cells were 70–90% confluent, they were transiently transfected. HEK293, SNU-475 and Colo 320DM were transfected with plasmid constructs using the jetOPTIMUS DNA transfection reagent (Polyplus catalog no. 117-15), as per manufacturer’s instructions using 0.375 µl of reagent per 50 µl of jetOPTIMUS buffer for 500 ng total DNA transfections in the 24-well format and 0.13 µl of reagent per 12.5 µl of buffer for 130 ng total DNA in the 96-well format. C3H cells (24-well) were transfected using Lipofectamine 3000 as per manufacturer’s instructions (ThermoFisher catalog no. L3000001). Finally, S2 cells (24-well) were transfected when they reached a cell count of 2×10^6^– 4×10^6^ cells/mL using Lipofectamine 2000 as per manufacturer’s instructions (ThermoFisher catalog no. 11668019). All transfections are detailed in Supplementary Table 2.

### Flow cytometry and data analysis

For DNA transfections, cells were harvested by trypsinization and resuspended in flow buffer (HBSS + 2.5 mg/mL bovine serum albumin) approximately 48 h after transfection and 24 h after adding the ligand. For RNA transfections, unless specified, cells were harvested 24h post-transfection with the same preparation process. After 40-µm straining, cells were analyzed by flow cytometry (Biorad ZE5 Cell Analyzer). For quantitative analysis on fluorescence intensities, data was processed with the CytoFlow Python package, and cells were gated on the highly transfected populations based on mCherry levels. Unless specifically stated, the 99.5 to 99.9 percentile mCherry (transfection marker) level of the lowest-transfected replicate is used to gate all samples within a comparison set as done in our previous work ^72^. In most figures, each plot consists of a comparison set in one experiment. In a few cases, we use broken x-axes to indicate separate comparison sets from different experiments within a plot. Unless specifically mentioned, all bars in one comparison set are gated using the same mCherry fluorescence gate. Values are reported as the means of at least three (in main figures) or two (in supplementary figures) biological replicates. For experiments comparing multiple groups, a Bonferroni-corrected two-tailed Student’s t-test was used to assess significance. * indicates P<0.05, ** indicates P< 0.01, *** indicates P< 0.001 and **** indicates P< 0.0001. For visualization and analysis on populational distributions, FlowJo software was applied.

### Cell surface staining

Cells in 24 well plate were harvested 48 hours post-transfection for surface staining by dissociation with CellStripper (Corning #25-056-CI) for 15 minutes, then washed twice with cold flow buffer. Cells were then incubated with 100 μL of corresponding antibody dilutions in flow buffer for 30 minutes at 4 °C. For staining eLIDAR surface expression, 1:100 dilution of anti-Flag-APC (abcam #ab72569) and 1:200 dilution of anti-HA-AF488 (BioLegend #901509) were used. For staining membrane display of IgG, 1:750 dilution of anti-human IgG-AF488 (Southern Biotech 2015-30) was used, the same dilution factor is applied on the isotype control antibody (Southern Biotech 0109-30). After incubation, cells were washed twice with flow buffer, then filtered through a 40 μm sieve to be prepared for flow cytometry.

### Cell line construction and selection

Polyclonal cell lines were generated using piggyBac stable integration, by co-transfecting the cargo plasmids together with a helper plasmid expressing the transposon (see **Supplementary Table 2**) into WT HEK293 cells. eLIDAR cell lines were generated through a double piggyBac integration, where the first construct is a LIDAR sensor RNA driven by SFFV with the selection marker (hygromycin resistance gene) driven by pSV40; the second construct is the two LIDAR receptor arms encoded in one transcript separated by an EMCV IRES site driven by EF1a promoter and a separate pSV40 driven puromycin resistance gene with P2A-TagBFP. Ligand-expressing sender cell lines in **Fig. 4**, were generated using a single transcript containing BFP and puromycin as markers. gLIDAR receiver cell lines were also generated through a double piggyBac integration, similar to that of eLIDAR. The first construct is the same as for eLIDAR; the second construct is a EF1a-driven GPCR-ADARdd with ARRB2-MCP separated by an IRES site and a T2A-puromycin resistance gene at the end of the transcript. Cell lines used for off-target editing analysis in **Supplementary Fig. 6** and ADAR2 overexpressing cell lines in **Supplementary Fig. 10E** were also generated using piggyBac stable integration and a single plasmid containing the specific protein, and BFP and puromycin as markers. Monoclonal cell line for apoptosis induction (cLSM01) were further selected by diluting polyclonal cell line to an average of 0.2 cell per well in a 96 well plate in selection media. Optimal monoclonals were selected based on their capacity to produce EGFP and start the apoptotic pathway. See **Supplementary Table 1 and 3** for the specific plasmids and their ratios used for integration.

### RNA Extraction & amplicon generation

Cells were seeded in 24-well plates for transfection and/or treatment with the right ligand. Each well was harvested as described in the **Flow cytometry and data analysis** section and spun down to form a pellet. Qiagen’s RNeasy Mini Kit (Qiagen, Cat. No. 74104) was used exactly as described in their protocol to extract and purify RNA from the samples. Each pellet of cells was disrupted and homogenized using a QIAshredder column (not provided in the kit). After RNA extraction, a second in-solution DNA clean-up was performed only for samples that were transiently transfected to ensure full removal of plasmid DNA. Final RNA concentration was measured using a Nanodrop and ensuring both ratios (260/280 and 260/230) were above 2.0. RNA was either directly used for RNA-seq or reversed transcribed into cDNA using BioRad’s iScript (#1708840). A total of 1µL was used for cDNA synthesis. Amplicons of the reporter RNA were generated by running a 100µL PCR for 17-20 cycles using 20µL of diluted RNA (1:1 dilution). The exact number of cycles was dependent on whether a band could be seen for all samples after preliminary PCRs (10µL reactions; 17, 18, 19 and 20 cycles). The final 421bp amplicons were purified using Zymo DNA clean-up kit (DNA Clean & Concentrator-5; Zymo Research; Cat # D4003).

FWD primer: acactctttccctacacgacgctcttccgatctcgcgttggaagccagacacaaacag

REV primer; gactggagttcagacgtgtgctcttccgatctcctttgagacactcgagggtcctggattag

### Amplicon sequencing and data analysis

After successful amplicon generation and purification, 25µL (20ng/µL) per sample were submitted to Genewiz for NGS (Amplicon EZ; ∼50,000 2×250bp reads per sample). Results were analyzed via a custom python script. Briefly, pair reads were fused into single reads using FLASH - Fast Length Adjustment of SHort reads ^73^. After that, reads were filtered to keep only those with a mean QC > 30. Two substrings corresponding to the modified (CCTCCGTTTGGGTGGGTGG) and unmodified (CCTCCGTTTAGGTGGGTGG) reporter were compared against the resulting reads and counted to identify the portion of reads that were edited. The code used to analyze the data can be found in the github repository.

### RNA Sequencing and transcriptome data analysis

Purified RNA was sent to Novogene for RNA Sequencing (Human RNA Sequencing WOBI; NovaSeq X Plus Series PE150; 30M reads per sample). Three samples per condition were submitted and later combined to obtain a total of ∼90M reads/condition. Data was analyzed as described in Katrekar et al. ^29^ with subtle modifications. First, raw filtered reads from Novogene were aligned to a reference genome using STAR. The genome indices used for alignment were built using both, a primary assembly annotation from GENCODE (GRCh38.v46) and, depending on the sample, the DNA sequence that was integrated into the sample cells being analyzed. Default parameters were used to run STAR, except for the following relevant settings: runThreadN = 12, outSAMtype = BAM SortedByCoordinate, quantMode = GeneCounts, readMapNumber = −1, alignSJoverhangMin = 1, alignSJDBoverhangMin = 1, alignEndsType = Local, outFilterMultimapNmax = 1, outSAMunmapped = None, outSAMmultNmax = 1, outReadsUnmapped = Fastx, outFilterMismatchNoverLmax = 0.25.

The uniquely aligned pairs were further processed to identify duplicate reads using samtools markdup ^74^ and were removed from subsequent analysis. To avoid any potential bias due to the differences in number of aligned reads when comparing different samples in terms of significant edited sites, all samples in a same analysis group were down sampled to a fraction of their original number using samtools (samtools view -s). The down sampling fractions are reported in Supplementary Table 4. These fractions were calculated by dividing the lowest number of uniquely aligned reads among all samples in the group by the number of aligned reads available for the sample being down sampled ^75^.

To extract the coverage and editing frequency per genomic position, the down sampled reads were passed through REDItools ^75^. The default parameters were used expect for the following flags: -e -d -u -n 0.01 -v 1, which cause REDItools to exclude duplicate reads and multi hits, consider mapping quality, modify the minimum number of reads supporting the variation to 1 and the minimum frequency to 0.01. The resulting sites were further analyzed for each sample to identify those A and T sites with at least one read supporting an ADAR edit (A-G and T-C in the reverse strand). A final list of A and T sites was selected by choosing those sites that were common to all samples in the analysis group, this means all samples have a coverage of > 19 reads. In some cases, those where we observed an editing event in a sample with coverage > 19, we included the site in the final list if one of the other samples of zero editing frequency had a coverage > 9. The other sites, those not common to all samples or with no editing events in all samples, were discarded.

Finally, to identify statistically significant changes in editing between pair comparisons for each reference A or T site selected as described above, a Fisher exact test was carried out using a 2 × 2 contingency matrix (or Chi-square if all values of the contingency matrix were larger than 5). The *P* values calculated for all selected reference sites and for a given comparison of samples were adjusted for multiple testing using the Benjamini–Hochberg method (using Python’s statsmodel multitest function). A and T sites with adjusted *P* values less than an FDR of 1% and with a fold change of at least 1.1 in editing yield were selected as significant. The number of significant editing events are shown as *N*_sig_ and plotted in red in each panel in Supplementary Fig 8. On-target A-to-G editing yield on LIDAR’s reporter RNA is shown with a green arrow in Supplementary Fig 8A. Please refer to the github repository for more details on each script used to run this analysis.

### Co-Culture Experiments

For experiments where cells were transiently transfected, CCL20 sender cells and CCR6-based LIDAR receiver cells were transfected independently. CCL20 sender cells were transfected a day before the CCR6-based LIDAR receiver cells – to allow for the production and secretion of CCL20 into the media – and 24 h after the transfection of the receiver cells, the supernatant of the sender cells was transferred into the receiver cells. The output of the receiver cells was measured using flow cytometry 24h after supernatant transfer. For the membrane-bound CCL20 experiments, sender and receiver cells were transfected separately and then lifted and cultured together 24 h later. The output of the receiver cells was measured using flow cytometry 24 h after co-culture. For experiments using the generated cell lines, cells were cultured together – for both membrane-bound and secreted ligand conditions – and receiver cells were measured after 24 h.

### Co-Culture Experiments for Pattern Formation

Custom Polydimethylsiloxane (PDMS, Fisher Scientific #NC9285739) inserts were fabricated using 3D-printed resin molds and 1 (or 2) mm biopsy punches (Fisher Scientific #12-460-402). Printed molds were washed with isopropanol after printing, further cured under UV light for 30 min and placed at 75C for 8h to ensure the reaction had finished. After the mold was fully cured, PDMS was mixed 10:1 (w/w) (base:curing agent), de-gassed, poured into the first mold inside a petri dish and cured at 75C for 1-2 h. Resulting PDMS pieces were further processed using the biopsy punches to make a hole in the center and produce the final PDMS insert. A second mold was used to ensure the punched hole was centered. Final PDMS inserts were sterilized using the autoclave or by placing them under UV light for 30 min inside a petri dish with 70% ethanol. After sterilization, inserts were placed inside a well of a 24 well plate. Monoclonal receiver (cLSM01) and sender (XZCP07 or cLSM03) cells were lifted, spun down, resuspended on fresh media, counted and diluted to 10^6^ cells/mL. Sender cells were seeded (5uL of diluted cells) inside the insert and receiver cells on the outside (120 uL of diluted cells). The insert was removed 16-24 h later. Cells were incubated on top of a rocker at the lowest speed to ensure uniform diffusion. Images were taken 1-2 days after the insert was removed to allow cells to occupy the empty space (Supplementary Fig. 4A).

### Cell Death Assay

Monoclonal cLSM01 cells were seeded in a 24 well plate using 0.5 mL/well of resuspended cells (10^6^ cell/mL, final concentration) to ensure all the wells had the same initial confluency. Recombinant CCL20 was added 12 h after seeding and AP1903 was added 24 h after seeding. Refer to Supplementary Table 3 for ligand catalog and concentration used. Images were taken and flow cytometry was performed 12 after adding (36 h after seeding) AP1903. The full process was done within a doubling cycle (36 h) to primarily account for the effect of cell death and eliminate any interference coming from potential differences in growth rates between wells and conditions.

### Annexin V Staining

Cells were treated and lifted as described in **the Cell Death Assay** section. Cells were spun down to remove media and later resuspended in the annexin-binding buffer provided in the Invitrogen’s Annexin V Ready Flow Conjugates kit (Cat No. R37177). 8uL of Annexin V conjugate was added per 100 uL of cell suspension in binding buffer. After a 15 minutes incubation at 25C, cells were prepared for flow as described in Flow cytometry and data analysis after 40-μm straining.

### Image acquisition and processing

All images were taken using an EVOS M7000 Cell Imaging System. Co-culture cell patterning images were taken using the automated tile-stitching feature of the imaging system. Co-culture images were processed using ImageJ (FIJI for MacOS) by first subtracting the background for all channels (Process->Subtract background…) using a 100-pixel window (“rolling ball” in ImageJ) and then by linearly enhancing the brightness and contrast of the images to aid with visualization. Composite images were also generated in ImageJ by merging the images of previously processed individual channels (Image->Color->Merge channels). Cell death images were only processed to linearly enhance contrast and brightness. For the images in the first column of Supplementary Fig. 3A, binary masks were generated by segmenting (i.e., thresholding and binarization) the dead cells in the bright field images. Segmentation was done based on morphological differences between healthy and dead cells, i.e., the difference in brightness and roundness of dead cells compared to live cells. Final cell death images were generated by overlaying bright field images with the masks corresponding to death cells. For both co-culture and cell death figures, each channel was processed independently.

### Pattern Formation experiments quantification and plotting

Previously processed green channel images were analyzed using a MATLAB script. Average intensities were calculated from the center of the circular patterns outwards (Supplementary Fig. 4D) and then these values were normalized to account for different background and intensity values in each individual image. Given that the different circular patterns obtained had slightly different diameters due to cell growth, the data was further processed to align global maximas corresponding to the interface between the two populations of cells. A low-pass filter was also applied to the data by using a 3-value moving average. The final plots are shown in Fig. 4E. The codes used to process this data can be found in the supplementary materials.

### mRNA synthesis and transfection

The DNA templates for in vitro transcription were prepared by PCR on a plasmid template containing the LIDAR coding sequences flanked by optimal 5’ and 3’ UTRs as well as a T7 promoter. The plasmid template with optimal UTRs was a gift from Prof. Michael Elowitz. Poly-A tail was added to the DNA template by PCR with reverse primer containing a 200bp polyA sequence. In vitro transcribed mRNA was produced and purified in-house based on the DNA template, with CleanCap® Reagent AG (N-7113) as the capping analog and optional base modifications incorporated. The purified mRNAs were then transfected to specific HEK cell lines as needed using Thermo MessengerMax transfection reagent or Mirus TransIT®-mRNA Transfection Kit following the manufacturer’s instructions. A total of 1ug mRNA are co-transfected together per well in 24 well plate as 200 ng LIDAR reporter, 400 ng GPCR-ADARdd and 400 ng ARRB2-MCP. In addition, cells were transfected when they reached >90% confluency. When using Thermo transfection reagent, 1.5uL of transfection reagent are used per well in 24 well plate. When using Mirus transfection reagent, 3 μl of TransIT-mRNA reagent as well as mRNA boost was used per well in 24 well plate. The corresponding ligand was supplied together with mRNA transfection. Fluorescent protein expression was assayed through flow cytometry 24 hours post mRNA transfection if not specified.

### Data Availability

Plasmid maps can also be found in Supplementary Table 1. Raw data, as well as the code used for its analysis, is available upon request from the corresponding author.

## Supplementary Materials

Supplementary Figure 1

Supplementary Figure 2

Supplementary Figure 3

Supplementary Figure 4

Supplementary Figure 5

Supplementary Figure 6

Supplementary Figure 7

Supplementary Figure 8

Supplementary Figure 9

Supplementary Figure 10

Supplementary Table 1. List of plasmids (LIDAR)

Supplementary Table 2. Overview of transfections

Supplementary Table 3. Overview of cell lines

Supplementary Table 4. List of ligands

Supplementary Table 5. Down-sampling fractions for experiment in Supplementary Fig. 6.

