## Supplementary Data for "Post-Transcriptional Modular Synthetic Receptors"

### Supplementary Materials

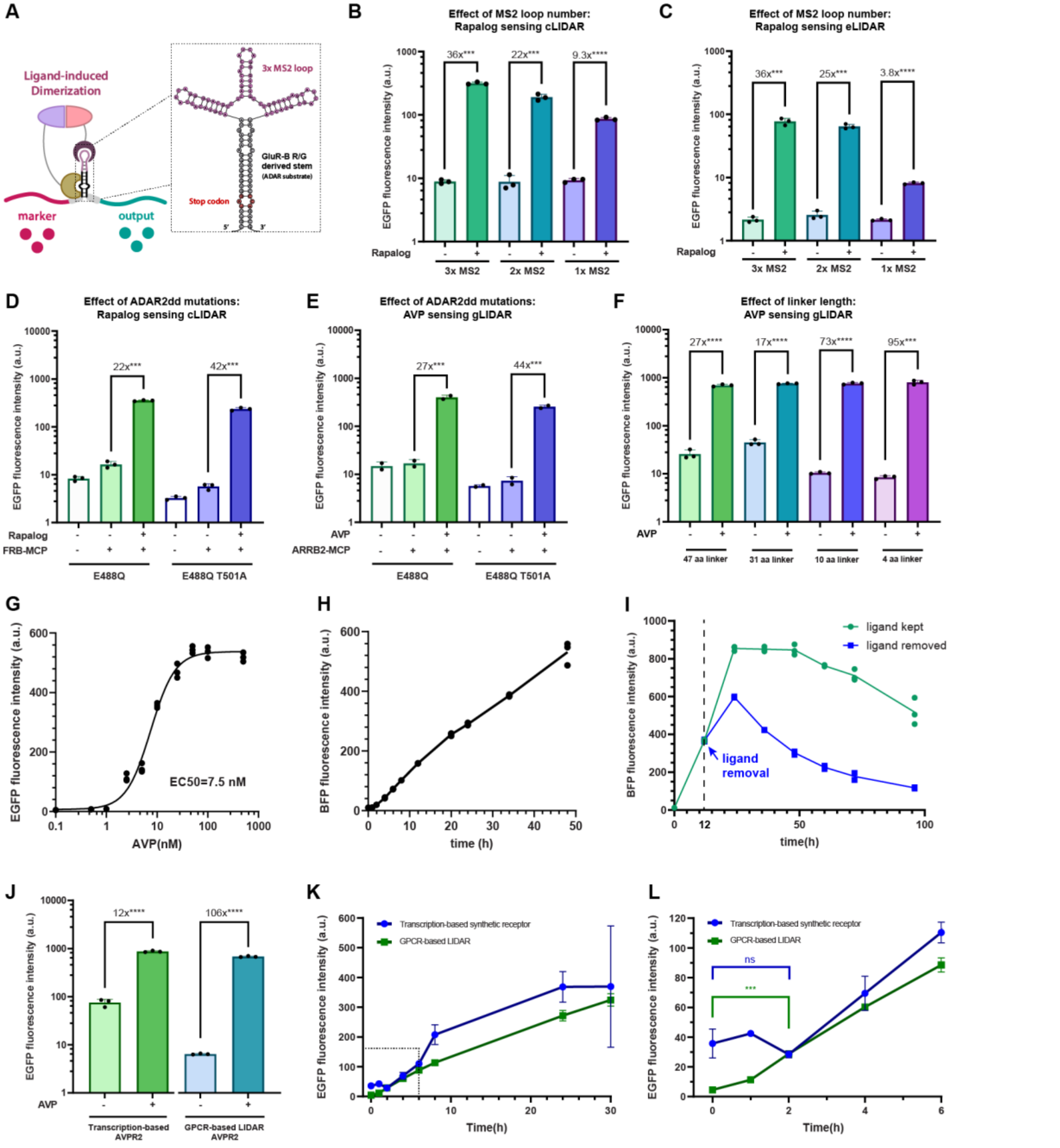


**Supplementary Fig. 1. Optimization and characterization of the LIDAR components.** (**A**) Schematic of the LIDAR reporter RNA and its predicted folding structure via RNAfold [1]. A GluR-B derived stem loop is used as the ADAR substrate [2] and 3x MS2 loops are included in this design to ensure strong MCP recruitment. (**B**) Data showing the differences between having 1, 2 or 3 RNA binding protein substrates (i.e., MS2) loops in the output levels for cLIDAR and (**C**) eLIDAR. (**D-E**) Supporting data for the rationale behind using a E488Q, T501A double-mutant ADAR2dd for LIDAR. The effects of ADARdd mutations were evaluated in (**D**) cLIDAR and (**E**) AVPR2-based gLIDAR. The results for panels (B-E) are representative and can be extrapolated to other LIDAR designs. (**F**) Data showing the effect of linker lengths between the fusion of GPCR C-terminal tail and ADARdd of AVPR2-based gLIDAR. A shorter linker lowers down baseline and yields high fold activation. (**G**) Dose-response curve for AVPR2-based gLIDAR via transient transfection. EC50 was estimated through fitting to a 4-parametric logistic function. (**H**) LIDAR activation as a function of time after adding 200 nM AVP in cLSM02 cell line (AVPR2-based gLIDAR stably integrated with a sensor that has mCherry as marker and TagBFP as output). (**I**) LIDAR signal dynamics after ligand removal in cLSM02 cells. Cells in both conditions were induced with 200 nM AVP for 12 hours and then, for one of the conditions, AVP was removed by replacing the media. For the other condition, media was also replaced and 200 nM AVP was added to continue the time course. Cell samples are taken for flow cytometry every 12 hours afterwards until 92 hours post initial ligand induction. (**J**) Direct comparison between a TF-based synthetic receptor based on TANGO [3] and LIDAR. Both receptors are based on the same GPCR (AVPR2) via transient transfection. (**K**) Overlaid time-response curves of an AVPR2-based gLIDAR and the TF-based synthetic receptor. (**L**) Close-up of the first 6 h after ligand induction (boxed region in (K)) depicts a statistically significant activation for LIDAR compared to lack of significance for the TF-based synthetic receptor.

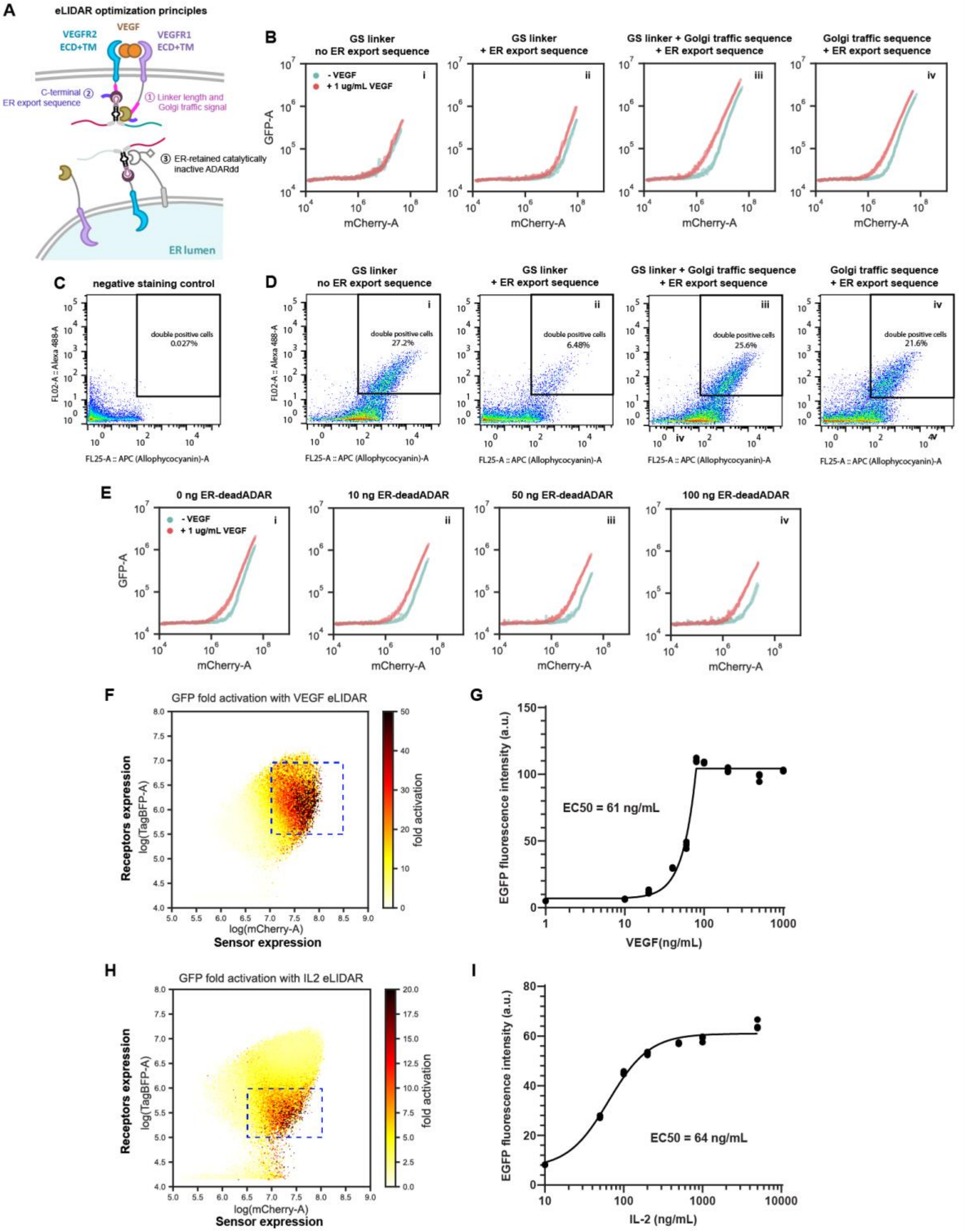


**Supplementary Fig. 2. Optimization steps towards building modular eLIDAR system.** (**A**) Schematics of eLIDAR optimization principles. Optimizations are done by assaying the effect of 1) linkers and Golgi transport sequence between the transmembrane domain and LIDAR intracellular domain (ICD); 2) C-terminal ER export sequence; 3) introduction of ER-retained catalytically inactive ADARdd (E396A). (**B**) The effect of linkers, Golgi traffic sequence and ER export sequence[4] on eLIDAR sensitivity to ligand revealed by running means plots. Running means plots are generated by calculating the rolling means of output (EGFP) levels on transfection marker (mCherry) expression using batch size = 1000. Each running means plot is comprised of duplicates of each transfection condition +/- ligand (VEGF-165). Each duplicate is plotted as a line with 50% transparency. GS linker: *GTGGSGSAS*; GS linker + Golgi traffic sequence: *GTGGSGSA*KSRITSEGEYIPLDQIDINV; Golgi traffic sequence: *GTAAA*KSRITSEGEYIPLDQIDINV. *Italic* characters denote linker region. (**C**-**D**) Surface staining on FLAG and HA tags to reveal the surface expression of eLIDAR receptors by transient transfection. Anti-FLAG-APC (abcam ab72569) was used to stain VEGFR1-ADAR2dd arm, and anti-HA-AF488 (BioLegend 901509) was used to stain VEGFR2-MCP arm. All cells are gated on positive co-transfection marker (BFP) expression. Negative control (no eLIDAR transfection) is shown in (C), and conditions with distinct linkers, Golgi traffic sequence and ER export sequence are shown in (D). Condition iv with Golgi traffic sequence + ER export sequence was chosen as optimal for further optimizations. (**E**) The effect of ER-retained catalytically inactive ADARdd (ER-deadADAR) on eLIDAR sensitivity to ligand revealed by running means plots. Condition iii with 50 ng ER-deadADAR was chosen as optimal since it yielded good separation of running means +/- ligand while not strongly hampering signal activation. (**F**) 2D fold activation plot of VEGF-sensing stably integrated eLIDAR cell line for finding optimal gating. To generate the 2D fold activation plot, we equally divided 200 bins across log scale on mCherry (marker on sensor) and TagBFP (marker on receptor components) for all cells in a specific condition (i.e. a specific VEGF concentration). Then for each of the 40,000 bins, we calculated the mean GFP (output) fluorescence and divided the value in the condition with saturating ligand concentration by the condition without ligand to generate a 2D matrix of GFP fold activation. Dashed blue box represents the optimal gating used in (G) and **Figure 2D**. (**G**) Dose-response curve of VEGF eLIDAR to recombinant VEGF-165 by fitting to a 5-parametric asymmetric logistic function. (**H**) 2D fold activation plot of IL-2 sensing eLIDAR stably integrated cell line for finding optimal gating. Dashed blue box represents the optimal gating used in (I) and **Figure 2G**. (**I**) Dose-response curve of IL-2 eLIDAR to recombinant IL-2 by fitting to a 4-parametric logistic function.

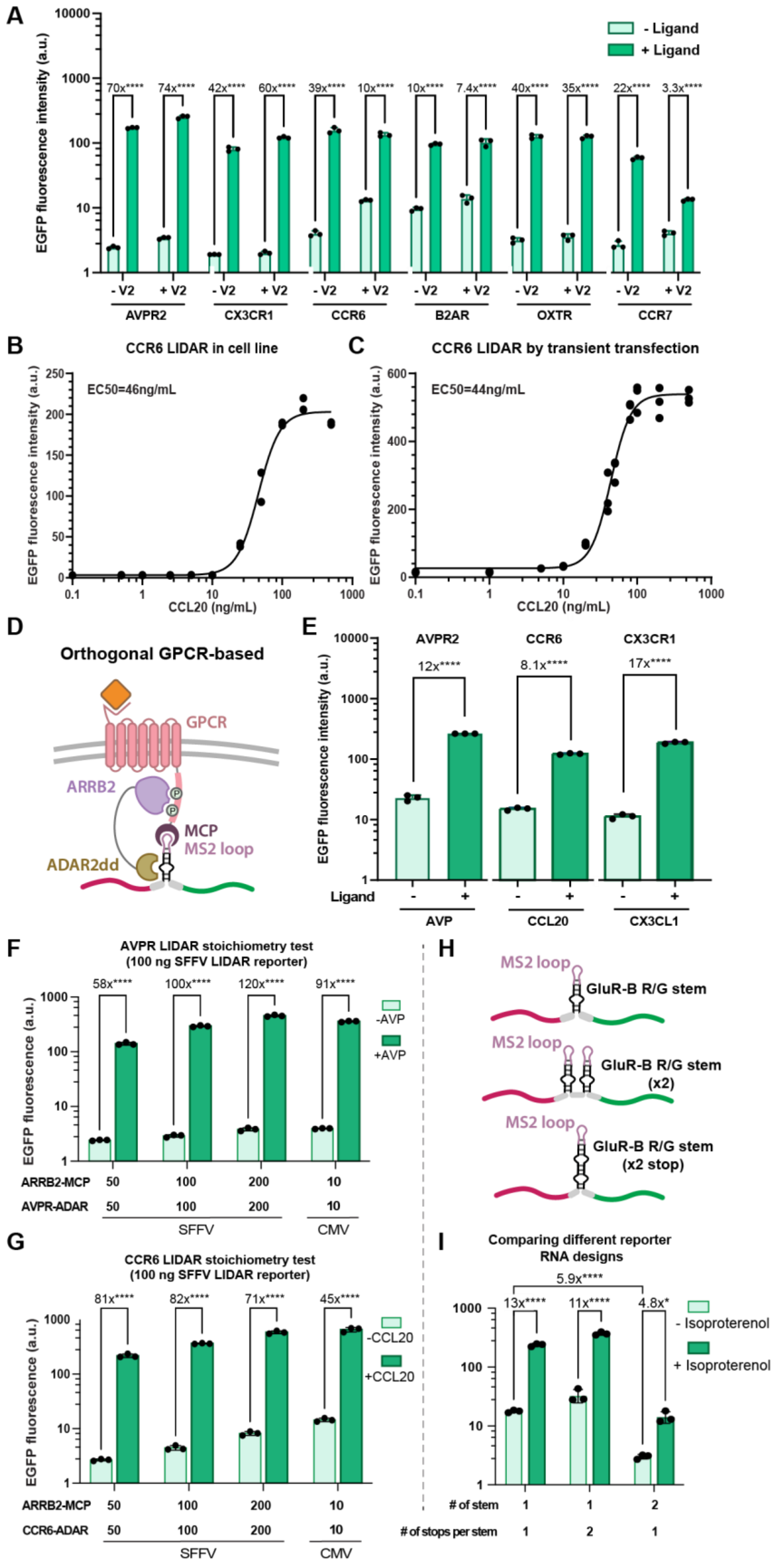
A
- Ligand
+ Ligand
EGFP fluorescence intensity (a.u.)
70x****
74x****
42x****
60x****
39x****
10x****
10x****
7.4x****
40x****
35x****
22x****
3.3x****
- V2
+ V2
AVPR2
CX3CR1
CCR6
B2AR
OXTR
CCR7
B
CCR6 LIDAR in cell line
EC50=46ng/mL
CCL20 (ng/mL)
C
CCR6 LIDAR by transient transfection
EC50=44ng/mL
CCL20 (ng/mL)
D
Orthogonal GPCR-based
GPCR
ARRB2
MCP
MS2 loop
ADAR2dd
E
AVPR2
CCR6
CX3CR1
12x****
8.1x****
17x****
Ligand
AVP
CCL20
CX3CL1
F
AVPR LIDAR stoichiometry test
(100 ng SFFV LIDAR reporter)
58x****
100x****
120x****
91x****
-AVP
+AVP
EGFP fluorescence (a.u.)
ARRB2-MCP
AVPR-ADAR
SFFV
CMV
G
CCR6 LIDAR stoichiometry test
(100 ng SFFV LIDAR reporter)
81x****
82x****
71x****
45x****
-CCL20
+CCL20
CCR6-ADAR
H
GluR-B R/G stem
GluR-B R/G stem (x2)
GluR-B R/G stem (x2 stop)
I
Comparing different reporter RNA designs
5.9x****
13x****
11x****
4.8x*
- Isoproterenol
+ Isoproterenol
### of stem
### of stops per stem

**Supplementary Fig. 3. Investigation and optimization of output levels in gLIDARs. (A)** Different gLIDARs have distinct performances depending on the composition of their C-terminal tails. gLIDAR with an extra V2 tail at their C-terminus exhibit lower baselines yet with lower activation levels. CCR7 is an exception. (**B-C**) Dose-response curves for CCR6-based gLIDAR **(B)** stably integrated and (**C**) transiently transfected. (**D**) Schematics showing the design of the orthogonal GPCR-based LIDAR (oLIDAR). (**E**) Data showing the performance of three different, individually transfected oLIDAR receptors. **(F-G)** The output levels of LIDAR are dependent on the stoichiometry of its components. This is shown for AVPR LIDAR **(F)** and CCR6 LIDAR **(G)**. All three components are driven by the SFFV promoter. All conditions in this experiment have 100ng of the reporter RNA compared to the 200ng normally used throughout this work. **(H)** Schematic showing the different reporter RNA designs tested to potentially lower the baseline in LIDAR. **(I)** Data showing the effect of the different reporter RNAs in **H** on the output levels of a B2AR-based LIDAR. B2AR LIDAR was chosen given its high baseline. The traditional reporter RNA still produces outputs with the best dynamic range, yet the 2 parallel stem loops (each containing a single stop codon) drastically decreases the baseline, which could be a desired feature for specific applications.

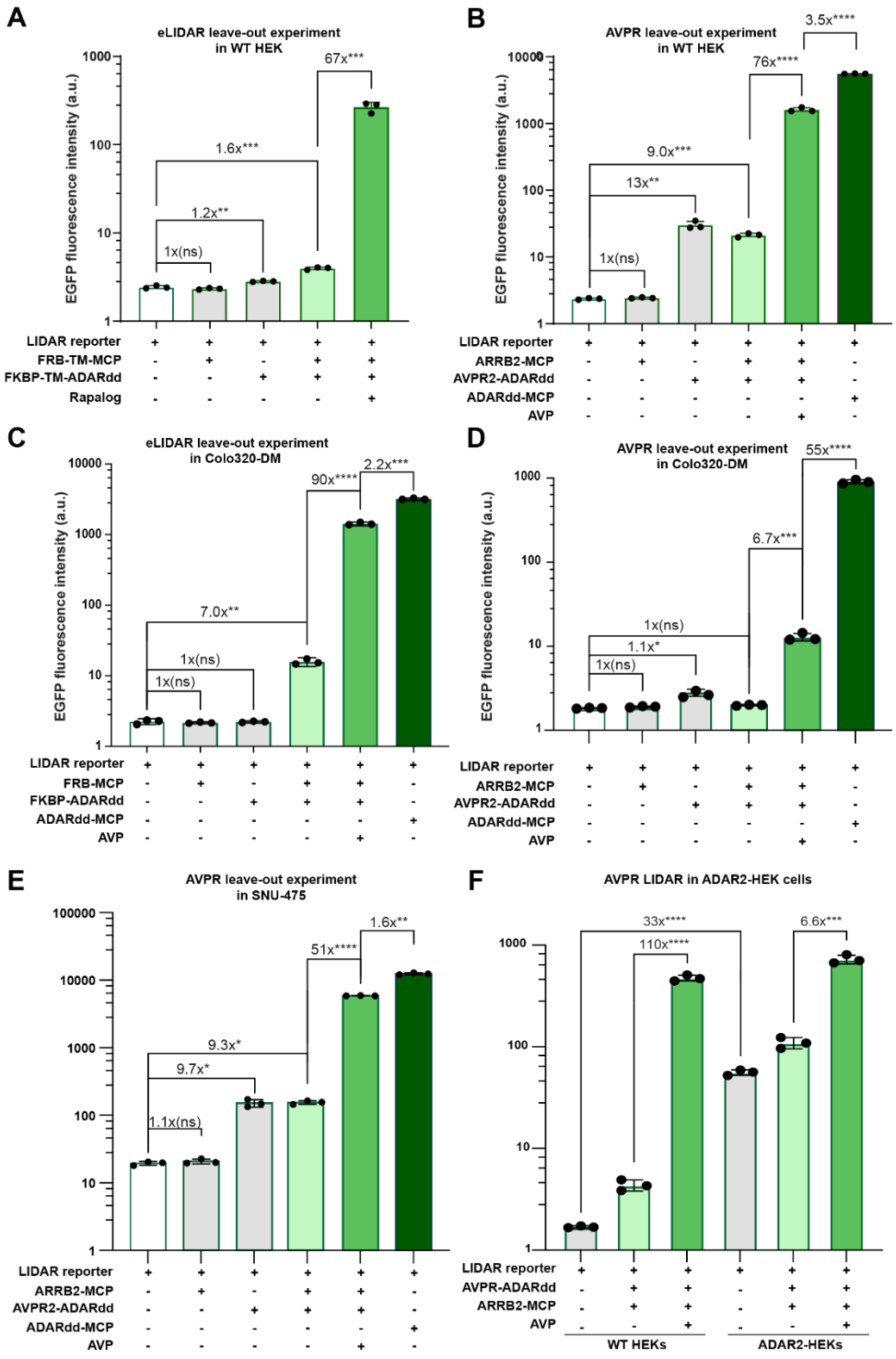


**Supplementary Fig. 4. Full set of control conditions for different LIDAR designs in different cell types. (A)** FKBP-FRB eLIDAR and **(B)** AVPR gLIDAR tested in WT HEK cells. Results show that baseline is mainly produced by the protein component containing ADAR2dd and not from the reporter RNA itself. Similarly, the efficiency of gLIDAR is comparable to that of a positive control where cytosolic ADAR2dd is fused to MCP. **(C)** FKBP-FRB eLIDAR and **(D)** AVPR gLIDAR tested in Colo320-DM. Like WT HEK cells, results show that LIDAR's baseline is mainly produced by the protein component containing ADAR2dd. Interestingly, AVPR performs worse in Colo320-DM than WT HEK293. **(E)** AVPR gLIDAR tested in SNU-475 cells. Consistent with LIDAR in other cells, high baseline levels are also attributed to the protein component containing the ADAR2dd. **(F)** AVPR gLIDAR tested in WT HEK293 overexpressing ADAR2 (cLSM12). High levels of ADAR2 increase the baseline of LIDAR by 33x. Interestingly, this increase in baseline is only prominent when LIDAR is delivered via DNA and not RNA (**Supplementary Fig. 10**). We believe this is because ADAR2 mainly localizes to the nucleus editing only plasmid-derived reporter RNAs.

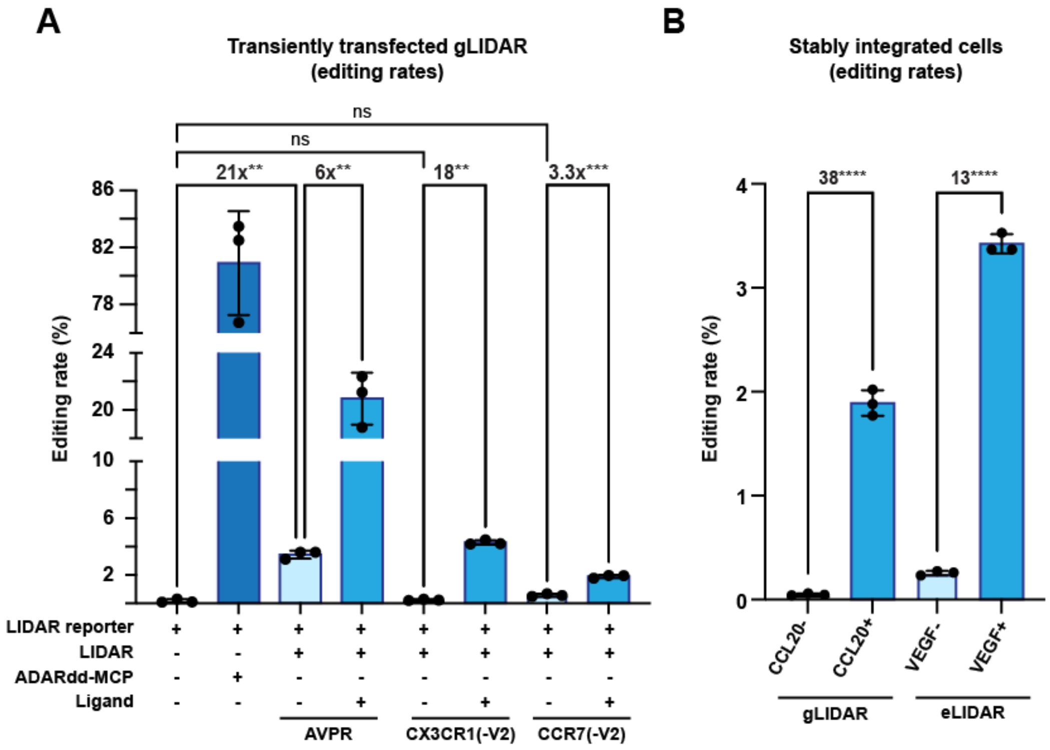


**Supplementary Fig. 5. Editing rates of different LIDAR architectures. (A)** Editing rates of different gLIDARs transiently transfected into WT HEK293. gLIDARs of different efficiencies were chosen to have a better understanding of how editing rates related to fluorescent output levels. As expected, AVPR(+V2) gLIDAR has a higher editing rate than CX3CR1(-V2) and CCR7(-V2), consistent with data in **Supplementary Fig. 3A**. These three also present lower rates compared to a positive control (cells transiently transfected with ADAR2dd-MCP and reporter RNA) and higher rates compared to a negative control (cells transiently transfected only with reporter RNA). **(B)** Editing rates of stably integrated LIDARs. Editing rate fold-changes are comparable to those observed for fluorescent output data.
Editing rate is subject to transfection efficiency, RNA extraction yields, amplification of the cDNA and/or sequencing.

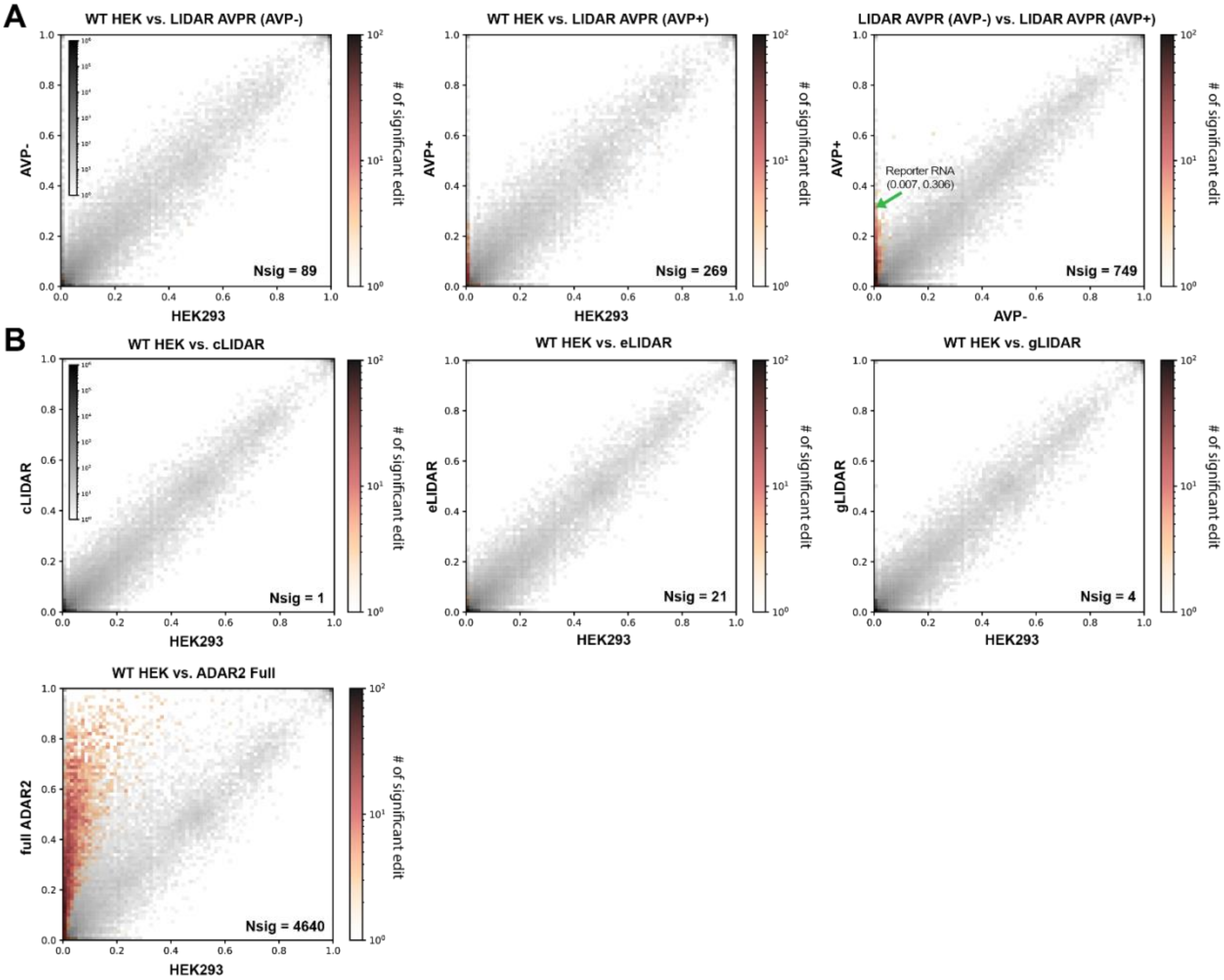


**Supplementary Fig. 6. Transcriptome-wide specificity profiles of RNA editing by LIDAR.** 2D histograms comparing the transcriptome-wide A>G editing yields observed in different LIDAR setups (y-axis) to the yields observed in a control sample (x-axis). Colored bins show the number of positions with significant A>G edits (p<0.01). Nsig is the total number of positions with a significant edit yield change. The analyses in A and B were performed separately. Different down sampling fractions were used and thus the resulting lists of sites are different. (**A)** RNA editing specificity of AVPR LIDAR in stable cell lines (cLSM02). Left: AVPR LIDAR without AVP is compared to WT HEK293. Middle: AVPR LIDAR treated with AVP compared to WT HEK293. Right: AVPR LIDAR treated with AVP compared to AVPR LIDAR without AVP. The red arrow points to the edited LIDAR reporter which shows an increase of almost 42x in editing rate (0.7% vs 30%). (**B)** RNA editing specificity of different LIDAR architectures. Cells were stably integrated only with the component that contains ADAR2dd (i.e., FKBP-tm-ADAR2dd, CCR6-ADAR2dd, FKBP-ADAR2dd) or full ADAR2 to account only for differences in editing yields due to exogeneous ADAR2. Top row: cLIDAR, eLIDAR and gLIDAR compared to the negative control (WT HEK) show very few significant off-target A>G edits. Bottom: ADAR2 compared to the negative control (WT HEK).

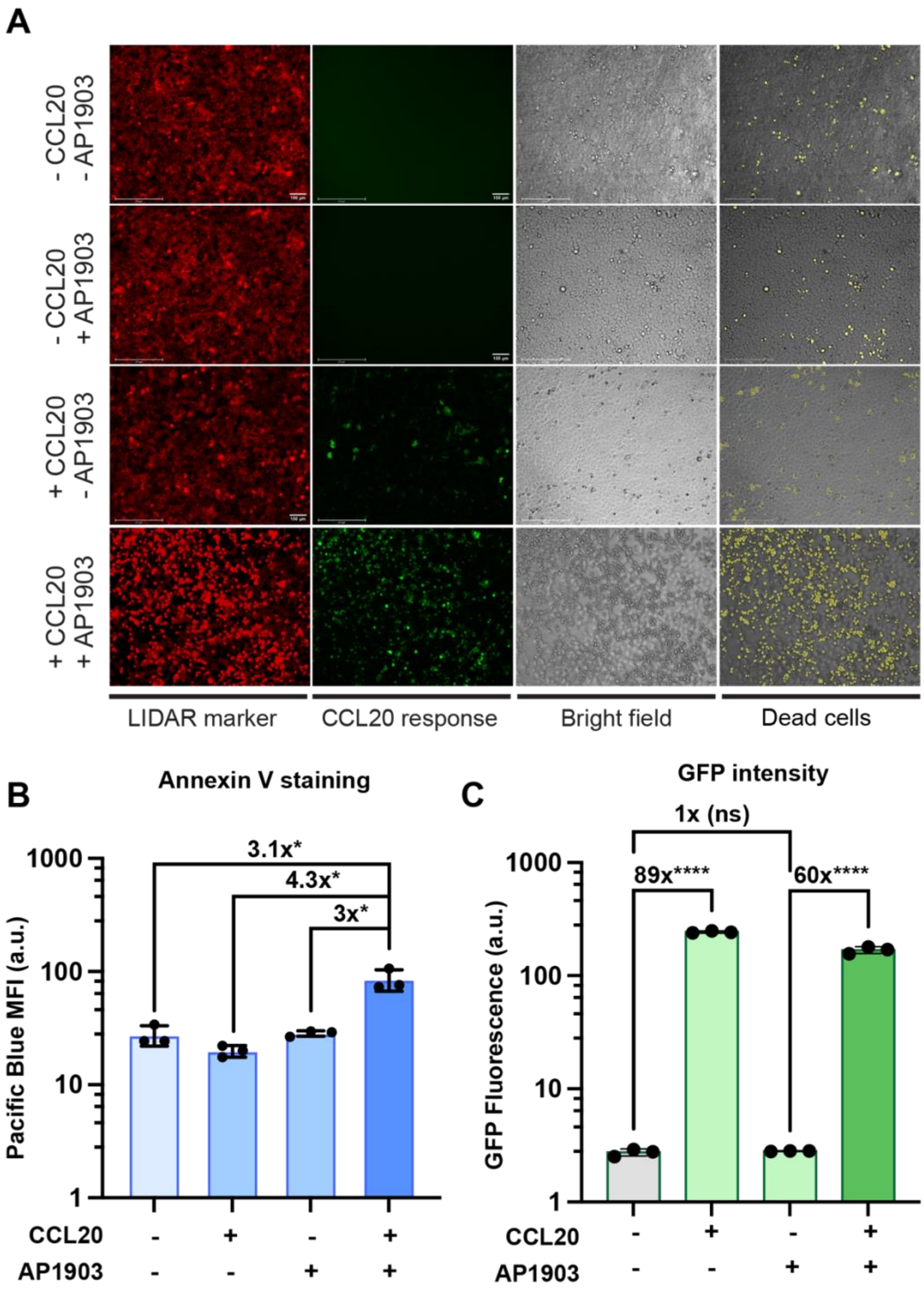


**Supplementary Fig. 7. Additional data supporting LIDAR-induced cell death. (A)** Images of each of the conditions for the LIDAR induced cell apoptosis experiments. Significant cell death (bottom panel) can be seen only for the condition where ligand (CCL20) and iCasp9 dimerization molecule (AP1903) were both administered to the cells. Healthy cells are observed for the rest of the conditions. Brightfield images with dead cells masked in yellow (left) are shown, emulating the staining of dead cells. These images were generated by computationally segmenting the death cells from brightfield images based on morphology. GFP positive cells are activated cells that express both iCasp9 and GFP as the LIDAR output. Marker mCherry fluorescence indicates the presence of the reporter RNA. **(B)** Annexin V staining on cells containing the same system as in **Fig. 3.** This is a different experiment from the one shown in Fig. 3 as staining and cell counting aren't compatible due to cell loss during staining. **(C)** GFP output data for the same experiment as in **B**. GFP is co-expressed with iCasp9 (separated by a 2A sequence) and can be used as a proxy for the amount of Casp9 being produced.

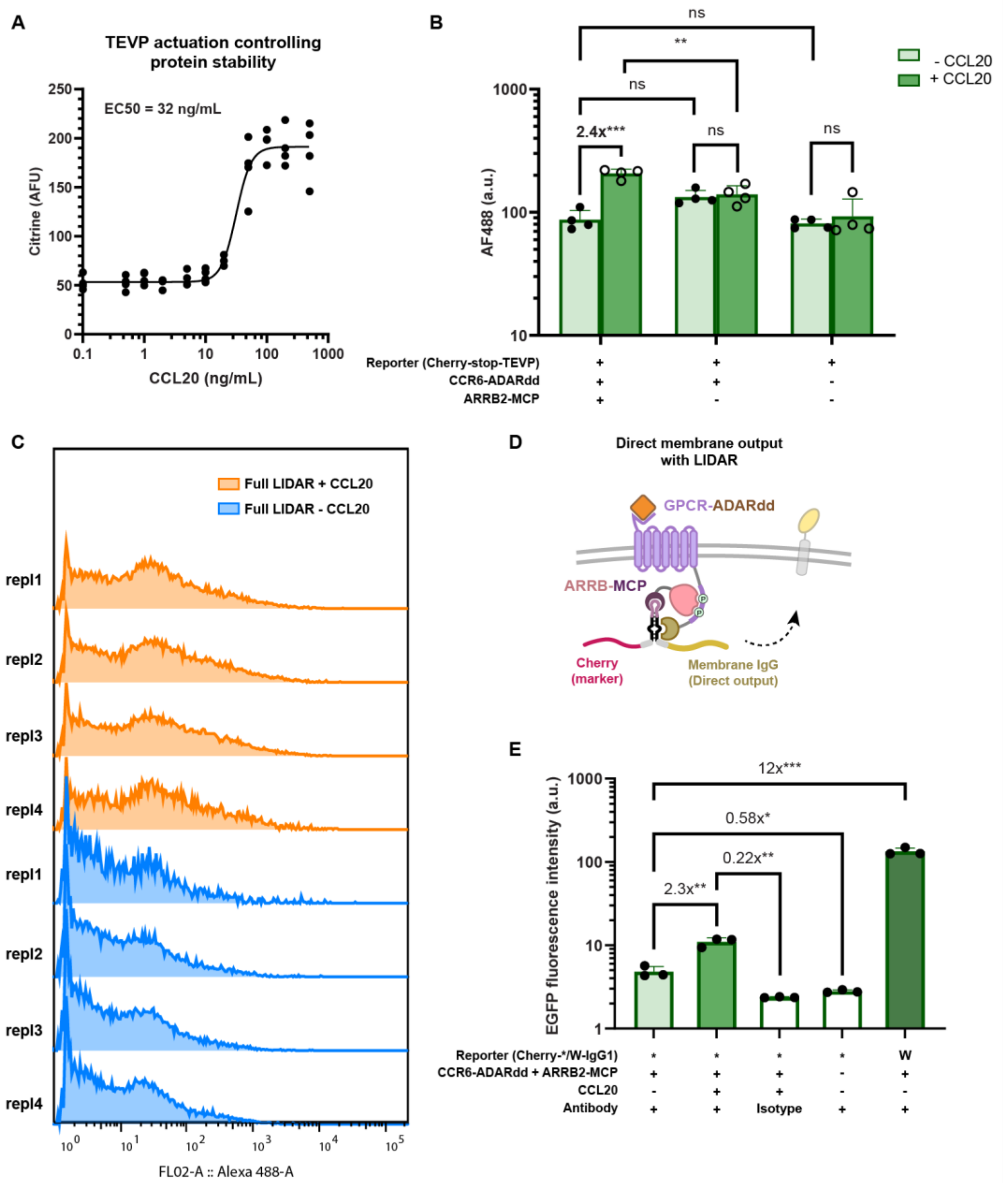


**Supplementary Fig. 8. Additional information regarding LIDAR with functional outputs.** (**A**) Dose-responsiveness curve of CCL20 gLIDAR with TEVP actuation controlling Citrine stability. Cells were gated on positive transfection marker population. (**B**) LIDAR-RELEASE with component drop-out negative controls. All conditions are co-transfected with ER-retained hIgG1. The first two columns are the same as **Fig. 3I**. All data are gated on all transfection marker positive cells as in **Fig. 3I**. (**C**) Histogram of AF488-A corresponds to data points in **Fig.3I**, gated on all transfection marker (mCherry+) positive cells. (**D**) Schematic of gLIDAR with direct output of a membrane-bound protein. (**E**) Benchmarking results of (D) using a membrane-bound hIgG1 as the direct output of LIDAR. This membrane-bound hIgG1 contains an Igκ leader signal peptide, and the extracellular and transmembrane domain is identical to the ER-retained hIgG1 used in **Fig. 3I** except for a lack of ER-retention motif. Cells are gated on all transfection marker positive cells. In the row of "Reporter (Cherry-*/W-IgG1)", "*" refers to the canonical reporter with a stop codon in the stem loop, and "W" refers to the positive control sensor where the stop codon is mutated to a

UGG tryptophan (W) codon. In the row of “Antibody”, “+” refers to staining with the specific antibody against human IgG, and “Isotype” refers to staining with an isotype control antibody. Refer to the **Methods** section for a detailed protocol on surface staining.

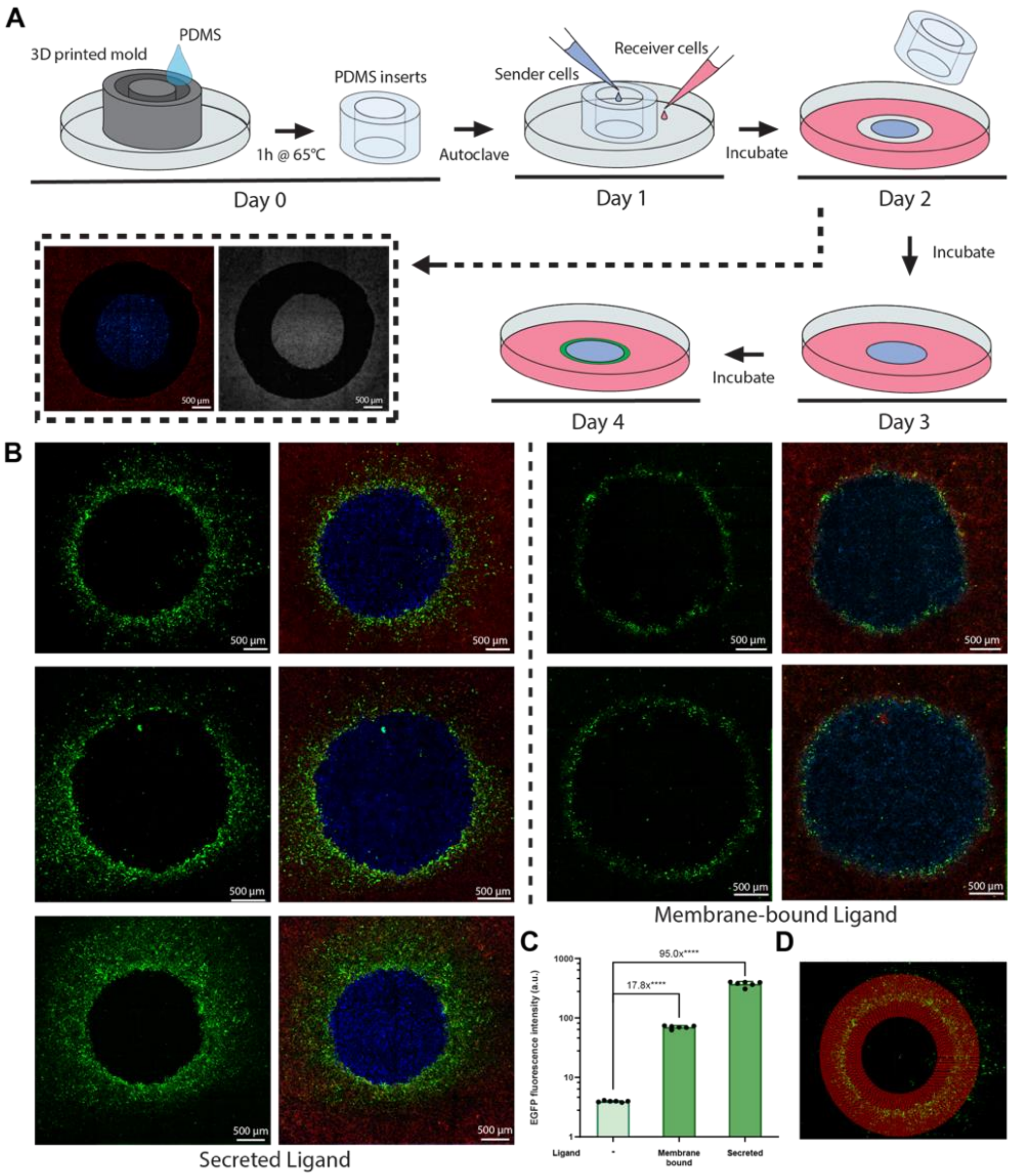


**Supplementary Fig. 9. Co-culture experiments for pattern formation.** (**A**) Fabrication of silicone inserts and seeding of sender and receiver cells for pattern formation experiments. A detailed explanation of the steps can be found in the Methods section. Cells are seeded in separate compartments using a PDMS insert with walls of approximately 500 µm. The day after seeding, the PDMS insert is removed to allow cells to grow and to come into contact with each other. After a couple of days, a pattern can be seen. (**B**) Additional patterns generated by the co-culture experiment. (**C**) Receiver cells (cLSM01) co-cultured with either secreted CCL20 or membrane-bound CCL20 cells. Data supports the difference in GFP intensities between the patterns generated by secreted and membrane-bound ligands. The data is from the same monoclonal cell line used for pattern formation and was gated on all mCherry positive cells. (**D**) Graphic representation of how the average intensity values were obtained to generate the intensity profiles for the patterns generated in the co-culture experiments. Each concentric red line represents a data point in the plots shown in **Fig. 4E**. For each pattern, 150 measurements (i.e., concentric red lines) were taken. Each of these measurements is the average value of 10,000 intensities. This means 10,000 data points are taken per red circle. A nonlinear background subtraction, as well as a linear enhancement of the brightness and contrast to aid with visualization, was performed on all images as detailed in the Methods section.

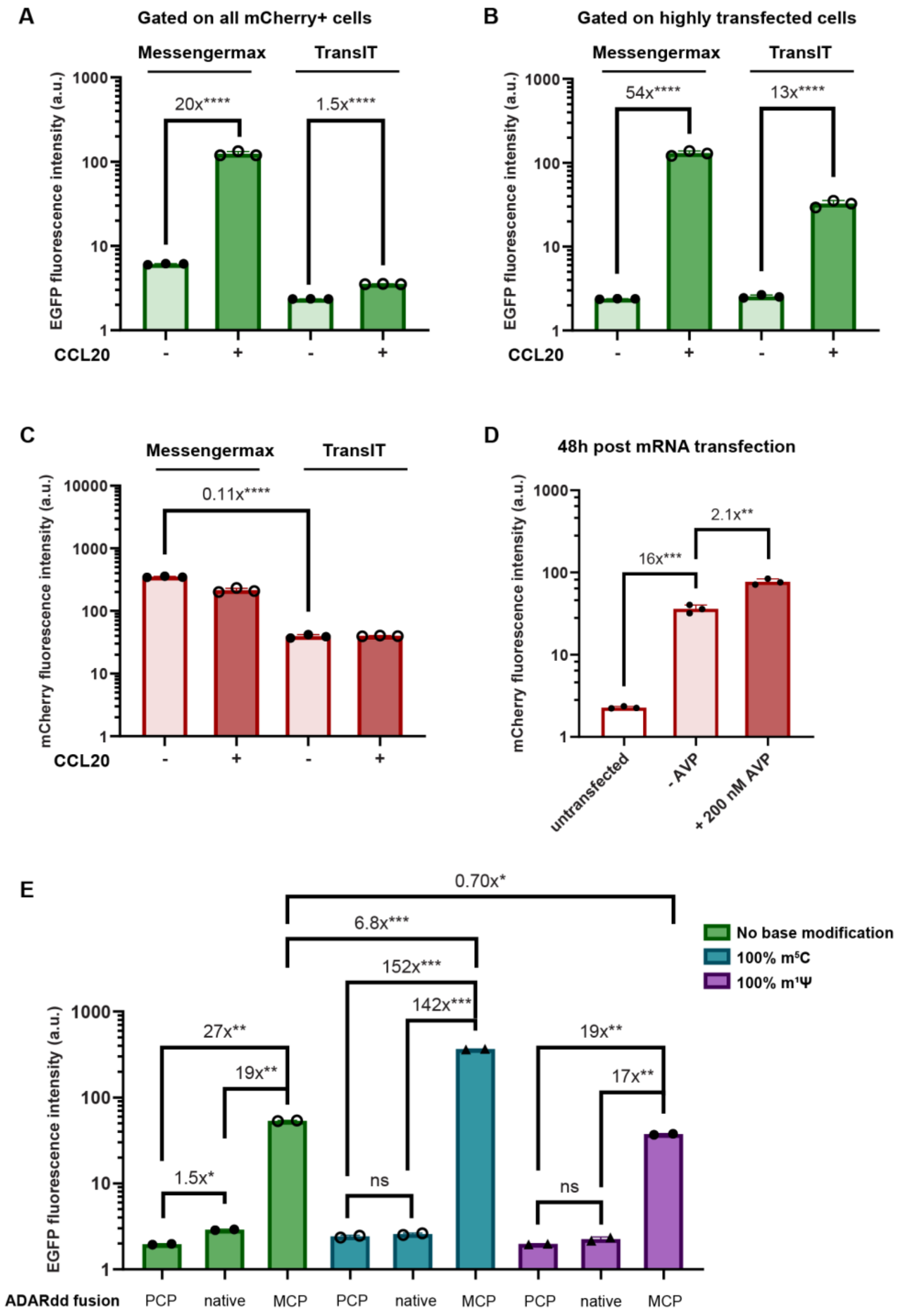


**Supplementary Fig. 10. Additional information of LIDAR with mRNA delivery.** (**A-C**) The effect of transfection reagents used on LIDAR fold activation and marker expression. (**A**) Comparison between Thermo MessengerMax and Mirus TransIT mRNA transfection reagent on EGFP output with CCL20 sensing gLIDAR, gated on all marker (mCherry) positive cells. CCL20 concentration: 100 ng/mL. (**B**) Data in (A) gated on highly transfected cells. (**C**) Overall (ungated) mCherry marker expression from the same experimental samples as in (A). (**D**) mCherry activation via AVP-sensing gLIDAR with Cre output assayed by flow 48 hours post mRNA transfection. (**E**) The effect of base modifications on editing capacity of LIDAR sensor mRNA. IVT LIDAR sensors with different base modifications are mRNA transfected into cell lines with stable expression of ADAR2dd-PCP fusion protein, ADAR2dd-MCP fusion protein or overexpression of native full-length ADAR2 (native).

**Supplementary Table 1.** List of plasmids (LIDAR).

| Plasmid | Link | Description | Source |
|---|---|---|---|
| EK0450 SFFV-mCherry-SGc(stem-1(3xMS2))cSG-EGFP | https://benchling.com/s/seq-2AXU87YTQeYS7XHKvTsS?m=slm-K2PJ08VrbHc5gDzkvC5a | LIDAR reporter RNA; mCherry as marker, EGFP as output | this work |
| EK0329 CMV-TO-FRB-MCP (FLP-IN) | https://benchling.com/s/seq-6OkIZha4M7lbxgLCfcTm?m=slm-poEy8QDMzH3X038Ibxi5 | FRB fused with MCP | this work |
| XZ135 CMV-TO-FKBP-ADARdd(QA) (FLP-IN) | https://benchling.com/s/seq-aHAXIPQVOsxv6PlAV7jd?m=slm-6LIGZ6TGAivHipUaKHiC | FKBP fused with ADAR2dd, with E488Q and T501A double mutation | this work |
| YH038 PGK-TO-FRB-tm-MCP | https://benchling.com/s/seq-9OibQKJveLgG6y44o5QA?m=slm-zHDb81Cv2kIlIcMweFlV | PGK promoter driven FRB fused with MCP, with a CD28 transmembrane domain in between | this work |
| YH039 PGK-TO-FKBP-tm-GS-XTEN-ADAR2(DD-E488Q-T501A) | https://benchling.com/s/seq-BmQBxHqOXwo4ruxEowAE?m=slm-JGIPwc9j1YXw04VVBbwl | PGK promoter driven FKBP fused with ADAR2dd (E488Q,T501A), with a CD28 transmembrane domain in between | this work |
| XZ024 CMV-TO-bArrestin2-MCP (FLP-IN) | https://benchling.com/s/seq-bh8Bj9ZK0csAjBV4M06E?m=slm-aSoStcNsBz2OMFmAh2Bs | ARRB2-tdMCP with a short linker | this work |
| XZ030 CMV-TO-AVPR2-ADAR2(DD-E488Q-T501A) (FLP-IN) | https://benchling.com/s/seq-sUeUvY2498uPF7HQmyHZ?m=slm-cVKrOE6agVNe2tXxYmw8 | AVPR2-V2-ADAR2dd(E488Q, T501A) with a short 4 amino acid linker and extra V2 tail | this work |
| XZ362 EF1a-VEGFR2sp-HA-VEGFR2(Ecto+TM)-TS-tdMCP-export | https://benchling.com/s/seq-uuKx8pqeH45Ve0X789ed?m=slm-hv1ylRKcEXrCfJ3Y9xt0 | Optimized VEGF-sensing eLIDAR with VEGFR2 ecto+TM domains fused with tdMCP, with Golgi traffic sequence and ER export sequence | this work |
| XZ363 EF1a-VEGFR1sp-3xFlag-VEGFR1(ecto+TM)-TS-ADAR2dd(QA)-export | https://benchling.com/s/seq-A9v8ZRSlMBEylC3tKkKz?m=slm-fSj76YGBHbaPLC5xpnAM | Optimized VEGF-sensing eLIDAR with VEGFR1 ecto+TM domains fused with ADAR2dd, with Golgi traffic sequence and ER export sequence | this work |
| YH001 CMV-TO-tm-GS+XTEN-ADAR2(DD-E396A)-RXR (FLP-IN) | https://benchling.com/s/seq-EvZcshwi3RXdhuSTRM5P?m=slm-1jvoGRxGKjO25UkMEDJu | ER-retained catalytically inactivae ADAR2dd (E396A mutation) | this work |
| EK0479 CMV-TO-VEGF-A(165) (FLP-IN) | https://benchling.com/s/seq-Ty39QAt08sczIkuVmlEY?m=slm-YKbSTsdBl7y8pIyx3JJU | Secreted human VEGF-A | this work |
| XZ377 (PB) EF1a-VEGFR1sp-3xFlag-VEGFR1(ecto+TM)-TS-ADAR2dd(QA)-export-IRESv0- | https://benchling.com/s/seq-IJc8X8Mp4fhLXZAqjJBh?m=slm-qTiKEf7qGQFLc7a987iR | PiggyBac donor plasmid with optimized VEGF-sensing eLIDAR receptor components separated by EMCV IRES, Puromycin resistance and BFP as marker | this work |

| | | | |
|---|---|---|---|
| VEGFR2sp-HA-VEGFR2(Ecto+TM)-TS-tdMCP-export (SV40-PuroR-BFP) | | | |
| XZ375 EF1a-IL2Rbsp-3xFLAG-IL2Rb(ECD+TM)-TS-ADAR2dd-export | https://benchling.com/s/seq-58dbLbf352drOPNoRZ0S?m=slm-A5DkvgbnddCbGL1YmuHl | Optimized IL-2-sensing eLIDAR with IL-2 beta chain ecto+TM domains fused with ADAR2dd, with Golgi traffic sequence and ER export sequence | this work |
| CMVTO-hIL2 | https://benchling.com/s/seq-0b77QpEMn0YfV7Tjutcb?m=slm-OhE8leQHhmElfFtS5Ma2 | secreted human IL-2 | this work |
| XZ393 (PB) EF1a-IL2Rbsp-FLAG-IL2Rb(ECD+TM)-TS-ADAR2dd-export-IRESv0-IL2Rgsp-3xHA-IL2Rg(ECD+TM)-TS-tdMCP-export (SV40-PuroR-BFP) | https://benchling.com/s/seq-qDgUYqS5oOlpIY89T3YS?m=slm-sqB6atLTjgNDwCKDA4pN | PiggyBac donor plasmid with optimized IL-2-sensing eLIDAR receptor components separated by EMCV IRES, Puromycin resistance and BFP as marker | this work |
| XZ374 EF1a-IL2Rgsp-3xHA-IL2Rg(ECD+TM)-TS-tdMCP-export | https://benchling.com/s/seq-Tb1kYDuvGa8TXLQZtjz0?m=slm-UhjfRkQb8CmJYkomoWRI | Optimized IL-2-sensing eLIDAR with IL-2 gamma chain ecto+TM domains fused with tdMCP, with Golgi traffic sequence and ER export sequence | this work |
| LSM104 CMV-TO-AVPR2-(noExtraV2)-ADAR2(DD-E488Q-T501A) (FLP-IN) | https://benchling.com/s/seq-LHTVjwaqizhT57BDyA1Q?m=slm-w9ypBuXnFG4kQSMqGE7A | CCR6-ADAR2dd(E488Q, T501A) with a short 4 amino acid linker and no extra V2 tail | this work |
| XZ038 CMV-TO-IGF1R-ADAR2(DD-E488Q-T501A) (FLP-IN) | https://benchling.com/s/seq-gES0zs4kWvHJOJQcNFrH?m=slm-U1SS9WOgC7LSc2EFSrgE | IGF1R-ADAR2dd(E488Q, T501A) | this work |
| XZ039 CMV-TO-Shc1-MCP (FLP-IN) | https://benchling.com/s/seq-IstmWwXD688xj1rCMKv4?m=slm-9pnqOIKH5rPPt2t5ttkL | Shc1-tdMCP | this work |
| EK0456 CMV-TO-IL2RBec-tm-ADAR2(DD-E488Q) (FLP-IN) | https://benchling.com/s/seq-P5rORCDitiq2RwBfRIdt?m=slm-IEAEyyExZwKWHsapWGeM | IL-2 sensing extracellular LIDAR: extracellular domain of IL-2Rβ fused with CD28-TM and ADAR2dd(E488Q) | this work |
| EK0457 CMV-TO-IL2RGec-tm-MCP (FLP-IN) | https://benchling.com/s/seq-1YcJk99EnjDHcECRfVDQ?m=slm-qwWLpLOBATghRzIZaIQh | IL-2 sensing extracellular LIDAR: extracellular domain of IL-2Rγ fused with CD28-TM and tdMCP | this work |
| CMVTO-hIL2 (AV) | https://benchling.com/s/seq-vaR2QlL46aD9ZZsCObs4/edit | Plasmid expressing secreted IL-2 | from lab members |
| XZ031 CMV-TO-CCR6-ADAR2dd(E488Q, T501A) | https://benchling.com/s/seq-Cb2pE4RYqT4tAgrntE5j?m=slm-Dxgj6727cGxsvQJWulT2 | CCR6-ADAR2dd(E488Q, T501A) with a short 4 amino acid linker and no V2 tail | this work |
| XZ032 CMV-TO-CCR6-V2-ADAR2dd(E488Q, T501A) | https://benchling.com/s/seq-LWA0E18utbuUodkltJbe?m=slm-FGFVgRBX13uMqBl8VPP5 | CCR6-V2-ADAR2dd(E488Q, T501A) with a short 4 amino acid linker and V2 tail | this work |

| | | | |
|---|---|---|---|
| LSM044-3 CMV-TO-CCR7-ADAR2dd(E488Q, T501A) (FLP-IN) | https://benchling.com/s/seq-HYqL7ToFC8zhjxMaN8Lk?m=slm-u7jNDGeknA806jFId71Q | CCR7-ADAR2dd(E488Q, T501A) with a short 4 amino acid linker and no V2 tail | this work |
| LSM044-2 CMV-TO-CCR7-V2-ADAR2dd(E488Q, T501A) (FLP-IN) | https://benchling.com/s/seq-Axp9V6ibGqMadjDF2PYB?m=slm-9NLgHPmdQLbKae7RjFAi | CCR7-V2-ADAR2dd(E488Q, T501A) with a short 4 amino acid linker and V2 tail | this work |
| LSM105 CMV-TO-CX3CR1-(noV2)-ADAR2dd(E488Q, T501A) (FLP-IN) | https://benchling.com/s/seq-qkDczDmlDyEeNseYdwv2?m=slm-0oq7OjcW5D4UrrkdwCXl | CX3CR1-ADAR2dd(E488Q, T501A) with a short 4 amino acid linker and no V2 tail | this work |
| LSM068 CMV-TO-CX3CR1-ADAR2dd(E488Q, T501A) (FLP-IN) | https://benchling.com/s/seq-7zIaSa8skxHXSXpPXvO0?m=slm-wO4pe3xIQMokgJq8j9BV | CX3CR1-V2-ADAR2dd(E488Q, T501A) with a short 4 amino acid linker and V2 tail | this work |
| XZ113 CMV-TO-signal-hOXTR-ADAR2dd(DD-E488Q-T501A) (FLP-IN) | https://benchling.com/s/seq-0idrTEpYZyboXpNbpwYT?m=slm-dEhAGxZHHrQA3wwCfGE3 | human oxytocin receptor-ADAR2dd(E488Q, T501A), no V2 tail | this work |
| XZ105 CMV-TO-signal-hOXTR-V2-ADAR2dd(DD-E488Q-T501A) (FLP-IN) | https://benchling.com/s/seq-0gM0nGZFq2zjTQVUqfpJ?m=slm-9VmLGgYrAytY8uC7zip5 | human oxytocin receptor-V2-ADAR2dd(E488Q, T501A) | this work |
| XZ142 CMV-TO-signal-B2AR-ADAR2dd(DD-E488Q-T501A) (FLP-IN) | https://benchling.com/s/seq-iz9oiacqsPpxHItmHrpw?m=slm-R6R4EKeqkVZMGB6WWjtF | β-2 adrenergic receptor-ADAR2dd(E488Q, T501A), no V2 tail | this work |
| XZ110 CMV-TO-signal-B2AR-V2-ADAR2dd(DD-E488Q-T501A) (FLP-IN) | https://benchling.com/s/seq-jRk84N3P7zomZbclRweM?m=slm-dUDuZARfqh6FSjHX1pqJ | β-2 adrenergic receptor-V2-ADAR2dd(E488Q, T501A) | this work |
| LSM060 CMV-TO-AVPR2-(10aa-linker)-MCP (FLP-IN) | https://benchling.com/s/seq-0rwLMWudhTlMeI54wlgr?m=slm-kcqazNtp7rye0O5VTio1 | AVPR2-V2-tdMCP with a 10 amino acid linker and extra V2 tail | this work |
| LSM102 CMV-TO-CX3CR1-10aa-PCP (FLP-IN) | https://benchling.com/s/seq-pUpM5j0kmQBtXm3yRbHY?m=slm-cmG6yA3C0LebhHIifObE | CX3CR1-V2-PCP with a 10 amino acid linker and V2 tail | this work |
| LSM103 CMV-TO-CX3CR1-10aa-MCP (FLP-IN) | https://benchling.com/s/seq-ypBig5z7offv0JEAil5J?m=slm-hHIZh9XlcroVU2pAxYzN | CX3CR1-V2-tdMCP with a 10 amino acid linker and V2 tail | this work |
| LSM052 CMV-TO-CCR6-PCP (FLP-IN) | https://benchling.com/s/seq-29E6iIPDqqeKwiAUCFEV?m=slm-OSXPksR8bPHDCGHydSZY | CCR6-V2-PCP with a 4 amino acid linker and V2 tail | this work |
| LSM059 CMV-TO-bArrestin2-(31-linker)-ADAR2dd(E488Q, T501A) (FLP-IN) | https://benchling.com/s/seq-fwQowr4q4Z8c1gThqpv1?m=slm-3NPrPLokDv0J4xT1D658 | ARRB2-ADAR2dd(E488Q, T501A) with a long 31 amino acid linker | this work |

| | | | |
|---|---|---|---|
| LSM041-3 SFFV-mCherry-SGc(stem-1(3xPP7))cSG-EGFP | https://benchling.com/s/seq-cKvCf0Sf0crDNbI8aBsz?m=slm-jMGVoRASJSWPCD8Q26HD | LIDAR reporter RNA with PCP binding site instead of MCP; mCherry as marker, EGFP as output | this work |
| LSM053 SFFV-mCherry-SGc(stem-1(3xPP7))cSG-mtagBFP (FLP-IN) | https://benchling.com/s/seq-nNpyjHprmqSFptA3DPvB?m=slm-ce9wmZsVoyalKcQbZB5y | LIDAR reporter RNA with PCP binding site instead of MCP; mCherry as marker, mTagBFP as output | this work |
| APL0038 Ac5-mCherry-SGc(stem-1(3xMS2))cSG-EGFP | https://benchling.com/s/seq-vO8XLKeH7ToMKPTNdx7R?m=slm-0WUYciUajw0SA7SzAJJp | AC5 promoter driven LIDAR reporter RNA for drosophila cell expression (mCherry as marker, EGFP as output) | this work |
| APL0040 Ac5-FKBP-ADARdd | https://benchling.com/s/seq-jDtUGVRxvyj9D4E0GtR3?m=slm-TtGbFPFI2y81JYFaj2DD | AC5 promoter driven FRBP-ADAR2dd for drosophila cell expression | this work |
| APL0041 Ac5-FRB-MCP | https://benchling.com/s/seq-YGbXEwWNp6FRR7ZV0c1Z?m=slm-RY4fDhkpLwrfsTxn8gcr | AC5 promoter driven FRB-MCP for drosophila cell expression | this work |
| LSM043_2 (PB)SFFV-LIDARSensor(mCherry->Casp9-2A-GFP)-hygro | https://benchling.com/s/seq-esvMr2zCzxlmb0wmksiI?m=slm-31nTWQUmdL7sfb3mIbBg | LIDAR reporter RNA; mCherry as marker, iCasp-P2A-EGFP as output | this work |
| EK0461 SFFV-mCherry-SGc(stem-1(3xMS2))cSG-TEVp (FLP-IN) | https://benchling.com/s/seq-4irvxViGMPV731Xn1u4B?m=slm-1lVhMceFf7ciMRB2bH8P | LIDAR reporter RNA with TEVP as the output | this work |
| AV324_CMVTO-hIgG1-B2AD-tevs-KKMP | https://benchling.com/s/seq-9fWRnOIkTqDcP2POqomL?m=slm-1MPE3HINuFmKFsT1R4SJ | RELEASE construct for hIgG1 secretion | Vlahos et al. (2022) [5] |
| CMV-TO-TEVP (FLP-IN) | https://benchling.com/s/seq-F2QHm3L0ApHSuTrmlKd6?m=slm-1wZQSyJUljv4atwdSKIw | CMV-TO-TEVP | Vlahos et al. (2022) [5] |
| PB-PGK-Cit-tevs-DHFR | https://www.addgene.org/116040/ | Citrine-tev cutting site-degron | Gao et al. (2018) [6] |
| XZ400 SFFV-mCherry-SGc(stem-1(3xMS2))cSG-TEVP | https://benchling.com/s/seq-JjXSoNTmFcjKrHCszCQM?m=slm-r7r8gX9iR7EuzVdWPyd5 | LIDAR reporter RNA; mCherry as marker, TEVP as output | this work |
| XZ392 SFFV-secNLuc-SGc(stem-1(3xMS2))cSG(tight)-ssSEAP | https://benchling.com/s/seq-NCmc96wUNdMKjd6ItnWd?m=slm-jPBiTguyZ1yWNwp50hJ8 | LIDAR reporter RNA; secreted NanoLuc Luciferace as marker, SEAP as output | this work |
| AV282_CMVTO-SEAP-26SFur-B2AD-tevs-RXR6.2 | https://benchling.com/s/seq-JWQzlbcctCK53n7vDHPE?m=slm-BsNIE2JMMVZU234K18fw | ER-retained SEAP | Vlahos et al. (2022) [5] |

| | | | |
|---|---|---|---|
| CMVTO-secNluc | https://benchling.com/s/seq-k1viRrCJ7BvIXV2zXnvh?m=slm-1lBJ9X4pFJT5pt5zhhFK | secreted NanoLuc Luciferase | Vlahos et al. (2022) [5] |
| XZ034 CMV-TO-CCL20 | https://benchling.com/s/seq-ohAcW1bjwheIzwIzbb3J?m=slm-9YzSBcvKign09VKTZU6K | plasmid expressing secreted soluble CCL20 | this work |
| XZ033 CMV-TO-CCL20-pre-mGRASP | https://benchling.com/s/seq-meSeMrTnHb2DxwhM3O2D?m=slm-njLpyEn7rDkgAGS3xPJj | plasmid expressing membrane-bound CCL20 by fusing with pre-mGRASP | this work |
| XZ079 (PB)EF1a-CCR6-ADARdd-P2A-PuroR-IRES-ARRB2-MCP | https://benchling.com/s/seq-3hIhl82C7vf7KdrWZVrF?m=slm-eI2fm6Gfrh7oZ38LuYtH | PiggyBac donor plasmid with CCR6-V2-ADAR2dd(E488Q,T501A) and ARRB2-tdMCP separated by EMCV IRES | this work |
| XZ054 (PB)SFFV-LIDARsensor (hygR) | https://benchling.com/s/seq-CEoB39JotlEjpurP02TJ?m=slm-dLQpKc3vOFFWPvrXYCDM | PiggyBac donor plasmid with SFFV driven LIDAR reporter RNA (mCherry as marker, EGFP as output) for stable integration | this work |
| XZ073 (IVT) CMV-TO-T7-LIDARsensor-hybridUTR3 | https://benchling.com/s/seq-K1euM67vxduUsiTwzMAH?m=slm-OlPsGDFUzFqCJ6NpoLwu | IVT template for LIDAR reporter mRNA; mCherry as marker, EGFP as output | this work |
| XZ088 (IVT) CMV-TO-T7-CCR6-V2-ADAR2dd-hybridUTR3 | https://benchling.com/s/seq-40D6RH0zn2Gko5GElQCv?m=slm-ES47gUS0MW0B8KXczWX9 | IVT template for CCR6-V2-ADAR2dd(E488Q, T501A) | this work |
| XZ089 (IVT) CMV-TO-T7-ARRB2-MCP-hybridUTR3 | https://benchling.com/s/seq-xf0ht9hrkW50VTzN2bJe?m=slm-ONtrbBkNsyKbavxOA2QT | IVT template for ARRB2-tdMCP | this work |
| XZ387 (IVT) CMV-TO-T7-AVPR2-V2-ADAR2dd-hybridUTR3 | https://benchling.com/s/seq-BwQ4kV5Tvc3HqCawxMXm?m=slm-dAjPQsnsjuA3q0hiRE5L | IVT template for AVPR2-V2-ADAR2dd(E488Q, T501A) | this work |
| XZ388 (IVT) CMV-TO-T7-EGFP-SGc(stem-1(3xMS2))cSG-Cre-hybridUTR3 | https://benchling.com/s/seq-LLTs1HH40jfeoMtFc7T8?m=slm-Y0o2jHLyncgPTnQcPADJ | IVT template for LIDAR reporter mRNA; EGFP as marker, Cre recombinase as output | this work |
| XZ390 (IVT) CMV-TO-T7-mCherry-SGc(stem-1(3xMS2))cSG-rTetR-NZF-NLS-hybridUTR3 | https://benchling.com/s/seq-kRVEaM1Hx2dYebhFWPPD?m=slm-hDiSJW8LId6sROelKyOR | IVT template for LIDAR reporter mRNA; mCherry as marker, rtetR-NZF as output | this work |
| XZ136 SFFV-mCherry-SGc(stem-1(2xMS2))cSG-EGFP (FLP-IN) | https://benchling.com/s/seq-u4O8nSvkmS1VKmUxL4yy?m=slm-bJ0GzBZEPlZbqc7GO0Jv | LIDAR reporter RNA with 2x MS2 loop | this work |
| EK0480 SFFV-mCherry-SGc(stem-1(1xMS2))cSG-EGFP (FLP-IN) | https://benchling.com/s/seq-rNB2DjK7K2dJMIYwJ2DT?m=slm-96eL2ph4Mt9zPfXmdAwJ | LIDAR reporter RNA with 1x MS2 loop | this work |

| | | | |
|---|---|---|---|
| XZ029 CMV-TO-AVPR2-ADAR2(DD-E488Q) (FLP-IN) | https://benchling.com/s/seq-IXB0xL9BeDXIE8WsQY9K?m=slm-c3stU6t1vDhm3hNEhdmt | AVPR2-V2-ADAR2dd(E488Q) with a short 4 amino acid linker and extra V2 tail | this work |
| EK0330 CMV-TO-FKBP-ADAR2(DD-E488Q) (FLP-IN) | https://benchling.com/s/seq-fIBi5nBt3UN26sjA0apq?m=slm-arji01P2X69UFI5cbino | FKBP fused with ADAR2dd, with E488Q single mutation | this work |
| XZ157 CMV-TO-AVPR2-10L-ADAR2dd(QA) (FLP-IN) | https://benchling.com/s/seq-4iNpWMEBt3j0OpznPKtl?m=slm-o4KhodlBrtk2bFjefQH9 | AVPR2-V2-ADAR2dd(E488Q, T501A) with a short 10 amino acid linker and extra V2 tail | this work |
| XZ156 CMV-TO-AVPR2-GSXTEN-ADAR2dd(QA) (FLP-IN) | https://benchling.com/s/seq-4jm07A8wBOObLwcccZxl?m=slm-EZx0FTPaxbPPo6PTdpuZ | AVPR2-V2-ADAR2dd(E488Q, T501A) with a short 31 amino acid linker and extra V2 tail | this work |
| XZ158 CMV-TO-AVPR2-45ECL-ADAR2dd(QA) (FLP-IN) | https://benchling.com/s/seq-NO634GGdLN70zbv4qjEU?m=slm-xvd0XfDLxkfVHupmUSSj | AVPR2-V2-ADAR2dd(E488Q, T501A) with a short 47 amino acid linker and extra V2 tail | this work |
| LSM042 (PB)EF1a-AVPR-ADARdd-P2A-PuroR-IRES-ARRB2-MCP | https://benchling.com/s/seq-9HQ1AT6SsmFb6tzjUxne?m=slm-3avkCBWnNHt903DhgS6v | PiggyBac donor plasmid AVPR2-V2-ADAR2dd(E488Q,T501A) and ARRB2-tdMCP separated by EMCV IRES | this work |
| LSM061 (PB)SFFV-LIDARsensor(mcherry-BFP) (hygR) | https://benchling.com/s/seq-q8G1CIjls4Ajf7wG7vh8?m=slm-kCOlBe1kTC76JYbgcXfg | PiggyBac donor plasmid with SFFV driven LIDAR reporter RNA (mCherry as marker, TagBFP as output) | this work |
| LSM108 CMV-TO-AVPR-TEVcs-(noLinker)-tTA2 (FLP-IN) | https://benchling.com/s/seq-CzkTQWqPcTnOMPhQPrYT?m=slm-ihvG3xteZ0GS1wWobz9m | AVPR2-TEVcs-tTA for TANGO assay | Kroeze et al. (2015) [7] |
| pCDNA3.1(+)-CMV-bArrestin2-TEV | https://www.addgene.org/107245/ | Beta-arrestin2-TEV fusion protein for TANGO assay | Kroeze et al. (2015) [7] |
| XZ087 SFFV-mCherry-sensor(tight)-hIgG1-TM | https://benchling.com/s/seq-OMDamWVNeTRZDU0ZDaSo?m=slm-3agsb2uhmUA956CHpd0S | LIDAR reporter RNA with a membrane-bound human IgG as output | this work |
| NK90-pTRE3G-EGFP-stop (PB) | https://benchling.com/s/seq-cNlPsIUIoCJEbFzVYr8A?m=slm-4fmuJG1mNkhony5COpWI | PiggyBac donor plasmid for TRE3G driven EGFP | from lab members |
| XZ098 SFFV-mCherry-LIDARsensor(W)-hIgG1-TM | https://benchling.com/s/seq-8oYBqAcwJgyyNm4XcT0E?m=slm-HNZbLzqGDpZLJdYPoD9A | Positive control LIDAR reporter RNA (UAG stop codon replaced with UGG) with a membrane-bound human IgG as output | this work |
| XZ093 SFFV-sensor-IRESv0-LIDARst(IRESv0-AVPR2) | https://benchling.com/s/seq-khd9SJnIvHG9xrZfhA9i?m=slm-Wnc0GrDuXTvkbKsQPZQa | LIDAR all components on single transcrip (as shown in FIgure 4A), using EMCV IRES to separate each component | this work |

| | | | |
|---|---|---|---|
| YX022 (PB)EF1a-LIDARsensor (hygR) | https://benchling.com/s/seq-55bX1njYZf6leQYbwoNG?m=slm-lHbYA4lfOv3B4zPOvPGf | EF1a-driven LIDAR reporter RNA | this work |
| LSM101 CMV-TO-ADAR2dd(E488Q,T501A)-MCP | https://benchling.com/s/seq-fAv4nr8uIkD9tyQi54mq?m=slm-U9LBYuM2Xo4BmxGEevQ1 | ADAR2dd(E488Q,T501A)-MCP (positive control that fuses ADAR2dd and MCP on the same protein) | this work |
| XZ076 (PB) CMV-mCCL20 (PuroR-BFP) | https://benchling.com/s/seq-tGEOkY31mk9Nhd5QSRVg?m=slm-ByM8NLYwAY06KBcB9Lse | PiggyBac donor plasmid, membrane-bound CCL20 | this work |
| XZ091 (PB) CMV-sCCL20 (PuroR-BFP) | https://benchling.com/s/seq-YOrLHCeBmRzQpZSL9N6R?m=slm-t7xWlGQ6zvhjz6arFxH2 | PiggyBac donor plasmid, secreted CCL20 | this work |
| XZ079 (PB)EF1a-CCR6-ADARdd-P2A-PuroR-IRES-ARRB2-MCP | https://benchling.com/s/seq-gdcaVPe6CDjsYw77dIkZ?m=slm-M9ggkDupVcl8VqQXuRSw | PiggyBac donor plasmid with CCR6-V2-ADAR2dd(E488Q,T501A) and ARRB2-tdMCP separated by EMCV IRES | this work |
| XZ344 EF1a-VEGFR2sp-HA-VEGFR2(Ecto+TM)-SL-tdMCP | https://benchling.com/s/seq-2E15cKyQPg790qrgZeOs?m=slm-bWiFxWKZ7I0Z63vw4XKL | VEGF-sensing eLIDAR with VEGFR2 ecto+TM domains fused with tdMCP with a short GS linker in between | this work |
| XZ345 EF1a-VEGFR1sp-3xFlag-VEGFR1(ecto+TM)-SL-ADAR2dd(QA) | https://benchling.com/s/seq-xy92Z0XQVHEzp0dH8ibq?m=slm-yvL3m1Nooqiq4DLGYE1I | VEGF-sensing eLIDAR with VEGFR1 ecto+TM domains fused with ADAR2dd with a short GS linker in between | this work |
| XZ358 EF1a-VEGFR2sp-HA-VEGFR2(Ecto+TM)-SL-tdMCP-export | https://benchling.com/s/seq-EToXIsmN7r7stxyYevJl?m=slm-KqGMBmSfkK5pv9HX37dq | VEGF-sensing eLIDAR with VEGFR2 ecto+TM domains fused with tdMCP with a short GS linker in between, with C-terminal ER export sequence | this work |
| XZ359 EF1a-VEGFR1sp-3xFlag-VEGFR1(ecto+TM)-SL-ADAR2dd(QA)-export | https://benchling.com/s/seq-SCjCGOSyHVoo0DN07rHD?m=slm-5NW44qbJOTTcENDp1D2x | VEGF-sensing eLIDAR with VEGFR1 ecto+TM domains fused with ADAR2dd with a short GS linker in between, with C-terminal ER export sequence | this work |
| XZ360 EF1a-VEGFR2sp-HA-VEGFR2(Ecto+TM)-SL-TS-tdMCP-export | https://benchling.com/s/seq-Um5Uva465h7SREmJu2pK?m=slm-abh3uqod4OEyrEfos3VA | VEGF-sensing eLIDAR with VEGFR2 ecto+TM domains fused with tdMCP with a short GS linker in between, with Golgi traffic sequence and ER export sequence | this work |
| XZ361 EF1a-VEGFR1sp-3xFlag-VEGFR1(ecto+TM)-SL-TS-ADAR2dd(QA)-export | https://benchling.com/s/seq-jCPm0QCgaS9NIwzCkdDe?m=slm-XFvJMGdXmpTu4Cf5e9FH | VEGF-sensing eLIDAR with VEGFR1 ecto+TM domains fused with ADAR2dd with a short GS linker in between, with Golgi traffic sequence and ER export sequence | this work |
| CMV-TagBFP | https://benchling.com/s/seq-CXppV6e2AJdC4f0aHz1Q?m=slm-vxL31RcpvyBvmNH5qILq | CMV driven TagBFP | from lab members |
| APL0030 SFFV-bArrestin2-MCP (FLP-IN) | https://benchling.com/s/seq-qNqT6gxJe3hTlVrljmzy?m=slm-MWyX2aQZWpNDRJX5K432 | SFFV driven ARRB2-tdMCP with a short linker | this work |

| | | | |
|---|---|---|---|
| APL0031 SFFV-AVPR2-ADAR2dd (FLP-IN) | https://benchling.com/s/seq-KqmISxp6oMMTWoUWgQ6Q?m=slm-E9tPPTsXTVHteVyLxdcs | SFFV driven AVPR2-ADAR2dd (as in XZ030) | this work |
| APL0043 SFFV-CCR6-ADAR2dd (FLP-IN) | https://benchling.com/s/seq-V4lvq3SSPHfKZ8A2tNOt?m=slm-ypRWsbxyWVCA6k7FtKNW | SFFV dirven CCR6-ADAR2dd (as in XZ032) | this work |
| LSM075 SFFV-mCherry-SGc(stem-1(3xMS2)x2)cSG-EGFP (FLP-IN) | https://benchling.com/s/seq-VJC2oTYUjIzTPuPb2ZZP?m=slm-DnLp3tYv97DiWpQ4D0PC | SFFV driven LIDAR reporter with 2 tandem substrate stem loops | this work |
| LSM081 SFFV-mCherry-SGc(stem-1(3xMS2)2-stops)cSG-EGFP (FLP-IN) | https://benchling.com/s/seq-MraNnRegUvO2oej1FxQz?m=slm-59DFVbwfykA79jcSaGJI | SFFV driven LIDAR reporter with an additional stop codon on the stem loop | this work |
| LSM0250 (PB)EF1a-CCR6V2-ADAR2dd-IRESv0-PuroR-P2A-BFP | https://benchling.com/s/seq-lmgpbTI7rNegKGi7zlH1?m=slm-M8drgZiu3hfmCp27oFZ7 | PiggyBac donor plasmid to overexpress CCR6-ADAR2dd (gLIDAR) | this work |
| LSM0251 (PB)EF1a-FKBP-tm-ADAR2ddIRESv0-PuroR-P2A-BFP | https://benchling.com/s/seq-smWKsyIt03Z2FFOjYHDV?m=slm-WFfwcfOdqJsdnpzpg8Tg | PiggyBac donor plasmid to overexpress FKBP-TM-ADAR2dd (eLIDAR) | this work |
| LSM0252 (PB)EF1a-FKBP-ADAR2dd-IRESv0-PuroR-P2A-BFP | https://benchling.com/s/seq-FY3taRQKJfiRRJtMieOE?m=slm-hc9BJMi8IKG5zKBZyj1k | PiggyBac donor plasmid to overexpress FKBP-ADAR2dd (cLIDAR) | this work |
| LSM0253 (PB)EF1a-ADAR2dd-MCP-T2A-PuroR-P2A-BFP | https://benchling.com/s/seq-Pe5vbEoWl9SDuuP1hU29?m=slm-xn02TBg7O0GNb43ZaDH7 | PiggyBac donor plasmid to overexpress ADAR2dd-MCP | this work |
| LSM0254 (PB)EF1a-ADAR2-T2A-PuroR-P2A-BFP | https://benchling.com/s/seq-sZ5eaRTI1K0qHBABmKxn?m=slm-i6ARnmfbppi4JWt25YyR | PiggyBac donor plasmid to overexpress a full length ADAR2 | this work |
| LSM0258 (PB)EF1a-ADAR2dd-PCP-T2A-PuroR-P2A-BFP | https://benchling.com/s/seq-JIVU6TJpmnbA9Xnoy8vF?m=slm-hTDKA25ZhF0YiCyadB73 | PiggyBac donor plasmid to overexpress ADAR2dd-PCP | this work |
| JKCC121 CMV-puro-Ef1a-DIO-mCherry | https://benchling.com/s/seq-2PlljNPUyWayz8AV3o9c?m=slm-ENHeSTVRgwDPB2Rg5WJ7 | Lentivial plasmid for generating DIO-mCherry cell line | from lab members |

**Supplementary Table 2.** Overview of transfections.

| Figure | Cell line | Condition | Plasmid | Amount (ng) | Added ligand | Format |
|---|---|---|---|---|---|---|
| 1B | HEK293 | Cytosolic Binder-based LIDAR | EK0329 CMV-TO-FRB-MCP (FLP-IN) | 10 | AP21967 (100 nM) | 24-well |
| | | | XZ135 CMV-TO-FKBP-ADARdd(QA) (FLP-IN) | 10 | | |
| | | | EK0450 SFFV-mCherry-SGc(stem-1(3xMS2))cSG-EGFP | 200 | | |
| 1C | HEK293 | Extracellular Binder-based LIDAR | YH038 PGK-TO-FRB-tm-MCP | 50 | AP21967 (100 nM) | 24-well |
| | | | YH039 PGK-TO-FKBP-tm-GS-XTEN-ADAR2(DD-E488Q-T501A) | 50 | | |
| | | | EK0450 SFFV-mCherry-SGc(stem-1(3xMS2))cSG-EGFP | 200 | | |
| 1D | HEK293 | GPCR-based LIDAR | XZ024 CMV-TO-bArrestin2-MCP (FLP-IN) | 10 | AVP (50nM) | 24-well |
| | | | XZ030 CMV-TO-AVPR2-ADAR2(DD-E488Q-T501A) (FLP-IN) | 10 | | |
| | | | EK0450 SFFV-mCherry-SGc(stem-1(3xMS2))cSG-EGFP | 200 | | |
| 1E | HEK293 | RTK-based LIDAR | XZ038 CMV-TO-IGF1R-ADAR2(DD-E488Q-T501A) (FLP-IN) | 10 | IGF1 (200ng/mL) | 24-well |
| | | | XZ039 CMV-TO-Shc1-MCP (FLP-IN) | 10 | | |
| | | | EK0450 SFFV-mCherry-SGc(stem-1(3xMS2))cSG-EGFP | 200 | | |
| 2C | HEK293 | VEGF sensing eLIDAR with autocrine VEGF | EK0450 SFFV-mCherry-SGc(stem-1(3xMS2))cSG-EGFP | 200 | NA | 24-well |
| | | | XZ362 EF1a-VEGFR2sp-HA-VEGFR2(Ecto+TM)-TS-tdMCP-export | 50 | | |
| | | | XZ363 EF1a-VEGFR1sp-3xFlag-VEGFR1(ecto+TM)-TS-ADAR2dd(QA)-export | 50 | | |
| | | | YH001 CMV-TO-tm-GS+XTEN-ADAR2(DD-E396A)-RXR (FLP-IN) | 50 | | |
| | | | EK0479 CMV-TO-VEGF-A(165) (FLP-IN) | 50 | | |
| | | VEGF sensing eLIDAR with recombinant VEGF | EK0450 SFFV-mCherry-SGc(stem-1(3xMS2))cSG-EGFP | 200 | VEGF-165 (200ng/mL) | |
| | | | XZ362 EF1a-VEGFR2sp-HA-VEGFR2(Ecto+TM)-TS-tdMCP-export | 50 | | |
| | | | XZ363 EF1a-VEGFR1sp-3xFlag-VEGFR1(ecto+TM)-TS-ADAR2dd(QA)-export | 50 | | |
| | | | YH001 CMV-TO-tm-GS+XTEN-ADAR2(DD-E396A)-RXR (FLP-IN) | 50 | | |
| 2D, S2F-G | HEK293 | Stable integration of VEGF-sensing eLIDAR | XZ054 (PB)SFFV-LIDARsensor (hygR) | 200 | VEGF-165 (200ng/mL) | 24-well |
| | | | XZ377 (PB) EF1a-VEGFR1sp-3xFlag-VEGFR1(ecto+TM)-TS-ADAR2dd(QA)-export-IRESv0-VEGFR2sp-HA-VEGFR2(Ecto+TM)-TS-tdMCP-export (SV40-PuroR-BFP) | 200 | | |

| | | | | | | |
|---|---|---|---|---|---|---|
| | | | Piggybac Transposase from Gao et al. (2018) | 100 | | |
| 2F | HEK293 | Transient autocrine IL-2 sensing eLIDAR | EK0450 SFFV-mCherry-SGc(stem-1(3xMS2))cSG-EGFP | 200 | NA | 24-well |
| | | | XZ375 EF1a-IL2Rbsp-3xFLAG-IL2Rb(ECD+TM)-TS-ADAR2dd-export | 50 | | |
| | | | XZ374 EF1a-IL2Rgsp-3xHA-IL2Rg(ECD+TM)-TS-tdMCP-export | 50 | | |
| | | | YH001 CMV-TO-tm-GS+XTEN-ADAR2(DD-E396A)-RXR (FLP-IN) | 50 | | |
| | | | CMV-hIL2 | 50 | | |
| | | Transient recombinant IL-2 sensing eLIDAR | EK0450 SFFV-mCherry-SGc(stem-1(3xMS2))cSG-EGFP | 200 | Recombinant human IL-2 (1 µg/mL) | 24-well |
| | | | XZ375 EF1a-IL2Rbsp-3xFLAG-IL2Rb(ECD+TM)-TS-ADAR2dd-export | 50 | | |
| | | | XZ374 EF1a-IL2Rgsp-3xHA-IL2Rg(ECD+TM)-TS-tdMCP-export | 50 | | |
| | | | YH001 CMV-TO-tm-GS+XTEN-ADAR2(DD-E396A)-RXR (FLP-IN) | 50 | | |
| 2G, S2H-I | HEK293 | Stable integration of Il-2-sensing eLIDAR | XZ054 (PB)SFFV-LIDARsensor (hygR) | 200 | Recombinant human IL-2 (1 µg/mL) | 24-well |
| | | | XZ393 (PB) EF1a-IL2Rbsp-FLAG-IL2Rb(ECD+TM)-TS-ADAR2dd-export-IRESv0-IL2Rgsp-3xHA-IL2Rg(ECD+TM)-TS-tdMCP-export (SV40-PuroR-BFP) | 200 | | |
| | | | Piggybac Transposase from Gao et al. (2018) | 100 | | |
| 2I | HEK293 | CCR6 | XZ024 CMV-TO-bArrestin2-MCP (FLP-IN) | 10 | CCL20 (100ng/mL) | 24-well |
| | | | XZ031 CMV-TO-CCR6-ADAR2dd(E488Q, T501A) | 10 | | |
| | | | EK0450 SFFV-mCherry-SGc(stem-1(3xMS2))cSG-EGFP | 200 | | |
| | | CCR7 | XZ024 CMV-TO-bArrestin2-MCP (FLP-IN) | 10 | CCL19 (120nM) | 24-well |
| | | | LSM044-3 CMV-TO-CCR7-ADAR2dd(E488Q, T501A) (FLP-IN) | 10 | | |
| | | | EK0450 SFFV-mCherry-SGc(stem-1(3xMS2))cSG-EGFP | 200 | | |
| | | CX3CR1 | XZ024 CMV-TO-bArrestin2-MCP (FLP-IN) | 10 | CX3CL1 (12nM) | 24-well |
| | | | LSM068 CMV-TO-CX3CR1-ADAR2dd(E488Q, T501A) (FLP-IN) | 10 | | |
| | | | EK0450 SFFV-mCherry-SGc(stem-1(3xMS2))cSG-EGFP | 200 | | |
| | | hOXTR | XZ024 CMV-TO-bArrestin2-MCP (FLP-IN) | 10 | Oxytocin (2uM) | 24-well |
| | | | XZ113 CMV-TO-signal-hOXTR-ADAR2dd(DD-E488Q-T501A) (FLP-IN) | 10 | | |
| | | | EK0450 SFFV-mCherry-SGc(stem-1(3xMS2))cSG-EGFP | 200 | | |
| | | B2AR | XZ024 CMV-TO-bArrestin2-MCP (FLP-IN) | 10 | | 24-well |

| | | | | | | |
|---|---|---|---|---|---|---|
| | | | XZ142 CMV-TO-signal-B2AR-ADAR2dd(DD-E488Q-T501A) (FLP-IN) | 10 | Isoproterenol (100uM) | |
| | | | EK0450 SFFV-mCherry-SGc(stem-1(3xMS2))cSG-EGFP | 200 | | |
| 2K | HEK293 | oLIDAR with AVP and CX3CL1 sensing | LSM060 CMV-TO-AVPR2-(10aa-linker)-MCP (FLP-IN) | 5 | CX3CL1 (12nM); AVP (50nM) | 24-well |
| | | | LSM102 CMV-TO-CX3CR1-10aa-PCP (FLP-IN) | 5 | | |
| | | | LSM059 CMV-TO-bArrestin2-(31-linker)-ADAR2dd(E488Q, T501A) (FLP-IN) | 10 | | |
| | | | LSM053 SFFV-mCherry-SGc(stem-1(3xPP7))cSG-mtagBFP (FLP-IN) | 150 | | |
| | | | EK0450 SFFV-mCherry-SGc(stem-1(3xMS2))cSG-EGFP | 150 | | |
| 2L | Colo320-DM | Rapalog sensing eLIDAR | YH038 PGK-TO-FRB-tm-MCP | 50 | AP21967 (100 nM) | 24-well |
| | | | YH039 PGK-TO-FKBP-tm-GS-XTEN-ADAR2(DD-E488Q-T501A) | 50 | | |
| | | | EK0450 SFFV-mCherry-SGc(stem-1(3xMS2))cSG-EGFP | 200 | | |
| | | CCL20 sensing gLIDAR | XZ024 CMV-TO-bArrestin2-MCP (FLP-IN) | 10 | CCL20 (100ng/mL) | 24-well |
| | | | XZ032 CMV-TO-CCR6-V2-ADAR2dd(E488Q, T501A) | 10 | | |
| | | | EK0450 SFFV-mCherry-SGc(stem-1(3xMS2))cSG-EGFP | 200 | | |
| 2M | SNU-475 | Rapalog sensing eLIDAR | YH038 PGK-TO-FRB-tm-MCP | 50 | AP21967 (100 nM) | 24-well |
| | | | YH039 PGK-TO-FKBP-tm-GS-XTEN-ADAR2(DD-E488Q-T501A) | 50 | | |
| | | | EK0450 SFFV-mCherry-SGc(stem-1(3xMS2))cSG-EGFP | 200 | | |
| | | CCL20 sensing gLIDAR | XZ024 CMV-TO-bArrestin2-MCP (FLP-IN) | 10 | CCL20 (100ng/mL) | 24-well |
| | | | XZ032 CMV-TO-CCR6-V2-ADAR2dd(E488Q, T501A) | 10 | | |
| | | | EK0450 SFFV-mCherry-SGc(stem-1(3xMS2))cSG-EGFP | 200 | | |
| 2N | C3H | Rapalog sensing eLIDAR | YH038 PGK-TO-FRB-tm-MCP | 50 | AP21967 (100 nM) | 24-well |
| | | | YH039 PGK-TO-FKBP-tm-GS-XTEN-ADAR2(DD-E488Q-T501A) | 50 | | |
| | | | EK0450 SFFV-mCherry-SGc(stem-1(3xMS2))cSG-EGFP | 200 | | |
| 2O | S2 | Rapalog sensing cLIDAR | APL0038 Ac5-mCherry-SGc(stem-1(3xMS2))cSG-EGFP | 150 | AP21967 (100 nM) | 24-well |
| | | | APL0040 Ac5-FKBP-ADARdd | 300 | | |
| | | | APL0041 Ac5-FRB-MCP | 300 | | |
| 3B, S7A-C | HEK293 | Stable integration of CCR6-based gLIDAR | Piggybac Transposase from Gao et al. (2018) | 100 | CCL19 (120nM), AP1903 (4nM) | 24-well |
| | | | XZ079 (PB)EF1a-CCR6-ADARdd-P2A-PuroR-IRES-ARRB2-MCP | 200 | | |
| | | | LSM043_2 (PB)SFFV-LIDARSensor(mCherry->Casp9-2A-GFP)-hygro | 200 | | |

| | | | | | | |
|---|---|---|---|---|---|---|
| 3D | HEK293 | Cytosolic Binder-based LIDAR | XZ400 SFFV-mCherry-SGc(stem-1(3xMS2))cSG-TEVP | 200 | AP21967 (100 nM) | 24-well |
| | | | PB-CMV-Cit-hivs-DHFR | 50 | | |
| | | | XZ135 CMV-TO-FKBP-ADARdd(QA) (FLP-IN) | 10 | | |
| | | | EK0329 CMV-TO-FRB-MCP (FLP-IN) | 10 | | |
| 3E, S8A | HEK293 | CCR6 gLIDAR controlling citrine stability with TEV output | EK0461 SFFV-mCherry-SGc(stem-1(3xMS2))cSG-TEVp (FLP-IN) | 80 | CCL20 (100ng/mL) | 96-well |
| | | | PB-CMV-Cit-hivs-DHFR | 20 | | |
| | | | XZ024 CMV-TO-bArrestin2-MCP (FLP-IN) | 4 | | |
| | | | XZ032 CMV-TO-CCR6-V2-ADAR2dd(E488Q, T501A) | 4 | | |
| 3G | HEK293 | Rapalog-sensing cLIDAR controlling SEAP secretion | XZ392 SFFV-secNLuc-SGc(stem-1(3xMS2))cSG(tight)-ssSEAP | 200 | AP21967 (100 nM) | 24-well |
| | | | AV282_CMVTO-SEAP-26SFur-B2AD-tevs-RXR6.2 | 10 | | |
| | | | CMVTO-secNluc | 10 | | |
| | | | XZ135 CMV-TO-FKBP-ADARdd(QA) (FLP-IN) | 10 | | |
| | | | EK0329 CMV-TO-FRB-MCP (FLP-IN) | 10 | | |
| 3I, S8B, S8C | HEK293 | CCL20-sensing gLIDAR controlling IgG membrane display via RELEASE | AV324_CMVTO-hIgG1-B2AD-tevs-KKMP | 4 | CCL20 (100ng/mL) | 96-well |
| | | | EK0461 SFFV-mCherry-SGc(stem-1(3xMS2))cSG-TEVp (FLP-IN) | 80 | | |
| | | | XZ032 CMV-TO-CCR6-V2-ADAR2dd(E488Q, T501A) | 4 | | |
| | | | XZ024 CMV-TO-bArrestin2-MCP (FLP-IN) | 4 | | |
| 4A | HEK293 | Sender cell - secreted | XZ034 CMV-TO-CCL20 | 450 | Conditioned media from sender cells producing secreted CCL20 | 24-well |
| | | Receiver cells | XZ024 CMV-TO-bArrestin2-MCP (FLP-IN) | 10 | | |
| | | | XZ032 CMV-TO-CCR6-V2-ADAR2dd(E488Q, T501A) | 10 | | |
| | | | EK0450 SFFV-mCherry-SGc(stem-1(3xMS2))cSG-EGFP | 200 | | |
| 4B | HEK293 | Sender cell - membrane | XZ033 CMV-TO-CCL20-pre-mGRASP | 450 | Co-culture with sender cells expressing membrane CCL20 | 24-well |
| | | Receiver cells | XZ024 CMV-TO-bArrestin2-MCP (FLP-IN) | 10 | | |
| | | | XZ032 CMV-TO-CCR6-V2-ADAR2dd(E488Q, T501A) | 10 | | |
| | | | EK0450 SFFV-mCherry-SGc(stem-1(3xMS2))cSG-EGFP | 200 | | |
| 4C-F, S9 | HEK293 | Stable integration of receiver cell: | Same as 3B | Same as 3B | NA | 24-well |

| | | | | | | |
|---|---|---|---|---|---|---|
| | | CCR6-based gLIDAR (cLSM01) | | | | |
| | HEK293 | Stable integration of sender cell: secreted CCL20 | Piggybac Transposase from Gao et al. (2018) | 50 | NA | 24-well |
| | | | XZ091 (PB) CMV-sCCL20 (PuroR-BFP) | 450 | | |
| | HEK293 | Stable integration of sender cell: membrane-bound CCL20 | Piggybac Transposase from Gao et al. (2018) | 50 | NA | 24-well |
| | | | XZ076 (PB) CMV-mCCL20 (PuroR-BFP) | 50 | | |
| 5B-C | HEK293 | Single transcript | XZ093 SFFV-sensor-IRESv0-LIDARst(IRESv0-AVPR2) | 400 | AVP (100nM) | 24-well |
| 5E | HEK293 | mRNA delivered AVP sensing gLIDAR | IVT mRNA from XZ387 (IVT) CMV-TO-T7-AVPR2-V2-ADAR2dd-hybridUTR3 (100% m1Ψ) | 400 | AVP (200nM) | 24-well |
| | | | IVT mRNA from XZ089 (IVT) CMV-TO-T7-ARRB2-MCP-hybridUTR3 (100% m1Ψ) | 400 | | |
| | | | IVT mRNA from XZ073 (IVT) CMV-TO-T7-LIDARsensor-hybridUTR3 (0% m1Ψ, 0% m5C) | 200 | | |
| 5F, S5A-C | HEK293 | mRNA delivered CCL20 sensing gLIDAR | IVT mRNA from XZ088 (IVT) CMV-TO-T7-CCR6-V2-ADAR2dd-hybridUTR3 (100% m1Ψ) | 400 | CCL20 (100ng/mL) | 24-well |
| | | | IVT mRNA from XZ089 (IVT) CMV-TO-T7-ARRB2-MCP-hybridUTR3 (100% m1Ψ) | 400 | | |
| | | | IVT mRNA from XZ073 (IVT) CMV-TO-T7-LIDARsensor-hybridUTR3 (0% m1Ψ, 0% m5C) | 200 | | |
| 5H, S5D | cJKCC1 21 (derived from HEK293 ) | AVP sensing gLIDAR with Cre output | IVT mRNA from XZ387 (IVT) CMV-TO-T7-AVPR2-V2-ADAR2dd-hybridUTR3 (100% m1Ψ) | 400 | AVP (200nM) | 24-well |
| | | | IVT mRNA from XZ089 (IVT) CMV-TO-T7-ARRB2-MCP-hybridUTR3 (100% m1Ψ) | 400 | | |
| | | | IVT mRNA from XZ388 (IVT) CMV-TO-T7-EGFP-SGc(stem-1(3xMS2))cSG-Cre-hybridUTR3 (0% m1Ψ, 0% m5C) | 200 | | |
| 5J | cNK90( derived from HEK293 ) | AVP sensing mRNA delivered gLIDAR with rtetR-NZF output | IVT mRNA from XZ387 (IVT) CMV-TO-T7-AVPR2-V2-ADAR2dd-hybridUTR3 (100% m1Ψ) | 400 | AVP (200nM) | 24-well |
| | | | IVT mRNA from XZ089 (IVT) CMV-TO-T7-ARRB2-MCP-hybridUTR3 (100% m1Ψ) | 400 | | |
| | | | IVT mRNA from XZ390 (IVT) CMV-TO-T7-mCherry-SGc(stem-1(3xMS2))cSG-rTetR-NZF-NLS-hybridUTR3 (0% m1Ψ, 0% m5C) | 200 | | |
| 5K-L | HEK293 | Reporter without base modifications | IVT mRNA from XZ088 (IVT) CMV-TO-T7-CCR6-V2-ADAR2dd-hybridUTR3 (100% m1Ψ) | 400 | CCL20 (100ng/mL) | 24-well |

| | | | | | | |
|---|---|---|---|---|---|---|
| | | | IVT mRNA from XZ089 (IVT) CMV-TO-T7-ARRB2-MCP-hybridUTR3 (100% m1Ψ) | 400 | | |
| | | | IVT mRNA from XZ073 (IVT) CMV-TO-T7-LIDARsensor-hybridUTR3 (0% m1Ψ, 0% m5C) | 200 | | |
| | | Reporter with 100% m5C | IVT mRNA from XZ088 (IVT) CMV-TO-T7-CCR6-V2-ADAR2dd-hybridUTR3 (100% m1Ψ) | 400 | | |
| | | | IVT mRNA from XZ089 (IVT) CMV-TO-T7-ARRB2-MCP-hybridUTR3 (100% m1Ψ) | 400 | | |
| | | | IVT mRNA from XZ073 (IVT) CMV-TO-T7-LIDARsensor-hybridUTR3 (0% m1Ψ, 100% m5C) | 200 | | |
| | | Reporter with 100% m1Ψ | IVT mRNA from XZ088 (IVT) CMV-TO-T7-CCR6-V2-ADAR2dd-hybridUTR3 (100% m1Ψ) | 400 | | |
| | | | IVT mRNA from XZ089 (IVT) CMV-TO-T7-ARRB2-MCP-hybridUTR3 (100% m1Ψ) | 400 | | |
| | | | IVT mRNA from XZ073 (IVT) CMV-TO-T7-LIDARsensor-hybridUTR3 (100% m1Ψ, 0% m5C) | 200 | | |
| S1B | HEK293 | 1x MS2 | EK0329 CMV-TO-FRB-MCP (FLP-IN) | 10 | AP21967 (100 nM) | 24-well |
| | | | XZ135 CMV-TO-FKBP-ADARdd(QA) (FLP-IN) | 10 | | |
| | | | EK0450 SFFV-mCherry-SGc(stem-1(1xMS2))cSG-EGFP (FLP-IN) | 200 | | |
| | | 2x MS2 | EK0329 CMV-TO-FRB-MCP (FLP-IN) | 10 | AP21967 (100 nM) | |
| | | | XZ135 CMV-TO-FKBP-ADARdd(QA) (FLP-IN) | 10 | | |
| | | | XZ136 SFFV-mCherry-SGc(stem-1(2xMS2))cSG-EGFP (FLP-IN) | 200 | | |
| | | 3x MS2 | EK0329 CMV-TO-FRB-MCP (FLP-IN) | 10 | AP21967 (100 nM) | |
| | | | XZ135 CMV-TO-FKBP-ADARdd(QA) (FLP-IN) | 10 | | |
| | | | EK0450 SFFV-mCherry-SGc(stem-1(3xMS2))cSG-EGFP | 200 | | |
| S1C | HEK293 | 1x MS2 | YH038 PGK-TO-FRB-tm-MCP | 50 | AP21967 (100 nM) | 24-well |
| | | | YH039 PGK-TO-FKBP-tm-GS-XTEN-ADAR2(DD-E488Q-T501A) | 50 | | |
| | | | EK0450 SFFV-mCherry-SGc(stem-1(1xMS2))cSG-EGFP (FLP-IN) | 200 | | |
| | | 2x MS2 | YH038 PGK-TO-FRB-tm-MCP | 50 | AP21967 (100 nM) | |
| | | | YH039 PGK-TO-FKBP-tm-GS-XTEN-ADAR2(DD-E488Q-T501A) | 50 | | |
| | | | XZ136 SFFV-mCherry-SGc(stem-1(2xMS2))cSG-EGFP (FLP-IN) | 200 | | |

| | | | | | | |
|---|---|---|---|---|---|---|
| | | 3x MS2 | YH038 PGK-TO-FRB-tm-MCP | 50 | AP21967 (100 nM) | |
| | | 3x MS2 | YH039 PGK-TO-FKBP-tm-GS-XTEN-ADAR2(DD-E488Q-T501A) | 50 | AP21967 (100 nM) | |
| | | 3x MS2 | EK0450 SFFV-mCherry-SGc(stem-1(3xMS2))cSG-EGFP | 200 | AP21967 (100 nM) | |
| S1D | HEK293 | E488Q - no FRB | EK0330 CMV-TO-FKBP-ADAR2(DD-E488Q) (FLP-IN) | 10 | NA | 24-well |
| S1D | HEK293 | E488Q - no FRB | EK0450 SFFV-mCherry-SGc(stem-1(3xMS2))cSG-EGFP | 200 | NA | 24-well |
| S1D | HEK293 | E488Q | EK0329 CMV-TO-FRB-MCP (FLP-IN) | 10 | AP21967 (100 nM) | 24-well |
| S1D | HEK293 | E488Q | EK0330 CMV-TO-FKBP-ADAR2(DD-E488Q) (FLP-IN) | 10 | AP21967 (100 nM) | 24-well |
| S1D | HEK293 | E488Q | EK0450 SFFV-mCherry-SGc(stem-1(3xMS2))cSG-EGFP | 200 | AP21967 (100 nM) | 24-well |
| S1D | HEK293 | E488Q T501A - no FRB | XZ135 CMV-TO-FKBP-ADARdd(QA) (FLP-IN) | 10 | NA | 24-well |
| S1D | HEK293 | E488Q T501A - no FRB | EK0450 SFFV-mCherry-SGc(stem-1(3xMS2))cSG-EGFP | 200 | NA | 24-well |
| S1D | HEK293 | E488Q T501A | EK0329 CMV-TO-FRB-MCP (FLP-IN) | 10 | AP21967 (100 nM) | 24-well |
| S1D | HEK293 | E488Q T501A | XZ135 CMV-TO-FKBP-ADARdd(QA) (FLP-IN) | 10 | AP21967 (100 nM) | 24-well |
| S1D | HEK293 | E488Q T501A | EK0450 SFFV-mCherry-SGc(stem-1(3xMS2))cSG-EGFP | 200 | AP21967 (100 nM) | 24-well |
| S1E | HEK293 | E488Q - no ARRB2 | XZ029 CMV-TO-AVPR2-ADAR2(DD-E488Q) (FLP-IN) | 10 | NA | 24-well |
| S1E | HEK293 | E488Q - no ARRB2 | EK0450 SFFV-mCherry-SGc(stem-1(3xMS2))cSG-EGFP | 200 | NA | 24-well |
| S1E | HEK293 | E488Q | XZ024 CMV-TO-bArrestin2-MCP (FLP-IN) | 10 | AVP (50nM) | 24-well |
| S1E | HEK293 | E488Q | XZ029 CMV-TO-AVPR2-ADAR2(DD-E488Q) (FLP-IN) | 10 | AVP (50nM) | 24-well |
| S1E | HEK293 | E488Q | EK0450 SFFV-mCherry-SGc(stem-1(3xMS2))cSG-EGFP | 200 | AVP (50nM) | 24-well |
| S1E | HEK293 | E488Q T501A - no ARRB2 | XZ030 CMV-TO-AVPR2-ADAR2(DD-E488Q-T501A) (FLP-IN) | 10 | NA | 24-well |
| S1E | HEK293 | E488Q T501A - no ARRB2 | EK0450 SFFV-mCherry-SGc(stem-1(3xMS2))cSG-EGFP | 200 | NA | 24-well |
| S1E | HEK293 | E488Q T501A | XZ024 CMV-TO-bArrestin2-MCP (FLP-IN) | 10 | AVP (50nM) | 24-well |
| S1E | HEK293 | E488Q T501A | XZ030 CMV-TO-AVPR2-ADAR2(DD-E488Q-T501A) (FLP-IN) | 10 | AVP (50nM) | 24-well |
| S1E | HEK293 | E488Q T501A | EK0450 SFFV-mCherry-SGc(stem-1(3xMS2))cSG-EGFP | 200 | AVP (50nM) | 24-well |
| S1F | HEK293 | 47 aa linker length | XZ024 CMV-TO-bArrestin2-MCP (FLP-IN) | 10 | AVP (50nM) | 24-well |
| S1F | HEK293 | 47 aa linker length | XZ158 CMV-TO-AVPR2-45ECL-ADAR2dd(QA) (FLP-IN) | 10 | AVP (50nM) | 24-well |
| S1F | HEK293 | 47 aa linker length | EK0450 SFFV-mCherry-SGc(stem-1(3xMS2))cSG-EGFP | 200 | AVP (50nM) | 24-well |
| S1F | HEK293 | 31 aa linker length | XZ024 CMV-TO-bArrestin2-MCP (FLP-IN) | 10 | AVP (50nM) | 24-well |
| S1F | HEK293 | 31 aa linker length | XZ156 CMV-TO-AVPR2-GSXTEN-ADAR2dd(QA) (FLP-IN) | 10 | AVP (50nM) | 24-well |
| S1F | HEK293 | 31 aa linker length | EK0450 SFFV-mCherry-SGc(stem-1(3xMS2))cSG-EGFP | 200 | AVP (50nM) | 24-well |
| S1F | HEK293 | 10 aa linker length | XZ024 CMV-TO-bArrestin2-MCP (FLP-IN) | 10 | AVP (50nM) | 24-well |

| | | | | | | |
|---|---|---|---|---|---|---|
| | | | XZ157 CMV-TO-AVPR2-10L-ADAR2dd(QA) (FLP-IN) | 10 | | |
| | | | EK0450 SFFV-mCherry-SGc(stem-1(3xMS2))cSG-EGFP | 200 | | |
| | | 4 aa linker length | XZ024 CMV-TO-bArrestin2-MCP (FLP-IN) | 10 | AVP (50nM) | |
| | | | XZ030 CMV-TO-AVPR2-ADAR2(DD-E488Q-T501A) (FLP-IN) | 10 | | |
| | | | EK0450 SFFV-mCherry-SGc(stem-1(3xMS2))cSG-EGFP | 200 | | |
| S1G | HEK293 | AVPR2-based gLIDAR | XZ024 CMV-TO-bArrestin2-MCP (FLP-IN) | 10 | AVP (50nM) | 24-well |
| | | | XZ030 CMV-TO-AVPR2-ADAR2(DD-E488Q-T501A) (FLP-IN) | 10 | | |
| | | | EK0450 SFFV-mCherry-SGc(stem-1(3xMS2))cSG-EGFP | 200 | | |
| S1H-I | HEK293 | Stable integration of AVPR-based gLIDAR | Piggybac Transposase from Gao et al. (2018) | 100 | AVP (50nM) | 24-well |
| | | | LSM042 (PB)EF1a-AVPR-ADARdd-P2A-PuroR-IRES-ARRB2-MCP | 200 | | |
| | | | LSM061 (PB)SFFV-LIDARsensor(mcherry-BFP) (hygR) | 200 | | |
| S1J-L | HEK293 | AVPR, TF-based system inspired by TANGO | NK90-pTRE3G-EGFP-stop (PB) | 10 | AVP (50nM) | 24-well |
| | | | LSM108 CMV-TO-AVPR-TEVcs-(noLinker)-tTA2 (FLP-IN) | 10 | | |
| | | | pCDNA3.1(+)-CMV-bArrestin2-TEV | 10 | | |
| | | | CMV-TO-mCherry from Gao et al. (2018) | 20 | | |
| | | AVPR-based LIDAR | XZ024 CMV-TO-bArrestin2-MCP (FLP-IN) | 10 | AVP (50nM) | |
| | | | XZ030 CMV-TO-AVPR2-ADAR2(DD-E488Q-T501A) (FLP-IN) | 10 | | |
| | | | EK0450 SFFV-mCherry-SGc(stem-1(3xMS2))cSG-EGFP | 200 | | |
| S2B | HEK293 | VEGF eLIDAR with GS linker, no ER export sequence | EK0450 SFFV-mCherry-SGc(stem-1(3xMS2))cSG-EGFP | 200 | VEGF-165 (1 µg/mL) | 24-well |
| | | | XZ344 EF1a-VEGFR2sp-HA-VEGFR2(Ecto+TM)-SL-tdMCP | 50 | | |
| | | | XZ345 EF1a-VEGFR1sp-3xFlag-VEGFR1(ecto+TM)-SL-ADAR2dd(QA) | 50 | | |
| | | VEGF eLIDAR with GS linker and ER export sequence | EK0450 SFFV-mCherry-SGc(stem-1(3xMS2))cSG-EGFP | 200 | VEGF-165 (1 µg/mL) | |
| | | | XZ358 EF1a-VEGFR2sp-HA-VEGFR2(Ecto+TM)-SL-tdMCP-export | 50 | | |
| | | | XZ359 EF1a-VEGFR1sp-3xFlag-VEGFR1(ecto+TM)-SL-ADAR2dd(QA)-export | 50 | | |
| | | VEGF eLIDAR with GS linker + Golgi traffic | EK0450 SFFV-mCherry-SGc(stem-1(3xMS2))cSG-EGFP | 200 | VEGF-165 (1 µg/mL) | |
| | | | XZ360 EF1a-VEGFR2sp-HA-VEGFR2(Ecto+TM)-SL-TS-tdMCP-export | 50 | | |

| | | | | | | |
|---|---|---|---|---|---|---|
| | | sequence and ER export sequence | XZ361 EF1a-VEGFR1sp-3xFlag-VEGFR1(ecto+TM)-SL-TS-ADAR2dd(QA)-export | 50 | | |
| | | VEGF eLIDAR with Golgi traffic sequence (no GS linker) and ER export sequence | EK0450 SFFV-mCherry-SGc(stem-1(3xMS2))cSG-EGFP | 200 | VEGF-165 (1 µg/mL) | |
| | | | XZ362 EF1a-VEGFR2sp-HA-VEGFR2(Ecto+TM)-TS-tdMCP-export | 50 | | |
| | | | XZ363 EF1a-VEGFR1sp-3xFlag-VEGFR1(ecto+TM)-TS-ADAR2dd(QA)-export | 50 | | |
| S2C | HEK293 | Negative staining control | CMV-TagBFP | 10 | NA | 24-well |
| S2D | HEK293 | VEGF eLIDAR with GS linker, no ER export sequence, test by staining | CMV-TagBFP | 10 | NA | 24-well |
| | | | XZ344 EF1a-VEGFR2sp-HA-VEGFR2(Ecto+TM)-SL-tdMCP | 100 | | |
| | | | XZ345 EF1a-VEGFR1sp-3xFlag-VEGFR1(ecto+TM)-SL-ADAR2dd(QA) | 100 | | |
| | | VEGF eLIDAR with GS linker and ER export sequence, test by staining | CMV-TagBFP | 10 | NA | 24-well |
| | | | XZ358 EF1a-VEGFR2sp-HA-VEGFR2(Ecto+TM)-SL-tdMCP-export | 100 | | |
| | | | XZ359 EF1a-VEGFR1sp-3xFlag-VEGFR1(ecto+TM)-SL-ADAR2dd(QA)-export | 100 | | |
| | | VEGF eLIDAR with GS linker + Golgi traffic sequence and ER export sequence, test by staining | CMV-TagBFP | 10 | NA | 24-well |
| | | | XZ360 EF1a-VEGFR2sp-HA-VEGFR2(Ecto+TM)-SL-TS-tdMCP-export | 100 | | |
| | | | XZ361 EF1a-VEGFR1sp-3xFlag-VEGFR1(ecto+TM)-SL-TS-ADAR2dd(QA)-export | 100 | | |
| | | VEGF eLIDAR with Golgi traffic sequence (no GS linker) and ER export sequence, test by staining | CMV-TagBFP | 10 | NA | 24-well |
| | | | XZ362 EF1a-VEGFR2sp-HA-VEGFR2(Ecto+TM)-TS-tdMCP-export | 100 | | |
| | | | XZ363 EF1a-VEGFR1sp-3xFlag-VEGFR1(ecto+TM)-TS-ADAR2dd(QA)-export | 100 | | |
| S2E | HEK293 | VEGF eLIDAR with Catalytically inactive ADARdd | EK0450 SFFV-mCherry-SGc(stem-1(3xMS2))cSG-EGFP | 200 | VEGF-165 (200ng/mL) | 24-well |
| | | | XZ362 EF1a-VEGFR2sp-HA-VEGFR2(Ecto+TM)-TS-tdMCP-export | 100 | | |
| | | | XZ363 EF1a-VEGFR1sp-3xFlag-VEGFR1(ecto+TM)-TS-ADAR2dd(QA)-export | 100 | | |
| | | | YH001 CMV-TO-tm-GS+XTEN-ADAR2(DD-E396A)-RXR (FLP-IN) | 0/10/ 50/100 | | |

| | | | | | | |
|---|---|---|---|---|---|---|
| S3A | HEK293 | AVPR2 no V2 tail | LSM104 CMV-TO-AVPR2-(noExtraV2)-ADAR2(DD-E488Q-T501A) (FLP-IN) | 10 | AVP (50nM) | 24-well |
| | | | XZ024 CMV-TO-bArrestin2-MCP (FLP-IN) | 10 | | |
| | | | EK0450 SFFV-mCherry-SGc(stem-1(3xMS2))cSG-EGFP | 200 | | |
| | | AVPR2 with V2 tail | XZ030 CMV-TO-AVPR2-ADAR2(DD-E488Q-T501A) (FLP-IN) | 10 | AVP (50nM) | |
| | | | XZ024 CMV-TO-bArrestin2-MCP (FLP-IN) | 10 | | |
| | | | EK0450 SFFV-mCherry-SGc(stem-1(3xMS2))cSG-EGFP | 200 | | |
| | | CX3CR1 no V2 tail | LSM105 CMV-TO-CX3CR1-(noV2)-ADAR2dd(E488Q, T501A) (FLP-IN) | 10 | CX3CL1 (12nM) | |
| | | | XZ024 CMV-TO-bArrestin2-MCP (FLP-IN) | 10 | | |
| | | | EK0450 SFFV-mCherry-SGc(stem-1(3xMS2))cSG-EGFP | 200 | | |
| | | CX3CR1 with V2 tail | LM068 CMV-TO-CX3CR1-ADAR2dd(E488Q, T501A) (FLP-IN) | 10 | CX3CL1 (12nM) | |
| | | | XZ024 CMV-TO-bArrestin2-MCP (FLP-IN) | 10 | | |
| | | | EK0450 SFFV-mCherry-SGc(stem-1(3xMS2))cSG-EGFP | 200 | | |
| | | CCR6 no V2 tail | XZ031 CMV-TO-CCR6-ADAR2dd(E488Q, T501A) | 10 | CCL20 (100ng/mL) | |
| | | | XZ024 CMV-TO-bArrestin2-MCP (FLP-IN) | 10 | | |
| | | | EK0450 SFFV-mCherry-SGc(stem-1(3xMS2))cSG-EGFP | 200 | | |
| | | CCR6 with V2 tail | XZ032 CMV-TO-CCR6-V2-ADAR2dd(E488Q, T501A) | 10 | CCL20 (100ng/mL) | |
| | | | XZ024 CMV-TO-bArrestin2-MCP (FLP-IN) | 10 | | |
| | | | EK0450 SFFV-mCherry-SGc(stem-1(3xMS2))cSG-EGFP | 200 | | |
| | | B2AR no V2 tail | XZ142 CMV-TO-signal-B2AR-ADAR2dd(DD-E488Q-T501A) (FLP-IN) | 10 | Isoproterenol (100uM) | |
| | | | XZ024 CMV-TO-bArrestin2-MCP (FLP-IN) | 10 | | |
| | | | EK0450 SFFV-mCherry-SGc(stem-1(3xMS2))cSG-EGFP | 200 | | |
| | | B2AR with V2 tail | XZ110 CMV-TO-signal-B2AR-V2-ADAR2dd(DD-E488Q-T501A) (FLP-IN) | 10 | Isoproterenol (100uM) | |
| | | | XZ024 CMV-TO-bArrestin2-MCP (FLP-IN) | 10 | | |
| | | | EK0450 SFFV-mCherry-SGc(stem-1(3xMS2))cSG-EGFP | 200 | | |
| | | OXTR no V2 tail | XZ113 CMV-TO-signal-hOXTR-ADAR2dd(DD-E488Q-T501A) (FLP-IN) | 10 | Oxytocin (2uM) | |
| | | | XZ024 CMV-TO-bArrestin2-MCP (FLP-IN) | 10 | | |
| | | | EK0450 SFFV-mCherry-SGc(stem-1(3xMS2))cSG-EGFP | 200 | | |
| | | OXTR with V2 tail | XZ105 CMV-TO-signal-hOXTR-V2-ADAR2dd(DD-E488Q-T501A) (FLP-IN) | 10 | Oxytocin (2uM) | |

| | | | | | | |
|---|---|---|---|---|---|---|
| | | | XZ024 CMV-TO-bArrestin2-MCP (FLP-IN) | 10 | | |
| | | | EK0450 SFFV-mCherry-SGc(stem-1(3xMS2))cSG-EGFP | 200 | | |
| S3B | HEK293 | Stable integration of CCL20-sensing gLIDAR | Piggybac Transposase from Gao et al. (2018) | 100 | CCL20 (0-500ng/mL) | 24-well |
| | | | XZ079 (PB)EF1a-CCR6-ADARdd-P2A-PuroR-IRES-ARRB2-MCP | 200 | | |
| | | | XZ054 (PB)SFFV-LIDARsensor (hygR) | 200 | | |
| S3C | HEK293 | CCR6 - transient transfection | XZ032 CMV-TO-CCR6-V2-ADAR2dd(E488Q, T501A) | 10 | CCL20 (0-500ng/mL) | 24-well |
| | | | XZ024 CMV-TO-bArrestin2-MCP (FLP-IN) | 10 | | |
| | | | EK0450 SFFV-mCherry-SGc(stem-1(3xMS2))cSG-EGFP | 200 | | |
| S3E | HEK293 | AVPR2-based oLIDAR | LSM060 CMV-TO-AVPR2-(10aa-linker)-MCP (FLP-IN) | 10 | AVP (50nM) | 24-well |
| | | | LSM059 CMV-TO-bArrestin2-(31-linker)-ADAR2dd(E488Q, T501A) (FLP-IN) | 10 | | |
| | | | EK0450 SFFV-mCherry-SGc(stem-1(3xMS2))cSG-EGFP | 200 | | |
| | | CCR6-based oLIDAR | LSM052 CMV-TO-CCR6-PCP (FLP-IN) | 10 | CCL20 (100ng/mL) | |
| | | | LSM059 CMV-TO-bArrestin2-(31-linker)-ADAR2dd(E488Q, T501A) (FLP-IN) | 10 | | |
| | | | LSM041-3 SFFV-mCherry-SGc(stem-1(3xPP7))cSG-EGFP | 200 | | |
| | | CX3CR1-based oLIDAR | LSM103 CMV-TO-CX3CR1-10aa-MCP (FLP-IN) | 10 | CX3CL1 (12nM) | |
| | | | LSM059 CMV-TO-bArrestin2-(31-linker)-ADAR2dd(E488Q, T501A) (FLP-IN) | 10 | | |
| | | | EK0450 SFFV-mCherry-SGc(stem-1(3xMS2))cSG-EGFP | 200 | | |
| S3F | HEK293 | AVPR2-based gLIDAR with SFFV components | APL0030 SFFV-bArrestin2-MCP (FLP-IN) from EK0334/XZ024 | 50/100/200 | AVP (50nM) | 24-well |
| | | | APL0031 SFFV-AVPR2-ADAR2 (FLP-IN) from EK0334/XZ030 | 50/100/200 | | |
| | | | EK0450 SFFV-mCherry-SGc(stem-1(3xMS2))cSG-EGFP | 100 | | |
| | | AVPR2-based gLIDAR with CMV components | XZ024 CMV-TO-bArrestin2-MCP (FLP-IN) | 10 | | |
| | | | XZ030 CMV-TO-AVPR2-ADAR2(DD-E488Q-T501A) (FLP-IN) | 10 | | |
| | | | EK0450 SFFV-mCherry-SGc(stem-1(3xMS2))cSG-EGFP | 100 | | |
| S3G | HEK293 | CCR6-based gLIDAR with SFFV components | APL0030 SFFV-bArrestin2-MCP (FLP-IN) | 50/100/200 | CCL20 (100ng/mL) | 24-well |
| | | | APL0043 SFFV-CCR6-ADAR2 (FLP-IN) | 50/100/200 | | |
| | | | EK0450 SFFV-mCherry-SGc(stem-1(3xMS2))cSG-EGFP | 100 | | |
| | | CCR6-based gLIDAR with SFFV components | XZ024 CMV-TO-bArrestin2-MCP (FLP-IN) | 10 | | |
| | | | XZ032 CCR6-V2-ADAR2dd(E488Q, T501A) | 10 | | |
| | | | EK0450 SFFV-mCherry-SGc(stem-1(3xMS2))cSG-EGFP | 100 | | |
| S3I | HEK293 | B2AR-based gLIDAR, sensor | XZ142 CMV-TO-signal-B2AR-ADAR2dd(DD-E488Q-T501A) (FLP-IN) | 10 | Isoproterenol (100uM) | 24-well |

| | | | | | | |
|---|---|---|---|---|---|---|
| | | with 1 stem loop and 1 stop codon | XZ024 CMV-TO-bArrestin2-MCP (FLP-IN) | 10 | | |
| | | | EK0450 SFFV-mCherry-SGc(stem-1(3xMS2))cSG-EGFP | 200 | | |
| | | B2AR-based gLIDAR, sensor with 1 stem and 2 stop codons | XZ142 CMV-TO-signal-B2AR-ADAR2dd(DD-E488Q-T501A) (FLP-IN) | 10 | | |
| | | | XZ024 CMV-TO-bArrestin2-MCP (FLP-IN) | 10 | | |
| | | | LSM081 SFFV-mCherry-SGc(stem-1(3xMS2)2-stops)cSG-EGFP (FLP-IN) | 200 | | |
| | | B2AR-based gLIDAR, sensor with 2 stems and 1 stop codon each | XZ142 CMV-TO-signal-B2AR-ADAR2dd(DD-E488Q-T501A) (FLP-IN) | 10 | | |
| | | | XZ024 CMV-TO-bArrestin2-MCP (FLP-IN) | 10 | | |
| | | | LSM075 SFFV-mCherry-SGc(stem-1(3xMS2)x2)cSG-EGFP (FLP-IN) | 200 | | |
| S4A | HEK293 | Sensor only | EK0450 SFFV-mCherry-SGc(stem-1(3xMS2))cSG-EGFP | 200 | NA | 24-well |
| | | Sensor and FKBP-TM-ADAR | YH039 PGK-TO-FKBP-tm-GS-XTEN-ADAR2(DD-E488Q-T501A) | 50 | NA | 24-well |
| | | | EK0450 SFFV-mCherry-SGc(stem-1(3xMS2))cSG-EGFP | 200 | | |
| | | Sensor and FRB-TM-MCP module | YH038 PGK-TO-FRB-tm-MCP | 50 | NA | 24-well |
| | | | EK0450 SFFV-mCherry-SGc(stem-1(3xMS2))cSG-EGFP | 200 | | |
| | | Extracellular Binder-based LIDAR | YH038 PGK-TO-FRB-tm-MCP | 50 | AP21967 (100 nM) | 24-well |
| | | | YH039 PGK-TO-FKBP-tm-GS-XTEN-ADAR2(DD-E488Q-T501A) | 50 | | |
| | | | EK0450 SFFV-mCherry-SGc(stem-1(3xMS2))cSG-EGFP | 200 | | |
| | | ADAR-MCP control | LSM101 CMV-TO-ADAR2dd(E488Q, T501A)-MCP (FLP-IN) | 10 | NA | |
| | | | EK0450 SFFV-mCherry-SGc(stem-1(3xMS2))cSG-EGFP | 200 | | |
| S4B | HEK293 | Reporter only | EK0450 SFFV-mCherry-SGc(stem-1(3xMS2))cSG-EGFP | 200 | NA | 24-well |
| | | Reporter and ARRB2-MCP | XZ024 CMV-TO-bArrestin2-MCP (FLP-IN) | 10 | NA | 24-well |
| | | | EK0450 SFFV-mCherry-SGc(stem-1(3xMS2))cSG-EGFP | 200 | | |
| | | Reporter and AVPR2-ADARdd | XZ030 CMV-TO-AVPR2-ADAR2(DD-E488Q-T501A) (FLP-IN) | 10 | NA | 24-well |
| | | | EK0450 SFFV-mCherry-SGc(stem-1(3xMS2))cSG-EGFP | 200 | | |
| | | AVPR2-based gLIDAR | XZ024 CMV-TO-bArrestin2-MCP (FLP-IN) | 10 | AVP (50nM) | 24-well |
| | | | XZ030 CMV-TO-AVPR2-ADAR2(DD-E488Q-T501A) (FLP-IN) | 10 | | |
| | | | EK0450 SFFV-mCherry-SGc(stem-1(3xMS2))cSG-EGFP | 200 | | |
| | | ADAR-MCP control | LSM101 CMV-TO-ADAR2dd(E488Q, T501A)-MCP (FLP-IN) | 10 | NA | |
| | | | EK0450 SFFV-mCherry-SGc(stem-1(3xMS2))cSG-EGFP | 200 | | |
| S4C | | Reporter only | EK0450 SFFV-mCherry-SGc(stem-1(3xMS2))cSG-EGFP | 200 | NA | 24-well |

| | | | | | | |
|---|---|---|---|---|---|---|
| | Colo320 -DM | Reporter and FRB-TM-MCP | YH038 PGK-TO-FRB-tm-MCP | 10 | NA | 24-well |
| | | | EK0450 SFFV-mCherry-SGc(stem-1(3xMS2))cSG-EGFP | 200 | | |
| | | Reporter and FKBP-TM-ADAR | YH039 PGK-TO-FKBP-tm-GS-XTEN-ADAR2(DD-E488Q-T501A) | 10 | NA | 24-well |
| | | | EK0450 SFFV-mCherry-SGc(stem-1(3xMS2))cSG-EGFP | 200 | | |
| | | Extracellular Binder-based LIDAR | YH038 PGK-TO-FRB-tm-MCP | 10 | AP21967 (100 nM) | 24-well |
| | | | YH039 PGK-TO-FKBP-tm-GS-XTEN-ADAR2(DD-E488Q-T501A) | 10 | | |
| | | | EK0450 SFFV-mCherry-SGc(stem-1(3xMS2))cSG-EGFP | 200 | | |
| | | ADAR-MCP control | LSM101 CMV-TO-ADAR2dd(E488Q, T501A)-MCP (FLP-IN) | 10 | NA | |
| | | | EK0450 SFFV-mCherry-SGc(stem-1(3xMS2))cSG-EGFP | 200 | | |
| S4D | Colo320 -DM | Reporter only | EK0450 SFFV-mCherry-SGc(stem-1(3xMS2))cSG-EGFP | 200 | NA | 24-well |
| | | Reporter and ARRB2-MCP | XZ024 CMV-TO-bArrestin2-MCP (FLP-IN) | 10 | NA | 24-well |
| | | | EK0450 SFFV-mCherry-SGc(stem-1(3xMS2))cSG-EGFP | 200 | | |
| | | Reporter and AVPR2-ADARdd | XZ030 CMV-TO-AVPR2-ADAR2(DD-E488Q-T501A) (FLP-IN) | 10 | NA | 24-well |
| | | | EK0450 SFFV-mCherry-SGc(stem-1(3xMS2))cSG-EGFP | 200 | | |
| | | AVPR2-based gLIDAR | XZ024 CMV-TO-bArrestin2-MCP (FLP-IN) | 10 | AVP (50nM) | 24-well |
| | | | XZ030 CMV-TO-AVPR2-ADAR2(DD-E488Q-T501A) (FLP-IN) | 10 | | |
| | | | EK0450 SFFV-mCherry-SGc(stem-1(3xMS2))cSG-EGFP | 200 | | |
| | | ADAR-MCP control | LSM101 CMV-TO-ADAR2dd(E488Q, T501A)-MCP (FLP-IN) | 10 | NA | |
| | | | EK0450 SFFV-mCherry-SGc(stem-1(3xMS2))cSG-EGFP | 200 | | |
| S4E | SNU-475 | Reporter only | EK0450 SFFV-mCherry-SGc(stem-1(3xMS2))cSG-EGFP | 200 | NA | 24-well |
| | | Reporter and ARRB2-MCP | XZ024 CMV-TO-bArrestin2-MCP (FLP-IN) | 10 | NA | 24-well |
| | | | EK0450 SFFV-mCherry-SGc(stem-1(3xMS2))cSG-EGFP | 200 | | |
| | | Reporter and AVPR2-ADARdd | XZ030 CMV-TO-AVPR2-ADAR2(DD-E488Q-T501A) (FLP-IN) | 10 | NA | 24-well |
| | | | EK0450 SFFV-mCherry-SGc(stem-1(3xMS2))cSG-EGFP | 200 | | |
| | | AVPR2-based gLIDAR | XZ024 CMV-TO-bArrestin2-MCP (FLP-IN) | 10 | AVP (50nM) | 24-well |
| | | | XZ030 CMV-TO-AVPR2-ADAR2(DD-E488Q-T501A) (FLP-IN) | 10 | | |
| | | | EK0450 SFFV-mCherry-SGc(stem-1(3xMS2))cSG-EGFP | 200 | | |
| | | ADAR-MCP control | LSM101 CMV-TO-ADAR2dd(E488Q, T501A)-MCP (FLP-IN) | 10 | NA | |
| | | | EK0450 SFFV-mCherry-SGc(stem-1(3xMS2))cSG-EGFP | 200 | | |
| S4F | HEK293 | Reporter only | EK0450 SFFV-mCherry-SGc(stem-1(3xMS2))cSG-EGFP | 200 | NA | 24-well |

| | | | | | | |
|---|---|---|---|---|---|---|
| | | AVPR2-based gLIDAR | XZ024 CMV-TO-bArrestin2-MCP (FLP-IN) | 10 | AVP (50nM) | 24-well |
| | | | XZ030 CMV-TO-AVPR2-ADAR2(DD-E488Q-T501A) (FLP-IN) | 10 | | |
| | | | EK0450 SFFV-mCherry-SGc(stem-1(3xMS2))cSG-EGFP | 200 | | |
| | cLSM12 (derived from HEK293 ) | Reporter only | EK0450 SFFV-mCherry-SGc(stem-1(3xMS2))cSG-EGFP | 200 | NA | 24-well |
| | | AVPR2-based gLIDAR | XZ024 CMV-TO-bArrestin2-MCP (FLP-IN) | 10 | AVP (50nM) | 24-well |
| | | | XZ030 CMV-TO-AVPR2-ADAR2(DD-E488Q-T501A) (FLP-IN) | 10 | | |
| | | | EK0450 SFFV-mCherry-SGc(stem-1(3xMS2))cSG-EGFP | 200 | | |
| S5A | HEK293 | Reporter only | EK0450 SFFV-mCherry-SGc(stem-1(3xMS2))cSG-EGFP | 200 | Na | 24-well |
| | | ADAR-MCP control | LSM101 CMV-TO-ADAR2dd(E488Q, T501A)-MCP (FLP-IN) | 10 | NA | 24-well |
| | | | EK0450 SFFV-mCherry-SGc(stem-1(3xMS2))cSG-EGFP | 200 | | |
| | | AVPR2-based gLIDAR | XZ024 CMV-TO-bArrestin2-MCP (FLP-IN) | 10 | AVP (50nM) | 24-well |
| | | | XZ030 CMV-TO-AVPR2-ADAR2(DD-E488Q-T501A) (FLP-IN) | 10 | | |
| | | | EK0450 SFFV-mCherry-SGc(stem-1(3xMS2))cSG-EGFP | 200 | | |
| | | CX3CR1(noV2)-based gLIDAR | XZ024 CMV-TO-bArrestin2-MCP (FLP-IN) | 10 | CX3CL1 (12nM) | 24-well |
| | | | LSM105 CMV-TO-CX3CR1-(noV2)-ADAR2dd(E488Q, T501A) (FLP-IN) | 10 | | |
| | | | EK0450 SFFV-mCherry-SGc(stem-1(3xMS2))cSG-EGFP | 200 | | |
| | | CCR7(noV2)-based gLIDAR | XZ024 CMV-TO-bArrestin2-MCP (FLP-IN) | 10 | CCL19 (120nM) | 24-well |
| | | | LSM044-3 CMV-TO-CCR7-ADAR2dd(E488Q, T501A) (FLP-IN) | 10 | | |
| | | | EK0450 SFFV-mCherry-SGc(stem-1(3xMS2))cSG-EGFP | 200 | | |
| S6A | HEK293 | AVPR gLIDAR cell line (cLSM02) | Same as S1H-I | Same as S1H-I | AVP (50nM) | 24-well |
| S6B | HEK293 | Stable integration of FKBP-ADAR2dd (cLSM11) | Piggybac Transposase from Gao et al. (2018) | 50 | NA | 24-well |
| | | | LSM0252 (PB)EF1a-FKBP-ADAR2dd-IRESv0-PuroR-P2A-BFP | 450 | | |
| | | Stable integration of FKBP-tm-ADARdd (cLSM10) | Piggybac Transposase from Gao et al. (2018) | 50 | NA | |
| | | | LSM0251 (PB)EF1a-FKBP-tm-ADAR2dd-IRESv0-PuroR-P2A-BFP | 450 | | |
| | | | Piggybac Transposase from Gao et al. (2018) | 50 | NA | |

| | | | | | | |
|---|---|---|---|---|---|---|
| | | Stable integration of CCR6-ADARdd (cLSM09) | LSM0250 (PB)EF1a-CCR6V2-ADAR2dd-IRESv0-PuroR-P2A-BFP | 450 | | |
| | | Stable integration for overexpressing ADAR2 (cLSM12) | Piggybac Transposase from Gao et al. (2018) | 50 | NA | |
| | | | LSM0254 (PB)EF1a-ADAR2-T2A-PuroR-P2A-BFP | 450 | | |
| S8E | HEK293 | CCR6-based gLIDAR with direct hIgG1 output | XZ087 SFFV-mCherry-sensor(tight)-hIgG1-TM | 150 | CCL20 (100ng/mL) | 24-well |
| | | | APL0030 SFFV-bArrestin2-MCP (FLP-IN) | 150 | | |
| | | | APL0043 SFFV-CCR6-ADAR2 (FLP-IN) | 150 | | |
| | | CCR6-based gLIDAR with direct hIgG1 output, positive control reporter | XZ098 SFFV-mCherry-LIDARsensor(W)-hIgG1-TM | 150 | | |
| | | | APL0030 SFFV-bArrestin2-MCP (FLP-IN) | 150 | | |
| | | | APL0043 SFFV-CCR6-ADAR2 (FLP-IN) | 150 | | |
| S10E | cLSM12 /13/14 (derived from HEK293 ) | Reporter without base modifications | IVT mRNA from XZ073 (IVT) CMV-TO-T7-LIDARsensor-hybridUTR3 (0% m1Ψ, 0% m5C) | 500 | NA | 24-well |
| | | Reporter with 100% m5C | IVT mRNA from XZ073 (IVT) CMV-TO-T7-LIDARsensor-hybridUTR3 (0% m1Ψ, 100% m5C) | 500 | | |
| | | Reporter with 100% m1Ψ | IVT mRNA from XZ073 (IVT) CMV-TO-T7-LIDARsensor-hybridUTR3 (100% m1Ψ, 0% m5C) | 500 | | |

**Supplementary Table 3. Overview of cell lines**

| Cell line name | Description | Cargo plasmids | Selection marker | Fluorescence marker |
|---|---|---|---|---|
| XZCP07 | Membrane-bound CCL20 | XZ076 | Puromycin | TagBFP |
| XZCP08 | CCL20 sensing gLIDAR, EGFP output | XZ054, XZ079 | Puromycin, Hygromycin | mCherry |
| XZCP14 | VEGF-sensing eLIDAR, EGFP output | XZ054, XZ337 | Puromycin, Hygromycin | mCherry, TagBFP |
| XZCP17 | IL-2-sensing eLIDAR, EGFP output | XZ054, XZ393 | Puromycin, Hygromycin | mCherry |
| cLSM01 | CCL20 sensing gLIDAR, iCasp9-EGFP output | LSM043_2, XZ079 | Puromycin, Hygromycin | mCherry |
| cLSM02 | AVP sensing gLIDAR, TagBFP output | LSM042, LSM061 | Puromycin, Hygromycin | mCherry |
| cLSM03 | Secreted CCL20 | XZ091 | Puromycin | TagBFP |
| cLSM09 | CCR6-ADAR2dd | LSM250 | Puromycin | TagBFP |
| cLSM10 | FKBP-TM-ADAR2dd | LSM251 | Puromycin | TagBFP |
| cLSM11 | FKBP-ADAR2dd | LSM252 | Puromycin | TagBFP |
| cLSM12 | Overexpression of ADAR2 | LSM254 | Puromycin | TagBFP |
| cLSM13 | ADAR2dd-MCP | LSM253 | Puromycin | TagBFP |
| cLSM14 | ADAR2dd-PCP | LSM258 | Puromycin | TagBFP |
| cJKCC121 | EF1a-DIO-mCherry | JKCC121 | Puromycin | - |
| cNK90 | pTRE3G-EGFP | NK90 | Hygromycin | - |

**Supplementary Table 4.** List of ligands.

| Ligand | Alternative name | Standard Concentration | Source | Catalog No. |
|---|---|---|---|---|
| **Rapalog** | AP21967, A/C heterodimerizer | 100 nM | Takara Bioscience | 635055 |
| **AVP** | [Arg8]-Vasopressin | 50 nM | Sigma-Aldrich | V9879-1MG |
| **IGF-1** | - | 200 ng/mL | PeproTech | 100-11 |
| **VEGF-165** | VEGF-A | 100 ng/mL | PeproTech | 100-20 |
| **IL-2** | - | 1 µg/mL | PeproTech | 200-02 |
| **CCL20** | MIP-3 alpha | 100 ng/mL | PeproTech | 300-29A |
| **CCL19** | MIP-3 beta | 120 nM | Shenandoah Biotechnology | 10787-272 |
| **CX3CL1** | Fractalkine | 115 nM | PeproTech | 10773-804 |
| **Oxytocin** | - | 2 µM | Abcam | ab120186 |
| **Isoprenaline hydrochloride** | Isoproterenol | 100 µM | Abcam | ab120710 |
| **AP1903** | Rimiducid | 4 nM | Fisher Scientific | 50-115-1810 |

**Supplementary Table 5.** Down-sampling fractions for experiment in **Supplementary Fig. 6.**

| Sample | Uniquely aligned reads | fraction | Experiment |
|---|---|---|---|
| AVP(-) | 83989368 | 0.872089501 | Supplementary Fig. 6A |
| AVP(+) | 73246246 | 1 | |
| WT HEK | 126772528 | 0.577776961 | |
| full length ADAR2 | 82966794 | 1 | Supplementary Fig. 6B |
| WT HEK | 126772528 | 0.654454047 | |
| cLIDAR | 114728080 | 0.723160311 | |
| eLIDAR | 119548978 | 0.693998354 | |
| gLIDAR | 118393164 | 0.700773518 | |
